# Roles of Interstitial Albumin in the Subcutaneous Delivery of Therapeutic Peptides

**DOI:** 10.64898/2026.10.14.759110

**Authors:** David Juriga, Anton Norberg, Esther Olaniran Håkansson, Jeroen Wilco van den Heuvel, Johan Gråsjö, Luke R. Odell, Anna Orlova, Per Hansson

**Affiliations:** Pharmaceutical Physical Chemistry, Department of Medicinal Chemistry, Faculty of Pharmacy, Uppsala University, Biomedicinskt Centrum BMC, Husargatan 3, 751 23, Uppsala, Sweden; Department of Medicinal Chemistry, Faculty of Pharmacy, Uppsala University, Biomedicinskt Centrum BMC, Husargatan 3, 751 23, Uppsala, Sweden; Theranostics Group, Department of Medicinal Chemistry, Faculty of Pharmacy, Uppsala University, Biomedicinskt Centrum BMC, Husargatan 3, 751 23, Uppsala, Sweden

**Keywords:** Albumin, Subcutaneous administration, Therapeutic peptide, Extracellular matrix, Albumin binding

## Abstract

Subcutaneous administration of biotherapeutics is increasingly favoured for better safety profile, improved stability, and patient compliance compared with intravenous administration. Despite these advantages, the roles of extracellular matrix (ECM) components and interstitial proteins, particularly albumin, in the mechanisms governing therapeutic pharmacokinetics and bioavailability remain poorly understood. We present a versatile and customizable hydrogel platform resembling native ECM in composition, charge density, and microstructure, enabling systematic investigation of transport mechanisms in the subcutaneous space. Using this platform, we studied peptides with different net charges and albumin affinities. Based on results from confocal microscopy combined with measurements of diffusivities by means of FRAP, we propose a role for albumin beyond that of transport vehicle for drugs. We show that reversible peptide-albumin binding can both increase bioavailability by removing peptides bound to the ECM biopolymers and modulate peptide diffusivity and distribution, as albumin was found to have an affinity for matrix glycosaminoglycans. For cationic peptides, albumin had a major influence on diffusivity and distribution in the matrix, with more pronounced effects over time *in vitro*. These findings, complemented by exploratory *in vivo* observations in a limited cohort, highlight realistic ECM structure and albumin inclusion as key considerations for predictive *in vitro* subcutaneous models.

## Introduction

Subcutaneous (SC) administration of biotherapeutics has become increasingly favoured over intravenous or intramuscular injection due to reduced reliance on medical staff during administration^[1]^, improved patient compliance^[2,3]^, a better safety profile^[4]^, and lower overall cost^[2,5]^. Moreover, SC administration provides a faster route for delivering biotherapeutics such as peptides and proteins compared to oral administration, as it bypasses the gastrointestinal tract, where degradation occurs^[6]^, and avoids poor uptake across the intestinal epithelium^[7,8]^. In addition, SC administration may enable specific delivery strategies, including local delivery, targeting of the lymphatic system^[9,10]^, and resulting in prolonged systemic exposure^[11]^. However, it is not without limitations, such as the restricted injectable volume and pain at the injection site^[12]^. More importantly, predicting pharmacokinetics and bioavailability remains challenging due to limited mechanistic understanding of how therapeutic agents are absorbed into the circulation following injection^[1,5]^. Pronounced physiological and biological differences hinder the translation of *in vivo* animal results to humans^[13–15]^, while *in vitro* models are also facing similar translation problems. Together with the growing number of SC-administered products in clinical use and development^[11,16]^, this highlights a strong need for novel *in vitro* models to elucidate how physicochemical properties of therapeutic agents influence pharmacokinetics^[5,17]^. Among the poorly understood determinants of subcutaneous absorption, the role of interstitial albumin remains particularly underexplored.

A wide range of *in vitro* models has been developed to predict the *in vivo* performance of subcutaneously injected therapeutics, spanning from simple diffusion assays to advanced lab-on-a-chip systems^[13,18,19]^. One of the most frequently used platforms is the SCISSOR system, which employs a cartridge filled with a hyaluronic acid-based hydrogel intended to mimic the extracellular matrix. In addition to measuring transport rates, SCISSOR allows assessment of matrix-drug interactions via turbidity changes in the hyaluronic acid matrix^[20–22]^, and has recently been coupled with in silico approaches to predict the SC bioavailability of monoclonal antibodies^[23]^. Although SCISSOR represents a well-designed experimental platform, its hyaluronic acid matrix lacks sufficient similarity to the native extracellular matrix (ECM) of subcutaneous tissue. While several studies have emphasized the importance of more complex ECM-mimicking matrices^[13,24,25]^, none have captured the full complexity of the subcutaneous ECM, including variations in net charge and charge heterogeneity, the presence of spatially distinct macromolecular domains, or competitive molecular interactions^[26]^. Moreover, existing techniques largely focus on diffusion rate measurements and offer limited opportunity to directly interrogate molecular interactions within the matrix.

Beyond the choice of matrix, the composition of the surrounding medium has emerged as a critical factor for accurately characterizing release profiles and predicting bioavailability^[17,27]^. Torres-Terrán and coworkers developed simulated interstitial fluids based on detailed compositional analyses^[28,29]^ and demonstrated their strong influence on the diffusion of the GLP-1 agonist liraglutide. Similarly, Li and coworkers reported that inclusion of albumin in the BioJect model significantly altered the diffusion of different insulin formulations^[26]^. Although these studies highlight the substantial impact of albumin on therapeutic diffusion in SC *in vitro* models, only a limited number have addressed the underlying mechanisms in detail.

Human serum albumin (HSA) is the most abundant protein in blood, interstitial fluid, and lymph, accounting for approximately 55 %, 68 %, and 65 % of total protein content, respectively. HSA concentrations in interstitial fluid range from 16-28 mg/mL, approximately 20-30 % lower than in blood, giving rise to a unique circulation pattern within the human body^[30]^. Due to its higher concentration in blood, HSA continuously diffuses into the interstitium and predominantly returns to the bloodstream via the lymphatic system^[31]^, although recent studies suggest partial reabsorption directly into the circulation at the venular side of the capillary bed^[32]^. In addition to its diverse physiological roles, HSA contributes to interstitial homeostasis by transporting lipids and metabolic waste products back to the circulation^[33,34]^. Importantly, HSA possesses several binding sites, primarily Sudlow I and II, capable of accommodating a wide range of ligands, from small molecules and lipids to larger peptides^[35,36]^. This property has been exploited in therapeutic design to extend circulation times^[33]^, and in the concept of albumin hitchhiking for compounds administered into the interstitium^[37]^, whereby therapeutic agents associate with HSA and are transported back to the bloodstream via lymphatic drainage, profoundly altering pharmacokinetics and bioavailability. Although albumin hitchhiking has been demonstrated experimentally, no studies have elucidated the detailed mechanisms by which interstitial albumin modulates release kinetics following subcutaneous injections. Despite extensive work on monoclonal antibodies, mechanistic investigations focusing on therapeutic peptides remain absent.

Here, we address this gap by investigating how interstitial albumin regulates the diffusion of low-molecular-weight therapeutic peptides within the extracellular matrix following subcutaneous administration. To this end, we developed a set of ECM-mimicking hydrogel matrices composed of the three major components of subcutaneous tissue: type I collagen, hyaluronic acid, and chondroitin sulfate, allowing systematic modulation of matrix net charge, which varies with obesity and between patients. Five peptides with distinct charge properties and albumin-binding affinities were selected for interrogation. Using confocal microscopy and image analysis, we quantified peptide and albumin partitioning within the matrices, while local diffusion was measured by fluorescence recovery after photobleaching (FRAP). In addition, we developed a gradient diffusion setup enabling measurements within seconds of injection. Finally, we performed exploratory *in vivo* SPECT/CT experiments in a limited cohort of mice to track peptide fate following subcutaneous administration, providing preliminary contextual insights rather than direct validation of the *in vitro* hydrogel mechanism.

Together, this integrated approach highlights key limitations of simplified *in vitro* models and suggests that albumin serves as an active regulator of peptide partitioning and transport within glycosaminoglycan-rich ECM domains.

## Results

### Hydrogel platform with a tunable chemical structure to accurately mimic the ECM composition

To dissect the mechanistic role of interstitial albumin in subcutaneous peptide transport, we developed a chemically defined hydrogel system based on type I collagen (COL), hyaluronic acid (HA), and chondroitin sulfate (CS). The total polymer concentration was selected based on our previously published results and literature data to mimic the mechanical properties and composition of the native adipose tissue ECM^[38,39]^. Varying the relative amounts of these native ECM components allowed independent control over hydrogel charge density while recapitulating different states of the adipose tissue ECM and enabling systematic variation of albumin concentration. This design allowed us to test the effect of ECM negative-charge density, which can vary between patients and with obesity level *in vivo*^[40,41]^. We used HSA solutions at different concentrations, from low to physiological, as loading media to assess how albumin distribution and diffusion within the hydrogels influence the distribution and diffusion of therapeutic peptides with varying net charge and albumin-binding affinity. As a reference medium, we used neutral dextran with a molecular weight similar to HSA, which lacks specific interactions with both the peptides and the hydrogel matrix (Figure 1A). To our knowledge, this novel experimental setup has not been presented previously and provides a unique opportunity to dissect the individual mechanisms governing therapeutic peptide transport after subcutaneous administration.

**Figure 1:**
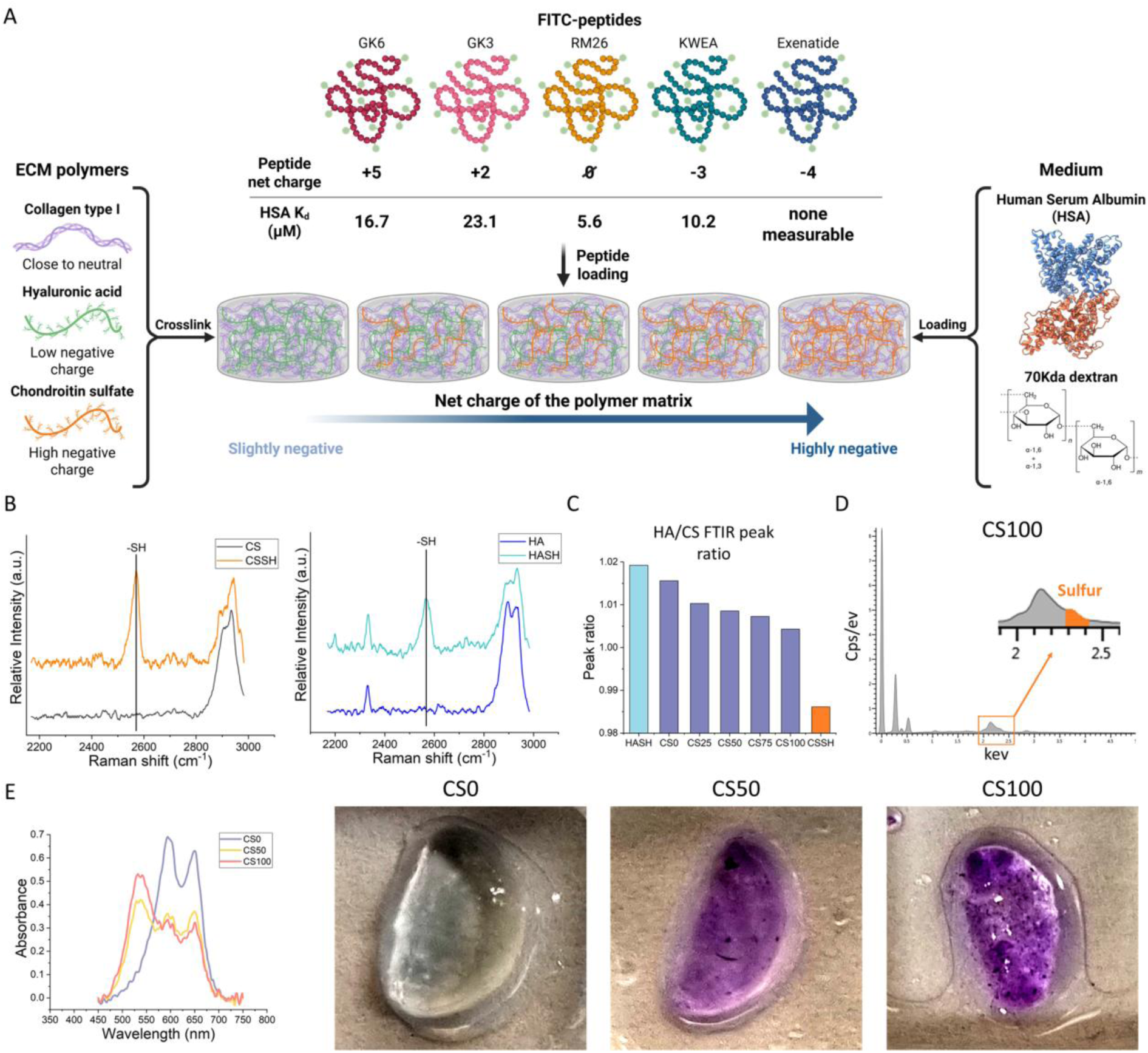
Schematic illustration of the *in vitro* tissue model setup (image was created by *Biorender*) (A); Raman spectra of native and functionalized hyaluronic acid and chondroitin sulfate (B); Ratios of HASH and CSSH specific FTIR peaks across different hydrogel compositions (C); EDS spectrum of the freeze-dried CS100 hydrogel (D); and DMMB staining and corresponding absorbance spectra demonstrating CS content of the hydrogels (E).

FITC-labeled peptides were used for FRAP analysis. Commercially labeled GK6 and GK3 were used as received, while KWEA and Exe were labeled in-house, probably via the N-terminus and RM26 on the PEG linker, synthesized as described in the supplementary information. The amino acid sequences of the peptides and the structure of RM26, with an assumed FITC location, are shown in Figure S1. As labeling slightly alters peptide charge, post-labeling net charges are summarized in Table 1. Successful labeling and the absence of free FITC were confirmed by LC-MS (Figure S2-S4). Although minor amounts of unlabeled peptides remained for KWEA and Exe, this did not affect the microscopy measurements, since they were not included in the measurements due to the absence of a fluorophore.

**Table 1.** Molecular weight, net charge, HSA dissociation constant, and hydrodynamic radii calculated from diffusion data of the different peptides after FITC labeling.

| Peptide | Molecular weight (Da) | Net charge | HSA dissoc. const. ($\mu\text{M}$) | Hydrodynamic radius (nm) |
| --- | --- | --- | --- | --- |
| F-GK6 | 2216 | +5 | 16.7±3.3 | 1.1±0.04 |
| F-GK3 | 2003 | +2 | 23.1±3.9 | 1.1±0.09 |
| F-RM26 | 1662 | 0 | 5.6±1.4 | 0.85±0.07 |
| F-KWEA | 2147 | -3 | 10.2±2.3 | 1.1±0.05 |
| F-Exe | 4575 | -4 | no binding | 1.4±0.04 |

Peptide binding to HSA was further investigated using Microscale Thermophoresis (MST), revealing measurable binding for all FITC-labeled peptides except Exe (Figure S5). The dissociation constants were determined by automated model fitting using the analysis software (Table 1), where lower values represent higher binding affinity. Beyond net charge, FITC influences both the molecular weight and HSA binding affinity of the peptides. Mao and coworkers reported that FITC can bind to two BSA binding pockets, suggesting a similar interaction with HSA^[42]^. Given the small size of the investigated peptides, this interaction significantly increases peptide binding, as shown in Table 1. Since all investigated peptides were FITC-labeled, this mechanism influenced all fluorescence-based experiments. However, MST results suggest that this effect is not equally pronounced across all peptides, yielding a panel of peptides with distinct HSA affinities, enabling the systematic investigation of this parameter.

To introduce cross-links among COL, HA, and CS, the polymers were functionalized using protocols published by Rodler and coworkers^[38]^. According to these protocols, approximately 40 % of the lysine residues in COL were functionalized with methacrylate groups^[38]^, and a fraction of the carboxyl groups in HA and CS were functionalized with thiol groups. To confirm the presence and stability of the thiol groups, Raman spectroscopy was performed (Figures S6 and 1B). The appearance of thiol-specific peaks (C-S stretching at 660 cm^−1^ and S-H stretching at 2570 cm^−1^) in the CSSH and HASH spectra confirms successful thiolation of the carboxyl groups, while the absence of peaks associated with disulfide bonds (~500 cm^−1^) indicates thiol stability. The extent of polymer modification was further confirmed by NMR spectroscopy (Figure S7), which indicated approximately 38 % and 80 % modification for HASH and CSSH, respectively, in good agreement with DTNB assay results. Based on a calibration curve (Figure S8), the DTNB assay revealed 45-48 % and 70-73 % modification of the carboxyl groups in HASH and CSSH, respectively. Together, these results demonstrate controlled polymer functionalization, which is essential for the reproducible synthesis of the hydrogel matrices. Multiple orthogonal methods, including FTIR, SEM-EDS, and the DMMB assay, confirmed controlled incorporation of CS into the hydrogels (Figure 1C-E). Using FTIR measurements, we first identified characteristic peaks associated with HASH (1032 cm^−1^) and CSSH (1021 cm^−1^) that do not overlap with COL peaks (Figure S9). Quantification of the HA-to-CS peak height ratio revealed a decreasing HA/CS ratio with increasing CS content in the hydrogel synthesis (Figure 1C).

SEM-EDS analysis of freeze-dried hydrogels further supported these findings as secondary evidence, as a sulfur-associated peak^[43]^ at 2.3 keV, next to the peak of the gold coating at 2.1 keV, appeared in CS50 and became more pronounced in CS100 (Figures S10 and 1D). Consistently, an increasing sulfur density was observed in the EDS layered SEM images (Figure S10). The addition of DMMB^[44]^ induces a characteristic absorbance shift from 595 nm to 525 nm, detected in the spectra of CS50 and CS100 but not in CS0, accompanied by the appearance of a deep purple color in the hydrogel (Figure 1E), supporting the previous findings. This level of compositional control was essential to decouple the effects of ECM charge density from peptide-albumin interactions, allowing systematic dissection of how albumin modulates peptide diffusion and retention within a physiologically relevant extracellular matrix. In the following sections, we use this platform to quantify the impact of albumin-peptide interactions on transport behavior under defined ECM conditions.

### ECM-mimicking hydrogels recapitulate native ECM heterogeneity, leading to heterogeneous molecular distribution

To assess whether the hydrogel system recapitulates the physical heterogeneity of the native extracellular matrix^[45]^, we characterized its microstructure and molecular transport properties at the microscale (Figure 2).

**Figure 2:**
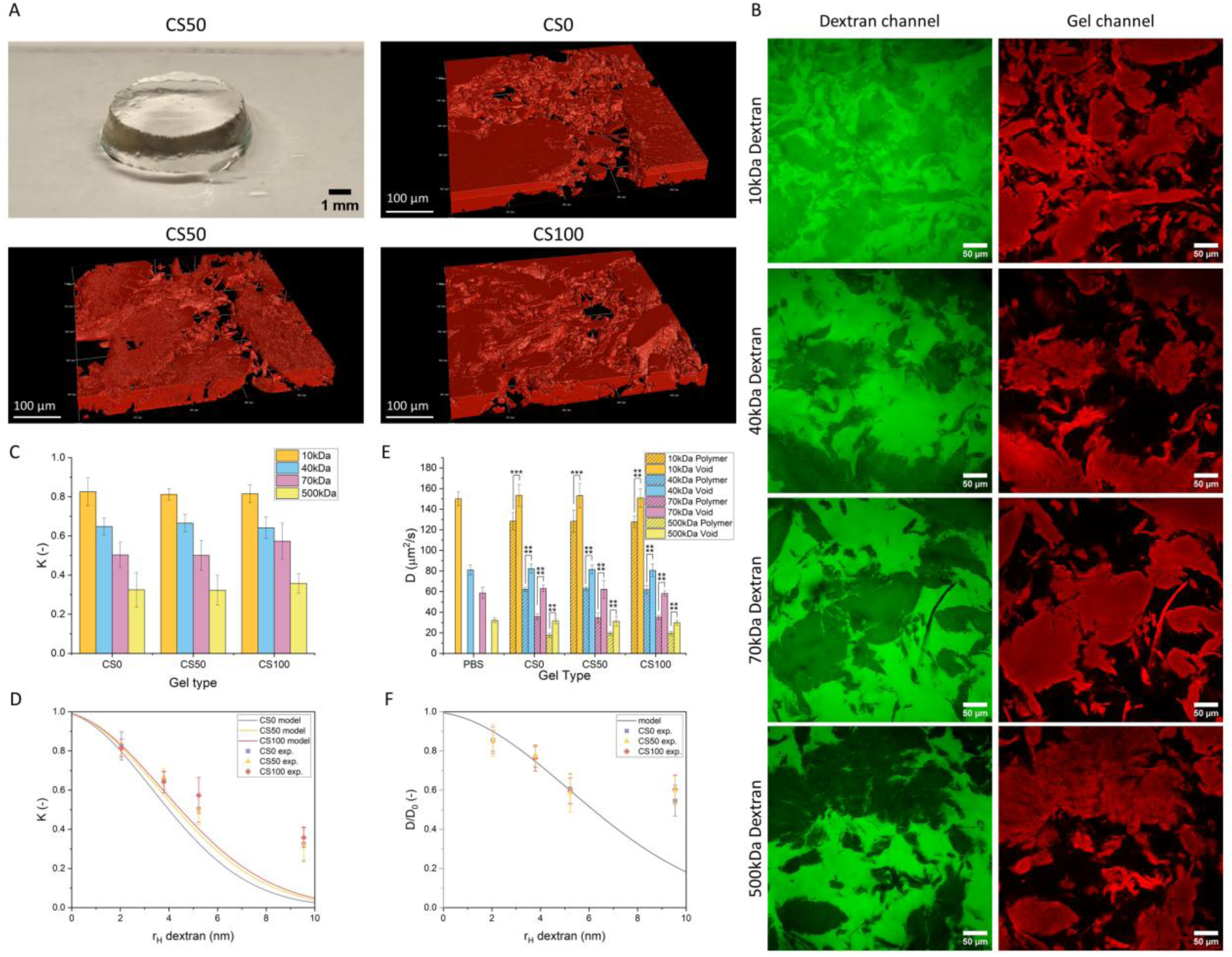
Macroscopic photograph of the CS50 hydrogel and three-dimensional confocal microscopy images of hydrogels with different compositions (A); confocal microscopy images of FITC-dextran probes with different molecular weights loaded into the CS50 hydrogel (B); partition coefficients (C); and diffusion coefficients (E) of FITC-dextran probes measured in the different hydrogels. Statistical significance was marked as p < 0.05 (*), p < 0.01 (**), p < 0.001 (***), and p < 0.0001 (****), N_g_=3 N_m_=4/2/2 for partitioning and N_m_=3/3/2 for diffusion coefficient. Solid curves in Figures D and F: Theoretical predictions according to Eq. (1) with r_f_=0.6 nm, Φ=0.006, and *φ* = 0.54 (CS0), 0.62 (CS50), and 0.66 (CS100) (D), and Eq. (3) with *ξ*= 13.1 nm and r_f_=0.6 nm (F).

Although the macroscopic appearance of the hydrogel is homogeneous and transparent (Figure 2A, upper left), the confocal microscopy revealed a heterogeneous microstructure consisting of polymer-rich domains separated by black void spaces (Figure 2A), consistent with previous observations^[24]^ and SEM imaging (Figure S11). Although the hydrogels exhibited similar microstructures, image analysis showed a progressive increase in the volume fraction of polymer-rich domains with increasing CS content (Figure S11 and text after).

Size-dependent exclusion was assessed using FITC-dextrans of increasing molecular weight (Figure 2B and Figure S12). Larger dextrans preferentially localized to the voids, whereas smaller dextrans distributed more evenly between domains (Figure 2B), indicating increasing steric exclusion from the polymer-rich domains with increasing molecular size. The partition coefficients determined by image analysis further supported this observation (Figure 2C). While the K value for 10 kDa dextran was approximately 0.8 in all hydrogels, it decreased to ~0.4 or lower for 500 kDa dextran, indicating enhanced exclusion from polymer-rich domains with increasing molecular weight. Statistical analysis showed no significant differences in K for a given dextran across different hydrogels, whereas most dextran sizes exhibited significant differences within individual hydrogel types (Table S1). The solid curves in Figure 2D show partition coefficients predicted by the Ogston theory (Eq. (1)), calculated as described in the Materials and Methods section. The good agreement with the experimental values for r_h_≤4 nm implies that the Ogston theory, combined with the volume fraction of COL, CS, and HA and an effective fiber radius obtained as a weighted average of the individual polymer radii, accounts for the partitioning of dextran probes with r_h_ values comparable to those of HSA and the peptides used in this work. According to the Ogston theory^[46]^, the average diameter of the largest sphere that fits into the space between the network strands is equal to 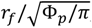, where Φ*_p_* is the polymer volume fraction. For the present hydrogels, this diameter is 9.8, 10, and 11 nm for CS0, CS50, and CS100, respectively, which can be interpreted as measures of the mesh size.

In addition to determining partition coefficients, we performed FRAP experiments to measure the diffusion coefficient of the dextran probes separately in the voids and in the polymer-rich domains (Figure 2E). The rFRAP model fitting proved to be highly accurate, demonstrating the reliability of the measurements (Figure S13). The diffusion coefficients of the different dextran probes showed statistically significant differences between PBS solution and those within the polymer-rich domains of the hydrogels (Table S2). FRAP analysis revealed that diffusion coefficients in the voids were indistinguishable from those measured in PBS, indicating that these regions behave as a polymer-free solution. In contrast, diffusion within polymer-rich domains was significantly reduced for all dextran sizes and was independent of hydrogel composition, consistent with steric hindrance by the polymer network, indicating that ECM charge modulation does not affect bulk diffusivity of neutral molecules at the length scale probed here. We determined the correlation length *ξ* in the hydrogel networks by fitting the Amsden diffusion model^[47,48]^ to the data in Figure 2F with *ξ* as a fitting parameter, omitting the data points for the dextran with the highest Mw. The resulting values obtained from the non-linear least-squares fit are 13.4±0.7, 13.1±0.7, and 13.1±0.7 nm for CS0, CS50, and CS100, respectively. For clarity, only the fit for CS100 is shown in Figure 2F. In the model, *ξ* equals the diameter of the largest spherical space that fits between openings between polymer chains, i.e., the Ogston model mesh size. It is rewarding that the estimates are in close agreement with the value calculated from Eq. (1).

The heterogeneous architecture of the hydrogels provides a structural basis for differential transport of macromolecules and is expected to strongly influence the distribution and mobility of albumin and albumin-bound peptides within the matrix.

### Albumin shows a more homogeneous distribution than dextran, indicating attraction to ECM polymers

After characterizing the hydrogels using dextran probes of different molecular weights, we performed the same confocal microscopy measurements with human serum albumin (HSA) at three concentrations (1, 7.5, and 25 mg/mL), as HSA concentration can also vary *in vivo* in the interstitial space^[49]^. As a reference, neutral 70 kDa dextran (similar in molecular weight to HSA) was used to distinguish size-dependent effects from specific interactions that could influence its distribution and diffusion. In addition, rheological and swelling measurements were performed to assess whether differences in these properties of the hydrogels could influence diffusion behavior (Figure 3).

**Figure 3:**
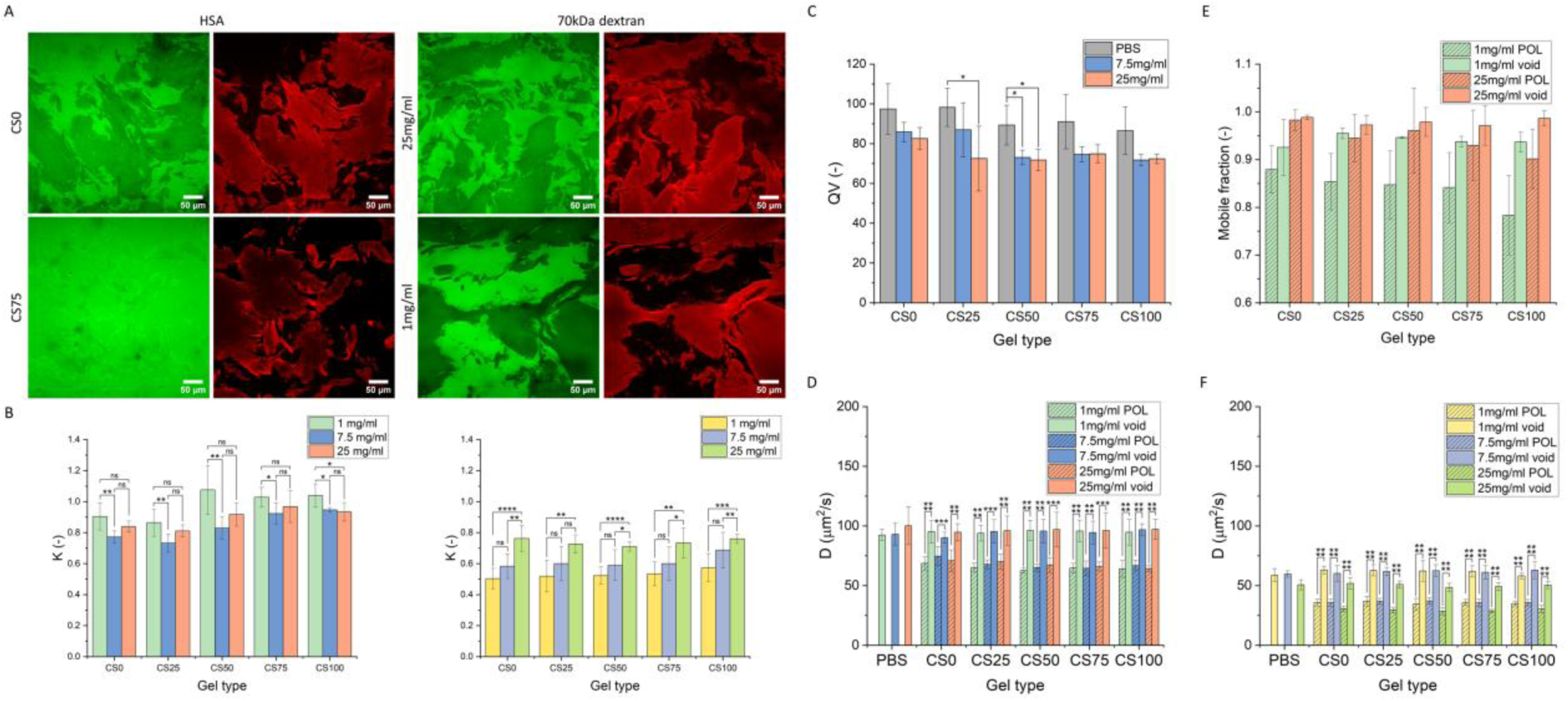
Confocal microscopy images of two ECM-mimicking hydrogels (CS0 and CS75) loaded with HSA (25 mg/mL) and of the CS50 hydrogel loaded with different concentrations of a 70 kDa dextran probe (A); partition coefficients of HSA and the dextran probe at different concentrations (B); volumetric degree of swelling (Q_V_) of the different hydrogels at different HSA concentrations (C) (N=4); Diffusion coefficients (D) and mobile fraction (E) of HSA (N_g_=3 N_m_=3//3/2) in different hydrogels; Diffusion coefficient of 70 kDa dextran (N_g_=3 N_m_=3/3/2) (F) in the different hydrogels. Statistical significance was marked as p < 0.05 (*), p < 0.01 (**), p < 0.001 (***), and p < 0.0001 (****)

In contrast to dextran of comparable molecular weight, the composition-dependent partitioning of HSA within the hydrogels indicates electrostatic attraction of HSA to the ECM polymers. Confocal images illustrate the effect of hydrogel charge density on HSA distribution between polymer-rich domains and voids (Figure 3A). At low negative charge (CS0), HSA was more strongly excluded from polymer-rich domains, as indicated by lower fluorescence intensity, than at high negative charge (CS75). Similar trends were observed at all HSA concentrations tested (Figure S14). By increasing the CS content of the hydrogel matrices, the partition coefficient (K) increased at all HSA concentrations studied and reached or exceeded 1 in CS100 hydrogels (Figure 3B), with statistically significant differences observed at 7.5 and 25 mg/mL HSA (Table S3). In addition, K values slightly decreased with increasing HSA concentration across all hydrogel compositions, indicating a concentration-dependent modulation of HSA partitioning within the ECM-mimicking matrices.

In contrast to HSA, the dextran probe of similar molecular weight exhibited significantly lower K values at all concentrations across all hydrogel compositions (Figure 3A, B). Moreover, hydrogel composition did not influence dextran partitioning, whereas increasing dextran concentration resulted in higher K values as a result of solute-solute steric interactions^[50]^. Due to the rigid structure of HSA and the repelling effect between HSA molecules, similar behavior was not observed. The nearly uniform partitioning of HSA between polymer-rich domains and voids shows that there are attractive interactions between HSA and the glycosaminoglycans within the polymer-rich domains, compensating for the size exclusion effect, suggesting a role of albumin-ECM interactions in governing the fate of subcutaneously administered biotherapeutics.

Besides distribution, we also investigated the effect of HSA and the dextran probe on the viscoelastic and swelling properties of the hydrogels. Neither HSA nor dextran significantly altered the viscoelastic properties of the hydrogels, indicating that differences in molecular transport are not caused by bulk mechanical changes (Figures S15, S16, and 3C). Furthermore, these results prove that hydrogels are stable at different HSA concentrations up to 7 days, similar to results published in a previous paper^[51]^. Although HSA did not influence G* (Figure S15B), it slightly decreased (by 15-18 %) the volumetric degree of swelling of all the hydrogels (Figure 3C), while the dextran probe did not influence these properties (Figure S15C and S15D).

FRAP experiments revealed that HSA diffusion in the voids was indistinguishable from diffusion in PBS, whereas diffusion within polymer-rich domains was significantly reduced (Figure 3D). On the other hand, the mobile fraction of HSA slightly decreased at 1 mg/ml concentration with increasing CS content, indicating slightly different interactions between HSA and CS compared to HSA and HA (Figure 3E). Dextran diffusion was also significantly hindered in the polymer-rich domains (Figure 3F), and it also showed a slight concentration dependency due to the previously mentioned solute-solute interaction, which decreases the hydrodynamic radius of the dextran molecules at high concentration, which effect was statistically significant in most hydrogel compositions (Table S4). The rFRAP fitting showed high accuracy for both HSA and dextran (Figures S17 and S18). Furthermore, HSA diffused faster than 70 kDa dextran under all conditions, consistent with its smaller hydrodynamic radius.

Together, these results demonstrate that albumin, in addition to its established role as an active transporter of other species, exhibits matrix-dependent partitioning and mobility, which is expected to modulate peptide transport in the subcutaneous space.

### Interstitial albumin modulates charge-driven peptide partitioning in ECM-mimicking hydrogels

After characterizing the distribution of HSA and dextran probes, we next investigated the effect of HSA concentration (1, 7.5, and 25 mg/mL) on peptide distribution. Due to molecular interactions, such as electrostatic interactions and HSA binding, certain therapeutic peptides can remain longer in the subcutaneous tissue than others, thereby influencing pharmacokinetics and bioavailability^[52]^. To capture this behavior as an additional dimension, measurements were performed at multiple time points after loading the peptides into the hydrogels (Figure 4).

**Figure 4:**
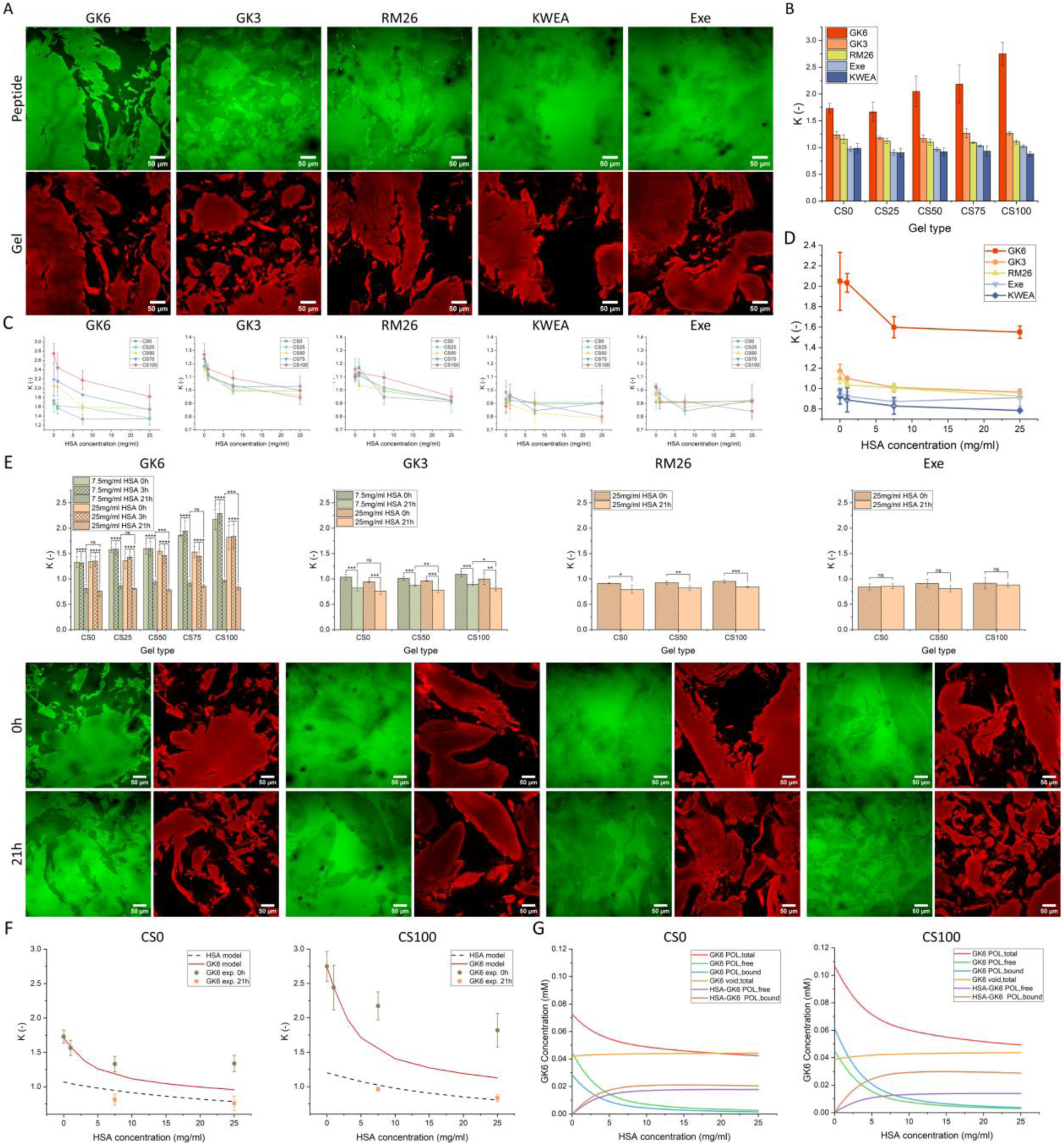
Distribution of different peptides in the CS50 hydrogel in the absence of HSA (A); partition coefficients of the different peptides across all hydrogel types without HSA (B); HSA dependence of the partition coefficients of the different peptides across all hydrogel types (C); comparison of HSA-dependent changes in partition coefficients in the CS50 hydrogel (D); and time-dependent changes in partition coefficients of the different peptides at 7.5 mg/mL and 25 mg/mL HSA, together with representative confocal microscopy images illustrating time-dependent redistribution in CS50 hydrogels at 25 mg/mL HSA (E). Calculated and experimentally determined partition coefficients of GK6 and HSA in CS0 and CS100 hydrogels as a function of HSA concentration (F). Calculated peptide concentrations in the different states for the systems shown in Figure F, respectively (G). Statistical significance was marked as p < 0.05 (*), p < 0.01 (**), p < 0.001 (***), and p < 0.0001 (****)

The confocal images in Figure 4A demonstrate the effect of the peptides’ charge on their distribution within the CS50 hydrogel. In the absence of albumin, peptide partitioning was dominated by electrostatic interactions: the highly cationic GK6 preferentially localized to polymer-rich domains, whereas less positively charged or anionic peptides showed progressively weaker or no enrichment. Increasing ECM charge density further enhanced GK6 retention, with the K value increasing from 1.7 to 2.7, while exerting minimal effects on the distribution of the other peptides (Figure 4A-B and Table S5).

In Figure 4C, we demonstrate the effect of HSA on the K of all peptides across the different hydrogels. HSA concentration was highly influential for positively charged peptides and for the neutral RM26, with statistically significant effects observed (Table S6-S8), whereas no notable effect was detected for KWEA or Exe (Table S9-S10). Confocal images show increased fluorescence intensity in the voids for GK6, GK3, and RM26 at 25 mg/mL HSA compared to PBS (Figure S19-S21), while this difference was less pronounced for KWEA (Figure S22) and for Exe (Figure S23). Among the peptides, GK6 exhibited the strongest response to HSA, which also depended on hydrogel charge density: 25 mg/mL HSA reduced K by 23 % in CS0 and by 35 % in CS100, indicating that albumin progressively attenuates charge-driven partitioning of GK6. Consistently, 25 mg/mL HSA lowered the K value of GK3 to approximately 1 and reduced the K of RM26 to below 1, irrespective of hydrogel charge density, while its influence remained negligible for KWEA and Exe (Figure 4C and Figure S24-S25). Comparison of the different peptides in CS50 is shown in Figure 4D, with corresponding analyses for the other hydrogels presented in Figure S25. These comparisons highlight the pronounced differences between GK6 and the other peptides, indicating the combined importance of peptide charge and polymer-rich domain charge density in governing peptide partitioning within the hydrogel matrices.

Notably, albumin induced a pronounced time-dependent redistribution of cationic peptides. The partitioning of the highly cationic GK6 showed a pronounced time-dependent response to HSA. While the K value of GK6 remained unchanged after 3 h, it had decreased significantly in all hydrogel types after 21 h of incubation at 34 °C, both at 7.5 and 25 mg/mL HSA. Initially, GK6 strongly partitioned into polymer-rich domains (K = 1.25-2.25, depending on hydrogel composition); however, after 21 h, K values converged to ~0.75 in all hydrogels, indicating preferential localization in the voids. This redistribution is clearly visible in the confocal microscopy images, which show a shift from dominant fluorescence intensity in polymer domains immediately after loading to higher intensity in the voids after prolonged incubation (Figure 4E).

The extent of this redistribution depended on both HSA concentration and ECM charge density. After 21 h, statistically significant differences between 7.5 and 25 mg/mL HSA were observed in hydrogels with higher CS content, and 25 mg/mL HSA eliminated differences in GK6 partitioning among hydrogel compositions, while 7.5 mg/mL significantly reduced them (Table S11). These results indicate that albumin progressively overrides the effect of ECM charge density on GK6 retention.

A similar but less pronounced time-dependent decrease in K was observed for the moderately cationic GK3 at both HSA concentrations, with K values converging to ~0.75 after 21 h across all hydrogel types (Figure 4E). Confocal images confirmed a redistribution pattern comparable to GK6, although statistical analysis revealed no significant differences between hydrogels at 21 h (Table S12). RM26 showed a smaller absolute shift in K (from ~0.95 to ~0.8), while no time-dependent redistribution was observed for the negatively charged exenatide. Confocal microscopy images provided under the corresponding graphs in Figure 4E also support these trends.

We hypothesized that HSA promotes a more uniform partitioning of cationic peptides between polymer-rich domains and voids by providing binding sites that compete with peptide binding to negatively charged polyelectrolytes in the ECM. To test this, we provide a thermodynamic model as a theoretical framework to investigate the proposed mechanism by which albumin, through peptide and ECM binding, promotes a more uniform distribution of GK6 between polymer-rich domains and voids in the ECM. The detailed description of the molecular thermodynamic calculations is presented in the Supplementary Information. Briefly, we first determined the binding constants of peptide and HSA to the glycosaminoglycans by fitting the model to experimental data for peptide-only and HSA-only systems. This approach incorporates system-specific effects related to chemical differences between carboxylate and sulfate groups directly into the binding constants. In the model, the HSA-CS complex involves a CS segment carrying 9 negative charges binding to a 4.5 nm positively charged patch on HSA with a net charge of +5.2, based on the model by Ly et al^[53]^. Likewise, one GK6 molecule (+5) binds to a 5.1 nm CS segment with a charge −10. Applying the same approach to HSA-HA and GK6-HA yielded lower binding constants than for CS, consistent with the experimental results, although the model underestimated the binding strength in both cases (Figures 4 F and G and in Table 2). Importantly, to describe peptide-HSA interactions, we used the experimentally determined peptide-HSA dissociation constant (*K_D_*) in all calculations, avoiding model-dependent estimates of this key parameter.

**Table 2.** Experimental partition coefficient (*K*), calculated binding constant (*K_b_*), and binding free energy (Δ*G*^0^) for HSA and GK6 in CS0 and CS100 gels at 307 K.

| Gel type | HSA |  |  | GK6 |  |  |
| --- | --- | --- | --- | --- | --- | --- |
| | $K_{HSA}$ | $K_{b,HSA}/10^3$ | $-\Delta G_{HSA}^0/kT$ | $K_{pep}$ | $K_{b,pep}/10^3$ | $-\Delta G_{pep}^0/kT$ |
| CS0 | 0.87<sup>a</sup> | 32 | 10.4 | 1.7 | 24 | 10.1 |
| CS100 | 1.1<sup>b</sup> | 58 | 11.0 | 2.7 | 54 | 10.9 |

a) Average value for three HSA concentrations: 1 mg/mL (0.954), 7.5 mg/mL (0.795), 25 mg/mL (0.859)
b) Valid for 1 mg/mL HSA

Figure 4F shows that HSA remains fairly uniformly distributed at all compositions. In agreement with experiments, the calculated partition coefficient of GK6 between polymer-rich domains and voids decreased with increasing HSA concentration in both CS0 and CS100 hydrogels. Moreover, the model captured the observed non-linear dependence on HSA concentration. However, it clearly overestimated the effect of HSA on peptide distribution after short incubation. The reason for this will be discussed later.

As a result, the peptide distribution in the hydrogel becomes more like that of HSA, consistent with the experimental observations after 21 h of incubation at 34 °C (Figure 4F). This is further supported by Figure 4G, which shows that the calculated peptide concentration in the polymer-rich domain decreases with increasing HSA concentration and approaches the concentration in the void. The concentration in the void remains nearly constant because the combined volume of the void and the external liquid phase is much larger than that of the polymer-rich domain. The calculated concentrations of peptide not bound to HSA show that, in the absence of HSA, the fraction bound to the polyelectrolyte exceeds the free fraction. As HSA concentration increases from 0 to 25 mg/mL, both fractions decrease substantially. At the same time, the ratio of polyelectrolyte-bound to free peptide decreases because HSA reduces the number of available binding sites on the polyelectrolyte. Furthermore, according to the model, the peptide remaining in the polymer-rich domain becomes progressively more associated with HSA the higher the HSA concentration. As a result, a very small fraction remains freely dispersed or directly bound to the polyelectrolyte at the highest HSA concentration.

To clarify the underlying mechanism, we performed a control calculation for the same system assuming no peptide binding to HSA (*K_D_*→∞) and no HSA binding to the ECM domains separately. Under the first condition, the peptide partition coefficient was nearly independent of HSA concentration, while without HSA-CS interaction, GK6 partitioning shows stronger dependence on HSA concentration (Figure S26). Model calculations indicate that peptide redistribution is driven by direct peptide–HSA binding rather than competitive displacement of the peptide by HSA at polyelectrolyte sites, though HSA–ECM polymer interactions contribute equally to the overall process. The results showed (Figure S26A) that HSA’s partition coefficient was well below unity (K ≈ 0.3) at all HSA concentrations, and that GK6’s partition coefficient decreased from 2.8 to 0.5 as the HSA concentration increased from 0 to 25 mg/mL. The result supports the idea that HSA modulates the partitioning and mobility of HSA-binding peptides not only by binding the peptides but also by binding to the glycosaminoglycans in the ECM. In summary, the results presented in this section demonstrate that peptide charge is the dominant determinant of initial ECM retention, while interstitial albumin dynamically counteracts electrostatic trapping over time. For highly cationic peptides such as GK6, albumin progressively neutralizes the influence of ECM charge density, driving redistribution from polymer-rich domains, revealing a dynamic, competitive mechanism potentially of great importance for subcutaneous transport of albumin-binding peptides. According to the thermodynamic model calculations, HSA induces the redistribution of GK6 mainly by providing binding sites for the peptide that tend to outcompete the peptide’s binding to the polyelectrolytes, but HSA affinity to the ECM polymers also has a significant role in the process. The mechanism is simple and conceptually clear, but the calculations are only in semi-quantitative agreement with the experiments, and therefore do not prove the mechanism directly.

### Albumin overrides charge-driven control of peptide diffusion

We have already demonstrated that HSA can strongly influence peptide distribution in ECM-mimicking hydrogels by modulating interactions between peptides and ECM macromolecules and through direct peptide binding. Since such molecular interactions can alter peptide diffusion properties, we next investigated peptide diffusion within the hydrogels using FRAP and a gradient diffusion measurement setup (Figure 5), also including 70 kDa dextran as a reference macromolecule.

**Figure 5:**
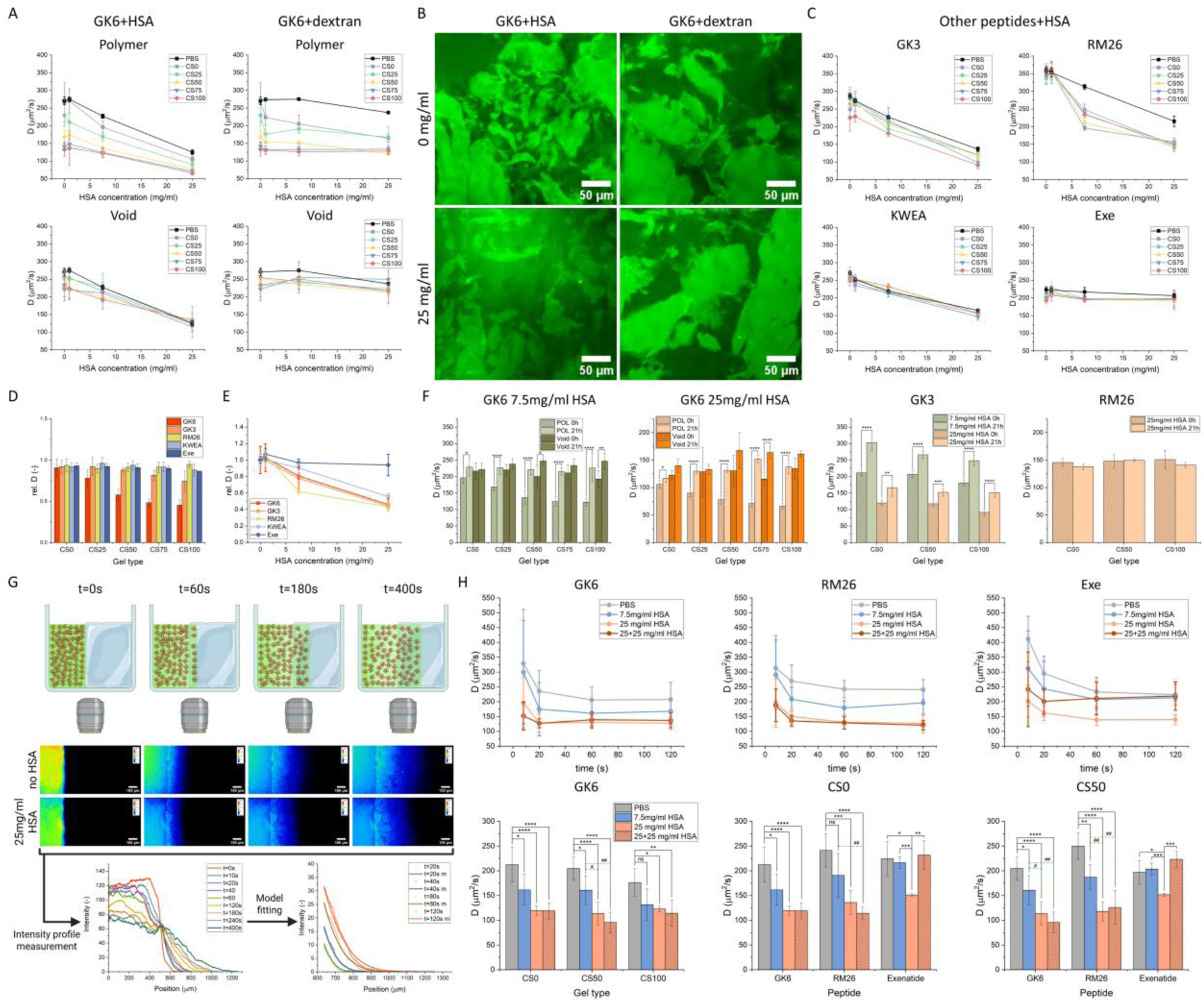
Diffusion coefficients of GK6 determined by FRAP separately in polymer-rich domains and voids at different concentrations of HSA and reference dextran probe (A); confocal microscopy images of the CS50 hydrogel loaded with GK6 acquired at different HSA and dextran concentrations (B); HSA-dependent diffusion coefficients of the other peptides (C); relative diffusion coefficients normalized to PBS for different peptides measured in all hydrogel types in the absence of HSA (D); relative diffusion coefficients normalized to PBS for different peptides measured in the CS50 hydrogel at different HSA concentrations (E); time-dependent effects of HSA on diffusion coefficients (F); experimental setup for gradient diffusion measurements (Image was created by *Biorender*) (G); and comparison of diffusion coefficients determined by gradient diffusion measurements for different peptides at varying HSA concentrations across different hydrogels (H) (N_g_=2 N_m_=3). Statistical significance was marked as p < 0.05 (*,#), p < 0.01 (**,##), p < 0.001 (***,###), and p < 0.0001 (****,####)

The diffusion of GK6 was strongly influenced by both hydrogel composition and HSA concentration, while dextran concentration exerted only a minor effect (Figure 5A). The rFRAP model fits the experimental data accurately (Figure S27). The D of GK6 within polymer-rich domains showed a statistically significant decrease with increasing CS content in the hydrogels, independent of HSA concentration (Figure 5A, upper left graph; Table S13). Increasing HSA concentration further reduced D and had a pronounced effect on GK6 diffusion across all hydrogel types (Figure 5A; Table S13). Although dextran concentration also affected GK6 diffusion, primarily in hydrogels with low negative charge density (CS0 and CS25) and in PBS, this effect was significantly weaker than that observed for HSA (Figure 5A, upper right graph; Table S14). Dextran concentration did not influence GK6 diffusion in the hydrogel voids (Figure 5A, lower right graph), whereas HSA concentration produced a distinct effect (Figure 5A, lower left graph). At low HSA concentration, modest differences in diffusion were observed among hydrogel types; however, these differences were eliminated at physiological HSA concentration. While no significant differences were detected between diffusion coefficients measured in voids and polymer-rich domains for CS0 and CS25 at low HSA concentration, higher HSA levels, particularly in more highly charged hydrogels, resulted in significantly lower diffusion coefficients in polymer-rich domains than in the voids (Table S15), indicating a combined role of albumin binding and electrostatic interactions in regulating GK6 diffusion. Consistent with these findings, confocal microscopy images show that HSA alters the spatial distribution of GK6, whereas dextran does not, supporting the mechanistic connection between albumin binding, ECM interactions, and peptide diffusion (Figure 5B).

Affinity to HSA proved to be the most relevant factor affecting the diffusion of the other peptides (Figure 5C). Diffusion of RM26, which exhibits the strongest HSA binding among the investigated peptides, was already drastically reduced at 7.5 mg/mL HSA and exhibited significantly lower values in the hydrogels than in PBS (Table S16). GK3 and KWEA, which show moderate HSA binding, exhibited a more gradual decrease with statistical significance (Table S17-18), but the difference between the values measured in hydrogels did not statistically differ from the values measured in PBS. In contrast, diffusion of Exe was not influenced by HSA concentration. Hydrogel composition only slightly affected the diffusion of GK3 and RM26 and reached statistical significance only at specific HSA concentrations (Table S16-S18). Replacing HSA with the dextran control medium affected the diffusion of GK3, RM26, and KWEA only in PBS but not in hydrogels, while dextran significantly reduced Exe diffusion in all media, due to its bigger size (Figure S28 and Table S19). The rFRAP model fits were also accurate for these peptides (Figure S29-S32).

To compare the different peptides, we examined the effects of hydrogel composition (Figure 5D) and HSA concentration (Figure 5E) on the relative change in diffusion coefficients (D) compared to those measured in PBS. As shown in Figure 5D, hydrogel composition influenced diffusion only for positively charged peptides, in a manner dependent on peptide charge density, whereas no effect was observed for neutral or negatively charged peptides. By comparing diffusion coefficients measured in PBS with those measured in hydrogels at each HSA concentration, increasing HSA concentration resulted in significant differences for GK3, RM26, and KWEA, while no effect was observed for Exe (Table S20). In contrast, HSA binding emerged as an equally important, or even dominant, determinant of peptide diffusion in hydrogels (Figure 5E), with RM26 showing a markedly stronger response than the other peptides at 7.5 and 25 mg/mL HSA, consistent with its higher albumin-binding affinity.

Interestingly, HSA exerted a pronounced time-dependent effect on the diffusion coefficient of GK6 within polymer-rich domains, significantly increasing D after 21 h of incubation at 34 °C (Figure 5F). After long incubation, HSA largely equalized diffusion coefficients across different hydrogel compositions (Table S21), eliminating the effect of the charge density in the polymer-rich domains. The diffusivities in the voids also increased slightly after 21 h at both 7.5 and 25mg/ml HSA concentrations; however, these changes were not statistically significant in most cases. A similar time-dependent increase in diffusion was observed for GK3 after 21 h of incubation, independent of hydrogel composition, whereas RM26 showed no statistically significant change at 25 mg/mL HSA (Table S21). Together with the partitioning results, these findings demonstrate that HSA dynamically modulates electrostatic interactions within ECM-mimicking hydrogels and enhances peptide diffusion over time by progressively releasing peptides from ECM trapping.

Besides FRAP measurements, which provide diffusion coefficients without a concentration gradient present, we designed an experimental setup to model diffusion immediately after injection (Figure 5G). As shown in Figure 5H, the determined diffusion coefficients depended on the time points selected for model fitting (upper graphs). In most cases, higher diffusion coefficients were obtained from the first 8 and 20 s of the measurements compared to later time points, probably due to a higher concentration gradient as well as partial drying of the hydrogel surface due to medium change. These differences between early and later stages were markedly reduced at higher HSA concentrations, accompanied by a decrease in standard deviation. Based on these observations, four time points between 20 and 120 s were selected for model fitting to enable robust comparison of diffusion coefficients across different peptides and HSA concentrations (Figure 5H, lower graphs).

For GK6, trends like those observed by FRAP were detected. Increasing HSA concentration resulted in progressively lower D. Moreover, physiological HSA concentration, as well as the addition of HSA to the peptide solution, eliminated the effect of hydrogel charge density on diffusion. These observations are consistent with the FRAP results, where HSA similarly reduced the influence of negative charge density within polymer-rich domains. The addition of 25 mg/mL HSA to the GK6 solution did not significantly affect D compared to hydrogels preincubated with 25 mg/mL HSA alone. Comparing the diffusion behavior of GK6, RM26, and Exe (Figure 5H, lower right graphs) revealed trends largely consistent with FRAP measurements but highlighted one key difference related to Exe. RM26 diffusion showed strong dependence on HSA concentration, while hydrogel composition was not a relevant factor. The addition of HSA to the RM26 solution did not result in a statistically significant difference compared to conditions where only the hydrogel was incubated with HSA. In contrast, low HSA concentration did not affect Exe diffusion, whereas at 25 mg/mL HSA, diffusion coefficients were significantly reduced. Notably, adding HSA directly to the Exe solution before injection abolished this reduction and restored diffusion coefficients to levels measured in the absence of HSA. The latter observation indicates that the effect is caused by transport of albumin out of the gel, lowering the diffusive flow of Exe inside the gel. Although the exact mechanism has not been investigated, since the effect is absent for GK6 and RM26, we hypothesize an electrodynamic interaction related to Exe’s negative charge and/or reflecting the different partitioning of the peptides in the gel.

Together, the diffusion measurements demonstrate that albumin-peptide binding and albumin-ECM interactions are the primary determinants of peptide mobility in the subcutaneous matrix. While electrostatic interactions initially govern diffusion of cationic peptides, physiological albumin concentrations progressively suppress charge-driven differences and dynamically enhance peptide transport over time. These findings show that interstitial albumin could function as an active regulator of subcutaneous peptide diffusion under both equilibrium and injection-relevant conditions.

### Exploratory In Vivo Evaluation of Subcutaneous Clearance and Spreading Kinetics

Besides the *in vitro* experiments, we also investigated the *in vivo* performance of GK6, ABD-RM26′, and Exe exploratorily on a limited cohort. GK6 was selected due to its high positive charge to test the relevance of electrostatic interactions, RM26 due to its high HSA binding affinity *in vitro*, and Exe as a control peptide since it lacks both properties. GK6 and Exe were labeled with ^12^⁵I using indirect lysine labeling for SPECT/CT measurements, whereas the RM26 structure was modified with a DOTA chelator to enable radiolabeling with ^111^In and with a small-molecule albumin-binding domain (ABD) to ensure high albumin binding *in vivo* (ABD-RM26’), without altering the charge and size significantly (Supplementary information methods section and Figure S33). Based on previously published data, the 4-(p-iodophenyl)butanoic acid group can significantly increase albumin binding of therapeutic peptides *in vivo*^[54–58]^, while the 600 Da increase in molecular weight does not significantly alter core transport behavior (Figure 2F), ensuring that the structures remain comparable. Furthermore, SPR measurements were carried out to compare *in vitro* albumin binding of ABD-RM26’ with the peptide used in the project and semaglutide, a strong albumin-binding therapeutic peptide^[59]^. The results showed that ABD-RM26’ K_d_ is two orders of magnitude smaller than semaglutide and three orders of magnitude smaller than the peptides investigated in the project (Figure S34 and Table S22). Spatial spreading was quantified by measuring the distribution volume, while peptide clearance was assessed by monitoring activity at the injection site for up to 2 h. After 2 h of continuous imaging under anesthesia, the animals were allowed to move freely for 1 h and imaged once more to assess the effect of motion on peptide distribution and clearance (Figure 6A).

**Figure 6:**
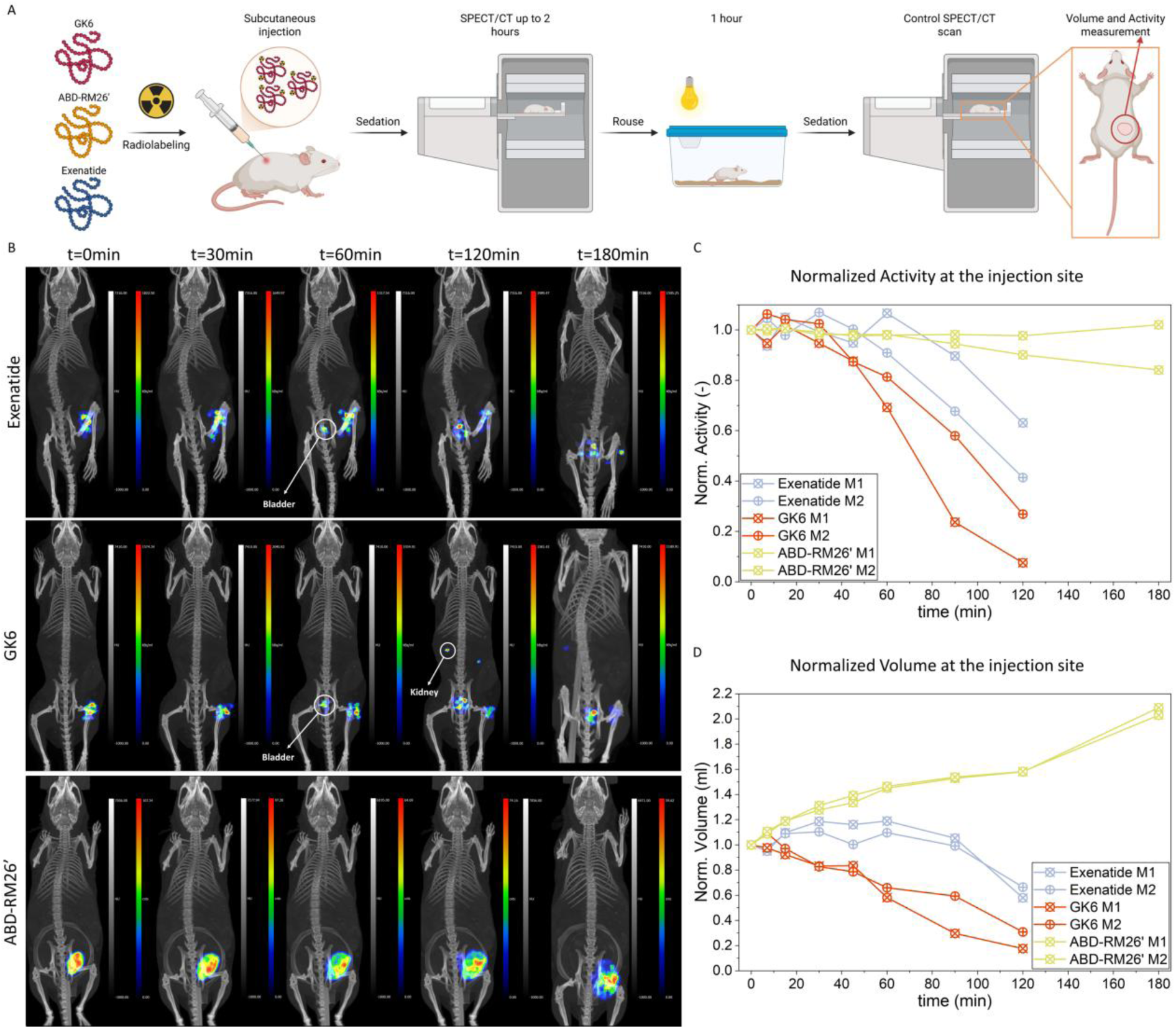
Schematic illustration of the *in vivo* experimental setup (image was created by *Biorender*) (A); SPECT/CT images of mice acquired at different time points after subcutaneous injection (B); time-dependent changes in activity at the injection site for the three peptides, normalized to the initial time point (C); and time-dependent changes in the volume at the injection site for the three peptides, normalized to the initial time point (D).

SPECT/CT imaging revealed pronounced differences in subcutaneous retention and clearance among the three peptides (Figure 6B). Exe showed rapid clearance from the injection site, with activity decreasing after 60 min and dropping sharply after 90 min (Figure 6C, blue, and Figure S35). The peptide spreads within the subcutaneous tissue, reaching maximal volume within 30 min, after which further expansion was limited by systemic clearance, as evidenced by early accumulation in the bladder (Figures 6B, S35, and Video S1). After 1 h of free movement, Exe was almost completely cleared from the injection site.

GK6, despite its strong ECM interactions observed *in vitro*, exhibited rapid clearance *in vivo*, similar to Exe. In contrast to Exe, GK6 remained confined to a more localized volume throughout the experiment (Figure 6B), but its amount declined faster, with near-complete disappearance from the injection site within 120 min in one animal, but complete clearance after the wake period in both (Figures 6C, S35, and Video S2). Activity accumulation in the kidneys at several time points suggests rapid renal clearance, consistent with its high positive charge^[60]^. However, renal clearance only affects peptides already present in the blood circulation and does not influence clearance from the subcutaneous tissue. In comparison, if the animals were awake for 60 min after injection, the clearance from the injection site was significantly faster for both peptides, probably due to constant motion of the animals (Figure S36).

In contrast, ABD-RM26′, which exhibits strong albumin binding, showed markedly prolonged retention at the injection site. Activity remained essentially constant throughout the experiment, even after 1 h of unrestricted movement (Figure 6C and S35). At the same time, ABD-RM26′ displayed greater spatial spreading within the subcutaneous tissue compared to the other peptides, as visible in the SPECT/CT images and reflected by the increasing distribution volume (Figures 6B, 6D, and Video S3), which is consistent with the high albumin flow in the interstitial space^[61]^.

These exploratory *in vivo* observations align qualitatively with the *in vitro* findings, showing that the albumin-binding domain exhibited prolonged subcutaneous residence and enhanced spatial distribution at the injection site relative to other peptides with low or non-specific binding. While structural differences across the series, such as net charge, chemical modifications, radiolabeling tags, and the inclusion of the ABD, preclude isolating albumin binding as the sole causal driver, the observed trends suggest that engineered albumin affinity plays an important role in extending local subcutaneous tissue residence. Furthermore, in this exploratory cohort, positive peptide charge alone appeared insufficient to prevent rapid clearance in the absence of strong albumin interaction. Overall, these preliminary animal data provide supportive context for the mechanistic behaviors observed in the hydrogel model, though expanded *in vivo* studies with larger cohorts and strictly matched molecular controls are required to fully isolate individual chemical drivers and establish definitive causality.

## Discussion

### Relevance of ECM-mimicking hydrogel

Subcutaneous injection is increasingly used for antibodies, proteins, and peptides, yet the prediction of pharmacokinetics and bioavailability remains challenging because mechanistic models of transport in subcutaneous tissue are still limited^[1,2,5,6]^. Existing *in vitro* systems mainly measure transport through simplified ECM models and therefore provide limited insight into the tissue properties that regulate biotherapeutic transport^[5,11,17]^. To address this gap, we developed a chemically crosslinked hydrogel platform based on the three major ECM components of subcutaneous tissue: collagen type I, hyaluronic acid, and chondroitin sulfate.

Using thiol-Michael addition chemistry^[38]^, we generated hydrogels with controlled viscoelastic properties and tunable negative charge density by varying the HA/CS ratio while keeping collagen content constant, resulting in a composition and mechanical properties similar to native ECM. In contrast to our previous collagen-HA network^[38,51]^, incorporation of CS provided robust control over charge density, which is particularly relevant because ECM charge may vary between individuals and with obesity^[62,63]^, influencing peptide transport. Although CS is present at lower levels than HA in adipose tissue^[64]^, its higher charge density suggests a stronger contribution to interactions with cationic species^[65]^.

Although the hydrogels appeared macroscopically homogeneous (Figure 2A), fluorescent labeling of collagen showed polymer-rich domains interspersed with voids identical to PBS, both ranging from approximately 10 to several hundred µm in thickness, similar to that reported in a previous paper^[51]^. This heterogeneity had a clear functional consequence: fluorescent dextran probes were increasingly excluded from the polymer-rich domains with increasing molecular weight. For probes with hydrodynamic radii up to 4 nm, the partitioning data were well described by the Ogston model, yielding a mesh size of 10-11 nm in the collagen-rich domains, while diffusion-based Amsden’s theory gave a mesh size of 13 nm. Although direct mesh-size measurements for adipose ECM are not available, pore-theory analyses of diffusion data have reported pore diameters of 12-20 nm^[66]^. Considering that our estimated mesh sizes apply only to the polymer-rich domains, which occupy 54, 62, and 66 % of the gel volume for CS0, CS50, and CS100, respectively, the agreement is reasonably good. This supports the conclusion that the hydrogel ECM model reproduces the net obstruction effects to diffusion relevant for adipose ECM.

The heterogeneous structure may also resemble the native subcutaneous matrix more closely than homogeneous gel models^[67]^. Histological studies have shown that adipose ECM is divided into at least two structurally distinct regions: the reinforced basement membrane in the narrow space between adipocytes (~2 µm) and the interlobular septa between fat lobules (~10 µm)^[68]^. These regions differ in composition, particularly in collagen content, and are therefore expected to differ in average mesh size.

This structural heterogeneity is consistent with the proposal that interstitial transport resembles size-exclusion chromatography, where smaller solutes access a larger fraction of tissue volume than larger ones^[66,67]^. The observation that even large nanoparticles can reach the lymphatics after interstitial injection has been interpreted as evidence for open channels of 50-100 nm^[66,69]^, which may form when interlobular septa are mechanically separated by injection pressure, as shown by X-ray tomography^[70]^. Together, these similarities suggest that our hydrogel model can mimic drug transport and drug-matrix interactions in adipose tissue across multiple length scales^[71]^.

In the present work, we investigated the behavior of small peptides in the ECM-mimicking hydrogels, with particular focus on electrostatic interactions with network components and on the effect of HSA, potentially important determinants of drug absorption rate and bioavailability after subcutaneous administration^[72,73]^.

### Albumin’s effect on peptide partitioning

In the absence of albumin, peptide partitioning was governed mainly by peptide charge and the charge density of the polymer-rich domains; the positively charged GK6 was strongly enriched in the polymer-rich domains, and this enrichment increased with increasing CS content, whereas negatively charged peptides were excluded. Albumin markedly altered this behavior, reducing the concentration of positively charged peptides in the polymer-rich domains, while showing little or no effect on the negatively charged peptides. At physiological HSA concentration, the differences in partitioning between peptides were largely eliminated after only a short incubation period. After longer incubation, albumin fully redistributed the positively charged peptides, resulting in partition coefficients below unity even for GK6, irrespective of hydrogel composition. Thus, HSA was able to weaken or outcompete the electrostatic attraction between positively charged peptides and the glycosaminoglycans (GAGs) in the polymer-rich domains even in hydrogels with high negative charge density, thereby decreasing peptide trapping and allowing more peptide to redistribute into the voids.

Albumin is the most abundant protein in interstitial fluid and has mainly been discussed in subcutaneous delivery through the concept of albumin hitchhiking^[37]^. Previous *in vitro* studies emphasized peptide-albumin binding^[26,28,74]^, whereas albumin interactions with the extracellular matrix and their effect on diffusion remained largely unexplored. Here, we found that HSA was distributed almost equally between polymer-rich domains and voids despite its relatively large size and overall negative charge. The partition coefficient increased slightly with increasing negative charge density of the hydrogel, reaching values above 1 at the highest CS content. In contrast, dextran of similar molecular weight was strongly excluded from the polymer-rich domains (Figure 3). HSA diffusion in the polymer-rich domains was reduced to ~67% of that in the voids, whereas dextran diffusion decreased to ~50%. In addition, the mobile fraction of HSA decreased with increasing CS concentration, particularly at 1 mg/mL HSA, indicating stronger interactions of HSA with CS-rich matrices than with HA-rich matrices, consistent with our thermodynamic modelling. This behavior is notable because HSA carries a net charge of approximately −15^[53,75]^. Based on gel composition, we estimated that the concentration of fixed positive charges from collagen in the polymer-rich domain was 1.4 mM, whereas the total concentration of fixed negative charges from CS and HA was 6-14 mM. The network should therefore generate a negative electrostatic potential relative to PBS, which, together with steric obstruction, would be expected to reduce HSA partitioning into the polymer-rich domains. The observation of partition coefficients close to unity, therefore, indicates attractive interactions between HSA and the network chains that overcompensate for these unfavorable effects. This conclusion is supported also by thermodynamic model calculations showing that, by cancelling the HSA-polyelectrolyte binding constant HSA is effectively redistributed to the voids (Fig. S26A). The affinity between albumin and negatively charged polysaccharides is known to depend on ionic strength, pH, and polymer charge density^[76]^. Seyrek and Dubin proposed that this attraction is primarily electrostatic^[76]^, and Yu et al. showed that sodium polyacrylate binds to a specific positively charged site on HSA^[53]^.

In this work, we used molecular thermodynamic modelling to test whether binding of CS and HA to a net positively charged surface groove on HSA accounts for its experimentally observed partitioning. After validating that the model reproduced the HSA-polyacrylate binding constant (Supplementary Information), we adapted it to glycosaminoglycans (GAGs) by determining the net charge of the CS-accessible site required to match HSA’s partition coefficient in the CS100 system. The resulting net charge (+5.2) was only slightly smaller than that determined for polyacrylate (+6), with the binding CS segment carrying 9 negative charges. Overcharging of the site by the polyelectrolyte, as observed with polyacrylate, aligns with the conclusion of Yu et al. that the entropy gain from the release of bound counterions is a primary driving force behind the interaction^[53]^. Applying identical site properties to HSA-HA binding slightly underestimated HSA’s partition coefficient (Table S22). A similar approach was used to define the binding geometry of GK6-CS complexes, which likewise slightly underestimated the partition coefficient when applied to GK6-HA. These discrepancies likely reflect that carboxylate and sulfate groups were not distinguished in terms of ionic character and hydration. To avoid overfitting, parameters determined for CS were applied to HA in both cases.

Thermodynamic modelling provided binding constants for HSA and GK6 binding to GAGs in the polymer-rich domain. Incorporating these alongside the HSA-GK6 dissociation constant into the thermodynamic framework enabled theoretical investigation of how HSA and GK6 affect each other’s partitioning when both are present in the system (Figures 4F and 4G). Notably, the model captured the shift toward a more even distribution of GK6 between polymer-rich domains and voids as HSA concentration increases. These calculations support our hypothesis that the peptide’s affinity for HSA is sufficient to transfer it from GAGs to HSA, thereby adopting an HSA-like distribution. Importantly, the partition coefficient of a control peptide with no HSA affinity, but otherwise identical to GK6 in the model, was only marginally affected by HSA. This allowed us, within the model framework, to rule out the competing mechanism where HSA displaces peptides by simply occupying polyelectrolyte binding sites.

While these theoretical results do not definitively prove the proposed mechanism, given the semi-quantitative agreement between model and experiment, direct experimental control is limited because highly cationic peptides inevitably bind electrostatically to negatively charged HSA. Nevertheless, species concentrations in the polymer-rich domain support the proposed mechanism. At a total polyelectrolyte charge concentration of 14 mM, partition coefficients indicate that GK6 and HSA concentrations should not exceed 0.14 mM and 0.3 mM, respectively. Assuming each peptide and HSA molecule occupies a segment corresponding to 10 polyelectrolyte charges, GK6 occupies 10 % and HSA 21 % of the polyelectrolyte, leaving at least two-thirds unoccupied (accounting for unbound fractions of both molecules). Thus, the calculations show peptide and HSA need not compete for binding sites. Further supporting peptide transfer from GAGs to HSA, the peptide diffusion coefficient decreases with increasing HSA concentration, converging at the highest concentration to the value measured for HSA in the CS100 polymer-rich domain.

As shown in Figure 4F, the thermodynamic model overestimates HSA’s ability to redistribute GK6 from polymer-rich domains to the voids. The origin of this discrepancy remains unclear. Test calculations showed that increasing the *K_D_* value improved model agreement, though no single value provided consistent fits across the full HSA concentration range (not shown). This discrepancy may arise because the experimental *K_D_* value does not apply to peptide binding to polyelectrolyte-bound HSA, or because GK6 binding enhances HSA affinity for CS/HA, driving a larger fraction of the peptide into polymer-rich domains than predicted. Furthermore, time-dependent partitioning observed after 21 h suggests that short-term data reflects systems temporally trapped in metastable states. Discrepancies thus arises because calculated *K* values represent thermodynamic equilibrium. Notably, peptide values measured at 21 h closely align with calculated HSA values, as expected if a dominant fraction of the peptide is HSA-bound. If the 21-h data represent the final equilibrium state, model agreement improves, albeit remaining semi-quantitative, as calculated values exceed experimental ones. Potential mechanisms underlying these time-dependent effects are discussed below.

### Albumin’s effect on peptide diffusion

Albumin binding emerged as the main determinant of peptide diffusion, whereas the charge density of the polymer-rich domains affected diffusion mainly for the highly cationic GK6. In the absence of HSA, GK6 diffusion decreased with increasing negative charge density of the hydrogels, consistent with strong electrostatic interactions with the matrix. Increasing HSA concentration progressively reduced this effect, indicating that albumin weakens charge-driven retention of GK6 in the polymer-rich domains but, at the same time, binding to albumin reduced the measured diffusion coefficients. By contrast, hydrogel composition had little or no effect on the diffusion of the other peptides. GK3 showed only a minor reduction in diffusion in the hydrogels relative to PBS, whereas KWEA and Exe showed no clear hydrogel-dependent differences.

A distinct pattern was observed for RM26. Although RM26 interacted only weakly with the hydrogel matrix itself, its diffusion in the hydrogels was strongly reduced at 7.5 and 25 mg/mL HSA. This difference compared to GK6 is mechanistically informative. For GK6, diffusion reflected both matrix charge and albumin concentration, whereas for RM26, the dominant effect appears to arise from favorable peptide-albumin binding, which hinders the diffusion of the peptide in the dense polymer-rich domains, combined with the affinity of albumin for the glycosaminoglycan-rich polymer domains. Thus, even in the absence of strong direct peptide-matrix interactions, albumin-mediated coupling to the matrix can substantially hinder peptide mobility. In contrast, Exe, which showed no measurable albumin binding and was distributed uniformly in the hydrogels, clearly displayed no dependence of diffusion on either hydrogel composition or albumin concentration.

After 21 h of incubation, the GK6 diffusion coefficient in the polymer-rich domains increased to nearly the same level as in the voids. A similar but weaker tendency was observed for GK3, whereas RM26 showed no time-dependent change. For GK6, this behavior is difficult to reconcile with a simple picture in which HSA is associated with the GAGs in the polymer-rich domains, because under such conditions, very little peptide would be expected to remain freely mobile there. One possible explanation is that, during incubation, the peptide slowly relocates to a site on HSA that weakens HSA binding to the matrix, thereby promoting redistribution of both HSA and peptide toward the voids.

As a complementary experiment, and to better mimic injection conditions, we also followed concentration gradient-driven diffusion of GK6, RM26, and Exe. These measurements reproduced the main FRAP trends with respect to HSA concentration, supporting the previous findings.

### In vivo – in vitro relationship

Exploratory *in vivo experiment*s were carried out by measuring the spatial spreading and residence time of the same three peptides after subcutaneous injection in NMRI mice using SPECT/CT. The goal was not to establish direct quantitative translation or isolate single chemical variables, but rather to identify qualitative patterns that could provide supportive context for the mechanisms proposed in our *in vitro* hydrogel model. To make RM26 more comparable to the *in vitro* conditions, its albumin binding was increased by introducing a previously developed small albumin-binding domain (ABD-RM26’)^[54–58]^, while avoiding significant changes in size, charge, and polarity.

In these pilot imaging scans, GK6 exhibited rapid local clearance from the subcutaneous site despite its strong electrostatic affinity for matrix components *in vitro*, and the occupied volume decreased continuously over time. This trend is qualitatively consistent with our hypothesis that interstitial albumin may dynamically modulate the association of cationic peptides with negatively charged ECM elements. While electrostatic albumin binding reduces GK6’s *in vitro* diffusivity, the dynamic nature of this interaction in a perfused subcutaneous tissue bed could hypothetically facilitate local movement and clearance toward vascular or lymphatic capillary pathways. In comparison, Exe showed a longer residence time but still rapid clearance and only limited spreading during the first hour. These qualitative variations across constructs may reflect compounding physical differences, including effectively larger size (including electrostatic double-layer interactions), by the repelling interaction with the negatively charged wall of the blood capillaries, and by the lack of detectable albumin binding. Conversely, ABD-RM26′ remained largely at the injection site even after 3 h but still displayed greater spreading than Exe. These preliminary observations align with the concept that strong albumin binding can promote movement within the tissue connected to interstitial albumin flow^[49]^, while at the same time limiting clearance from the subcutaneous ECM, due to hindrance of diffusion into blood capillaries^[37]^. The proposed mechanism may help explain how highly cationic species can escape subcutaneous tissue and reach the circulation^[14]^, contributing to design future biotherapeutics. Given the exploratory cohort size (N=2) and concurrent variations in peptide structure, net charge, ABD modification, and radiolabeling chemistry across constructs, these *in vivo* observations cannot isolate albumin binding as a single causal determinant of pharmacokinetics. Nevertheless, these qualitative trends offer useful preliminary context for the mechanistic behaviors observed in our ECM-mimicking hydrogels. Further systematically matched *in vivo* studies with larger cohorts will be required to rigorously isolate individual structural drivers and validate the proposed transport mechanisms.

While our work highlights a novel role for albumin in regulating peptide partitioning and transport within glycosaminoglycan-rich ECM domains, a concept with broader implications for future biotherapeutic development, several limitations can be addressed to refine the *in vitro* model and better mimic the *in vivo* environment. A key limitation is that fluorescence labeling alters the physicochemical properties and albumin affinity of the investigated peptides, which may affect quantitative measurements. However, this issue could be mitigated by selecting fluorophores that do not disrupt albumin binding, an effect that is also likely to become less pronounced with increasing molecular weight of the therapeutic agent. Additionally, the structural complexity of the ECM-mimicking hydrogels could be enhanced to better reflect native ECM composition by incorporating collagen fibers, immune or lymphatic components, and other matrix components such as fibrin or elastin. Furthermore, mimicking lymphatic and capillary drainage would represent a valuable addition for evaluating tissue escape mechanisms. Finally, while observations from our exploratory *in vivo* study align with this proposed mechanism, further *in vivo* studies utilizing larger cohorts and more comprehensive experimental setups, such as unified radiolabelling, remain essential for full validation. However, due to possible species-dependent anatomical differences, application of human-derived in vitro tissue models can also be a key segment to investigate the proposed mechanisms in detail.

## Conclusion

Taken together, our results suggest that albumin may regulate subcutaneous transport through a dual mechanism: by binding therapeutic compounds and, as a novel concept, by interacting with glycosaminoglycans, whereby the distribution and diffusion capacity, particularly of positively charged peptides, within the ECM are also altered. This may weaken electrostatic trapping of positively charged peptides and promote their clearance from the injection site, leading to rapid distribution in the whole body. A similar effect on transport across negatively charged capillary surfaces is also conceivable. Conversely, strong peptide-albumin binding may prolong local residence by limiting passage across the tightly packed capillary wall, while still allowing spreading with albumin flow through the interstitium. We believe that these factors are highly contributing to the prolonged release of compounds designed to bind to albumin, such as semaglutide or liraglutide, after subcutaneous administration. A more fundamental understanding of this dual role of interstitial albumin may therefore open new possibilities for rationally tuning pharmacokinetics and improving the bioavailability of subcutaneously administered biotherapeutics through molecular and formulation design. Although this work relies on fluorescence-based techniques, and fluorescent labeling may alter albumin and/or ECM binding as well as the physicochemical properties of biotherapeutics, the mechanistic insights obtained here remain highly relevant. Their translation, together with similar biomimetic analytical platforms, into industrial workflows could accelerate the optimization of already marketed subcutaneous therapeutics by enabling structure-guided exploitation of albumin-dependent transport mechanisms. Moreover, our work highlights that careful design and microfabrication of hydrogel matrices can improve the performance and reliability of *in vitro* models by more closely reproducing tissue microstructure, thereby making them more powerful tools for drug development across different tissue types.

## Materials and methods

### Materials

All chemicals were purchased from standard commercial suppliers and used as received unless otherwise stated. Detailed information on reagents is provided in the Supplementary Information file, while details of instruments and software are provided in the Methods section.

### Functionalization of the extracellular matrix polymers and preparation of ECM-mimicking hydrogels

ECM-mimicking hydrogels were prepared by photocrosslinking methacryloyl-functionalized type I collagen (MAC) with thiolated glycosaminoglycans. Type I collagen was methacryloylated as described previously by Rodler et al.^[38]^. Thiolation of hyaluronic acid (HA) and chondroitin sulfate (CS) followed the same protocol with the following modifications: HA (500 mg) was dissolved in 120 mL Milli-Q water (rather than 60 mL) to avoid gelation upon addition of 1-ethyl-3-(3-dimethylaminopropyl)carbodiimide (EDC); CS thiolation was extended from 40 min to 2 h. All subsequent steps were performed as described.

For hydrogel formation, MAC was dissolved in Milli-Q water at 1 w/w% and stored at 4 °C until fully dissolved (4–5 days). Thiolated HA (HASH) and thiolated CS (CSSH) were separately dissolved in nitrogen-flushed phosphate-buffered saline (PBS) at 1.5 w/w% immediately prior to use. The photoinitiator lithium phenyl-2,4,6-trimethylbenzoylphosphinate (LAP) was prepared in nitrogen-flushed PBS at 1.5 w/w%.

MAC, HASH, and CSSH solutions were combined with LAP and PBS to achieve the desired final polymer concentrations. The precursor solution was homogenized by brief vortexing and repeated pipetting to remove air bubbles. Defined volumes of the precursor solution (see experimental sections) were transferred to custom molds and crosslinked with a UVP CL-1000 crosslinker (Analytik Jena) for 5 min at a distance of 3 cm from the UV source with an energy of 150 mJ/cm^2^.

After crosslinking, hydrogels were equilibrated in air for 30 min, sealed overnight, and subsequently washed in PBS with two buffer exchanges over 48 h before use. The final theoretical MAC content was 0.7 w/w%; the mass ratio of MAC to HASH + CSSH was 5:1. Volumes used for 1 g gel synthesis are listed in Table M1.

**Table M1:** Amounts of the polymer solutions, LAP solution, and PBS used to prepare 1 g of gel.

| Gel type | 1.5w/w/ MAC solution (mg) | 1w/w% HASH solution (mg) | 1w/w% CSSH solution (mg) | 1.5w/w% LAP solution (mg) | PBS (mg) |
| --- | --- | --- | --- | --- | --- |
| CS0 | 700 | 93,3 | 0 | 70 | 136,7 |
| CS25 | 700 | 70 | 23,3 | 70 | 136,7 |
| CS50 | 700 | 46,7 | 46,7 | 70 | 136,7 |
| CS75 | 700 | 23,3 | 70 | 70 | 136,7 |
| CS100 | 700 | 0 | 93,3 | 70 | 136,7 |

### NMR spectroscopy

^1^H NMR spectra were recorded on a 400 MHz JEOL JNM-ECP FT NMR spectrometer (JEOL, Oxford NMR Instruments, UK). Samples were dissolved in 99.9% D₂O at concentrations of 4–5 mg mL^−1^. Spectra were acquired with at least 64 scans, and residual D₂O was used as the internal reference. Data were processed using MestReNova software (Mestrelab Research).

### Raman Spectroscopy

Raman spectra were acquired using a Renishaw inVia confocal Raman spectrometer (Renishaw plc, UK). A 633 nm laser was used for excitation, and spectra were collected using a 50× objective, in a range of 300-3000 cm^-1^. The spectra were analysed with Origin 2018 software (OriginLab Corporation, Northampton, MA, USA) by using adjacent average smoothing with a 10-point window, and the baseline was corrected manually.

### DTNB assay to determine the thiol content of HASH and CSSH

The thiol content of thiolated hyaluronic acid (HASH) and thiolated chondroitin sulfate (CSSH) was quantified using Ellman’s assay according to the manufacturer’s protocol. Samples were dissolved in nitrogen-flushed reaction buffer (0.1 M Na₃PO₄, 1 mM EDTA, pH 8.0) prior to analysis. Absorbance measurements were performed at 412 nm using an Agilent Cary 60 UV-Vis spectrophotometer (Agilent Technologies, Santa Clara, CA, USA). Thiol concentrations were calculated from a calibration curve generated using N-acetylcysteine as a standard (Figure S7).

### Attenuated Total Reflectance Fourier-Transform Infrared (ATR-FTIR) spectroscopy

Prior to analysis, hydrogels were washed in Milli-Q water to remove residual PBS and freeze-dried. FTIR spectra were acquired using a Jasco FT/IR-4700A spectrometer (JASCO Ltd., ATR Pro ONE, Japan) equipped with a diamond ATR accessory and a DTGS detector. Spectra were recorded in the range of 400–4000 cm^−1^ at a resolution of 1 cm^−1^ with 128 scans. Data processing was performed using Jasco Spectra Manager software.

### SEM and SEM-EDS

Surface morphology and elemental composition of the hydrogels were analyzed by scanning electron microscopy (SEM) combined with energy-dispersive X-ray spectroscopy (EDS). Prior to analysis, hydrogels were washed in Milli-Q water, freeze-dried, and coated with a thin Au–Pd layer using a sputter coater. Measurements were performed using a Zeiss LEO 1550 SEM (Carl Zeiss AG, Oberkochen, Germany) equipped with an InLens detector and an Oxford Aztec EDS system (Oxford Instruments, Abingdon, UK), operated at an accelerating voltage of 2 kV.

### Verification of the presence of CSSH in the hydrogels by the dimethyl methylene blue (DMMB) assay

Glycosaminoglycan (CSSH) content of the hydrogels was quantified using the dimethylmethylene blue (DMMB) assay according to a published protocol^[44]^ with minor modifications. A pH 3 buffer was used both for hydrogel incubation and for preparation of the DMMB reagent. Hydrogels were incubated overnight in the pH 3 buffer, followed by incubation for 2 hours in DMMB reagent solution prepared in the same buffer. After washing with the pH 3 buffer, absorbance spectra in the hydrogels were recorded using a NanoDrop UV–Vis spectrophotometer (Thermo Fisher Scientific, USA), and data were processed using Origin 2018 software (OriginLab Corporation, Northampton, MA, USA).

### Labeling of the functionalized type I collagen with Alexa 633 and preparation of the Alexa-labeled hydrogels

Methacrylated collagen (MAC) was fluorescently labeled following a published protocol with minor modifications^[77]^. MAC was dissolved in carbonate buffer (0.1 M, pH 7.9) at 1 mg mL^−1^ and reacted with Alexa Fluor 633 at a ratio of 5 mg MAC per 76 µg dye. The reaction was performed at room temperature for 3 hours, followed by precipitation, centrifugation, and purification by dialysis. Alexa-labeled MAC was subsequently used for hydrogel preparation as described above. Partial bleaching of the fluorophore occurred during UV crosslinking but did not prevent fluorescence imaging.

### Labeling of human serum albumin (HSA), RM26, KWEA, and Exenatide with fluorescein isothiocyanate (FITC)

Human serum albumin (HSA), RM26, KWEA, and Exenatide were fluorescently labeled with fluorescein isothiocyanate (FITC) following the manufacturer’s protocol. Labeling was performed in carbonate buffer (0.1 M, pH 9.0) using a peptide-to-dye molar ratio of 1:1.3 or an HSA-to-dye mass ratio of 1:20. Reactions were carried out for 30 min at room temperature, followed by incubation at 4 °C overnight.

Unreacted FITC was removed by size-exclusion chromatography using PD MidiTrap G-10 or G-25 columns (Cytiva, Uppsala, Sweden) according to the manufacturer’s gravity-flow protocol, with Milli-Q water as eluent. For peptide conjugates, the eluate was directly frozen and lyophilized. For HSA, an additional purification step was performed by dialysis (12-14 kDa MWCO, Spectra/Por 4, Spectrum Labs, CA, USA) against Milli-Q water prior to lyophilization. All labeled products were freeze-dried before further use. GK3 and GK6 were obtained pre-labeled from the manufacturer.

### Verification of labeling of peptides or HSA by LC-MS

Analytical LC–MS was performed to confirm the FITC labeling. Separation was achieved on a reversed-phase C18 column using a water/acetonitrile gradient containing 0.05–0.1% formic acid. Detection was carried out by electrospray ionization mass spectrometry in positive-ion mode. Product formation was verified by comparison of the observed and calculated molecular masses and by comparison with the spectra of the unlabeled peptide.

### Measurement of HSA binding of FITC-labeled peptides by Microscale thermophoresis

Microscale thermophoresis (MST) was used to determine binding affinities between FITC-labeled peptides and human serum albumin (HSA). Measurements were performed in PBS supplemented with 0.05% (v/v) Tween-20. A serial dilution of HSA was prepared and incubated with the corresponding peptide prior to analysis. MST measurements were conducted using a Monolith NT instrument (NanoTemper Technologies, Munich, Germany), and data were analyzed using MO Affinity Analysis software (version 2.3). The experiments were carried out with a total of 4 replicates.

### Confocal microscopy and determination of the partitioning coefficient using image analysis

Partitioning of FITC-labeled dextrans, HSA, and peptides in hydrogels was quantified by confocal microscopy. FITC-labeled compounds were dissolved in PBS (pH 7.4, ionic strength 0.15 M) at 1 mg mL^−1^ for dextrans and 50 µM for peptides; partitioning of 70 kDa dextran and HSA was additionally evaluated at 7.5 and 25 mg mL^−1^. Hydrogel discs (~20 mg) were immersed in 150 µL of solution and incubated for 3 h at 4 °C. For HSA-dependent measurements, hydrogel discs were pre-incubated in HSA solutions for two days and subsequently transferred to peptide solutions containing the same HSA concentration.

Confocal images were acquired at 34 °C using a Zeiss LSM 700 confocal microscope (Carl Zeiss AG, Oberkochen, Germany) with a 25× water-immersion objective (ZEN Black). Images were collected over a 465 × 465 µm^2^ field of view at 4.4 pixels µm^−1^ and 50 µm depth from the gel surface. Time-dependent effects of HSA on peptide partitioning were assessed after incubation at 34 °C for 3 h (GK6 only) and 21 h using the same imaging settings. For 3D visualization, z-stacks were acquired from 20–60 µm depth with a 1.3 µm step size. Images were deconvolved and analyzed in Huygens Professional (v25.04, Scientific Volume Imaging).

Partitioning coefficients were determined using the Object Analyzer module of Huygens Professional. After deconvolution, segmentation was performed on the Alexa Fluor 633 channel (collagen signal) using automatic thresholding to define the collagen-rich region of interest (ROI). The selected ROI was transferred to the FITC channel (corresponding to the analyte), where fluorescence intensity within the hydrogel was quantified after removal of signal outside the ROI. For the determination of external fluorescence intensity, the inverse ROI was applied to the FITC channel, and a similar mean intensity measurement was carried out. The partitioning coefficient was calculated as the ratio of the mean fluorescence intensity inside and outside the hydrogel region.

### Ogston partitioning coefficient model fitting

The partitioning of the dextrans between the collagen-rich domains and the voids (K) can be predicted according to a model derived by Ogston^[46,78]^, as shown elsewhere^[51]^, in which the collagen-rich domain is regarded as a randomly oriented array of cylindrical fibers. For uncharged spherical macromolecules, such as the dextrans, the expression is as follows:

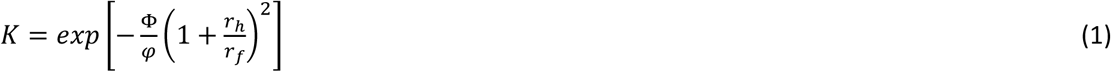

where Φ is the average volume fraction of polymer in the gel, *φ* is the volume fraction of the polymer-rich domains, r_h_ is the hydrodynamic radius of the sphere, and r_f_ is the radius of the polymer fibers. *φ* was determined by using 30 images from a Z-stack measurement and two additional images taken of Alexa-labeled hydrogels swollen in PBS using the same Zeiss microscope with the same setup mentioned in the previous paragraph. The raw images were deconvolved, and segmentation was carried out as mentioned before. The area inside and outside of the segmented ROI was measured, and *φ* was calculated as the ratio of the area inside to the area outside the ROI. Φ was calculated in the following way:

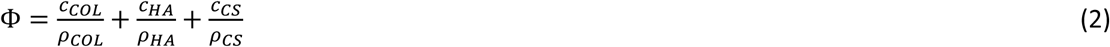

where *C_COL_* is the concentration of COL (7 g/l), *C_HA_* and *C_CS_* are the concentrations of HA and CS, respectively (*C_HA_* + *C_CS_* = 1.4 g/l), and *ρ_COL_*, *ρ_HA_* and *ρ_CS_* are the dry densities of COL (1.3 g/ml), HA (1.8 g/ml), and CS (2.5 g/ml), respectively. For all gel compositions, Eq. (2) gives Φ=0.006. The effective fiber radius r_f_ used in Eq. (2) was set to 0.6 nm. This is the weighted average value based on the relative total length of each polymer per volume of the gel, calculated from the volume fraction and radius of the respective polymer chain.

### Measurement of the self-diffusion coefficient of the dextrans, HSA, and the peptides by using Fluorescence recovery after photobleaching (FRAP)

FRAP measurements were performed on hydrogel discs (~20 mg) placed in 18-well Ibidi µ-slide plates and incubated in the respective solutions. For the determination of dextran self-diffusion, hydrogels were incubated in 120 µL of 1 mg mL^−1^ FITC–dextran solutions of varying molecular weights. Diffusion of 70 kDa dextran and HSA was assessed at concentrations of 1, 7.5, and 25 mg mL^−1^, where the highest concentration consisted of 7.5 mg mL^−1^ labeled and 17.5 mg mL^−1^ unlabeled compound.*

To determine peptide diffusion in the presence of dextran or HSA, hydrogels were pre-incubated in the corresponding dextran or HSA solutions at 4 °C for 24 h, followed by incubation in 50 µM peptide solutions containing the same additive concentration for 3 h at 4 °C. Time-dependent effects of HSA were evaluated after 21 h of incubation at 34 °C.

FRAP measurements were performed using a Zeiss LSM 700 confocal microscope (Carl Zeiss AG, Oberkochen, Germany) equipped with a 20× objective at 34 °C. Images were acquired over a 580 × 580 µm^2^ field of view at a resolution of 0.97 pixels µm^−1^. A rectangular bleaching region (100 × 100 µm^2^) was positioned at a fixed distance from the image edge, and bleaching was performed by increasing laser power to 100% until fluorescence was reduced to 70–80% of the initial intensity. Three pre-bleach images and 50 post-bleach images were recorded, with the acquisition interval adjusted to capture at least 95% fluorescence recovery.

Diffusion coefficients were extracted using the FRAP Solver software developed in-house (https://gitlab.com/jgernandt/frap-solver). Data were analyzed using the rectangular FRAP (rFRAP) model with pixel-wise double normalization*. A pure diffusion model was applied for measurements in PBS, while an immobile fraction rFRAP model was used for hydrogel measurements^[79]^.

\**Notes:*

*When only the labeled compound was used at a concentration of 25 mg mL^−1^, the extent of photobleaching was insufficient (<10%) to determine diffusion coefficients reliably*.

*Pixel-wise double normalization requires the selection of a reference region within the image, in which the average fluorescence intensity is used for normalization (see FRAP Solver documentation). Because the distribution of compounds within the hydrogels was frequently inhomogeneous, a region exhibiting comparable intensity and structural features was selected as the reference region (see examples in the Supplementary Information, section Supplementary Method 13, Figures S36-S42)*.

### Obstruction model for diffusion

A large number of models describing the diffusion of solutes in hydrogels have been derived and evaluated^[80]^. In the present work, we used the obstruction model by Amsden with the diffusion coefficient of the dextran probes with different sizes, based on Ogston’s expression for the distribution of spherical spaces between randomly oriented linear fibers^[47]^:

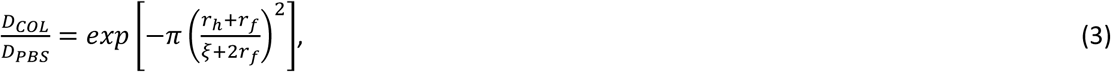

where *D_COL_* is the diffusion coefficient in the collagen-rich domain, *D_PBS_* is the diffusion coefficient in PBS, and *ξ* is the correlation length. In a recent article comparing the predictive ability of different models and measures of mesh size, Amsden showed that Eq. (3), in combination with the correlation length from scaling theory as the measure of mesh size, was found to be most consistent with experimental data^[48]^. In the present work, we determined the mesh size of the gel networks by fitting Eq. (3) to diffusion data (Kaleidagraph software, Levenberg-Marquardt algorithm) for dextran probes with different hydrodynamic radii, with *ξ* as a fitting parameter. The resulting effective mesh size was used in the modelling of the diffusion of peptides and HSA in the gels. The effective fiber radius *r_f_* was calculated as described above.

### Rheology

The storage (G′) and loss (G″) moduli of the hydrogels were measured using an ARES-G2 rheometer (TA Instruments, Sollentuna, Sweden) equipped with an 8 mm stainless-steel parallel upper plate and a 60 mm diameter APS flat lower plate. Measurements were performed at 34 °C using the Advanced Peltier System for temperature control. Oscillation amplitude sweeps were conducted at 1 Hz over a strain range of 0.1–20% to determine the linear viscoelastic (LVE) region. Frequency sweeps were subsequently performed within the LVE range at 1% strain over a frequency range of 0.1–10 Hz. During all measurements, the axial force was maintained at 40 ± 2 mN.

The mass of the hydrogel samples (8 mm diameter, 130 ± 10 mg) was measured in the swollen state. Following rheological characterization, gels were incubated in dextran or HSA solutions at 7.5 mg mL^−1^ and subsequently at 25 mg mL^−1^, and measurements were repeated after each incubation. After completion of rheological testing, gels were washed, freeze-dried, and used to determine mass and volumetric swelling ratios.

### Gradient diffusion measured by confocal microscopy

Gradient diffusion experiments were performed to simulate post-injection conditions and to assess peptide transport in the presence of human serum albumin (HSA). FITC-labeled GK6, RM26, and Exenatide were studied in the presence of 0, 7.5, and 25 mg mL^−1^ HSA. In addition, diffusion was evaluated using peptide solutions supplemented with 25 mg mL^−1^ HSA.

Experiments were conducted in 8-well Ibidi slides equipped with a custom 3D-printed insert separating the wells. The hydrogel precursor solution was injected into one compartment and crosslinked as described above. After 30 min of equilibration in air, PBS was added, the insert was removed, and the gels were washed prior to incubation in HSA solutions at the desired concentration.

Gradient diffusion measurements were performed using a Zeiss LSM 700 confocal microscope equipped with a 10× air objective at 34 °C. The focal plane was positioned at 120 µm from the bottom of the well at the hydrogel interface. After removal of the incubation medium, time-lapse imaging was initiated, and 80 µL of 50 µM FITC-labeled peptide solution was added to the opposing chamber. Images were acquired every 2 s during the first 2 min and subsequently every 5 s for the remainder of the experiment.

### Image analysis and determination of the diffusion coefficient from gradient diffusion measurements

Fluorescence intensity profiles were extracted from time-lapse image series using a custom ImageJ macro. Intensity distributions were obtained along three defined horizontal lines from each gel for each frame. For each setup, two separate gels were used. The resulting profiles were smoothed using adjacent averaging (20 points of window) prior to analysis.

Diffusion coefficients were determined by fitting experimentally obtained fluorescence profiles to numerical solutions of the diffusion equation, following the approach described by Degerstedt et al. ^[81]^ with minor modifications. Model fitting was performed using a Levenberg–Marquardt algorithm implemented in an in-house script. The upper boundary was defined at the top of the gel, applying a no-flux boundary condition (dc/dy = 0). For the lower limit, the concentration-time profile was approximated as a continuous piecewise linear function and constructed as described in Degerstedt et al^[81]^. Numerical fitting was performed using a temporal step size of Δt = 1.00 s and a spatial step size of Δy = 12.50 µm. For each experiment, data from four time points were fitted simultaneously over a spatial range extending from the defined lower boundary to the position at which the fluorescence intensity dropped below 5 in the initial image (t = 0 s).

### Determination of HSA binding of ABD-RM26’ by surface plasmon resonance

Surface plasmon resonance (SPR) measurements were performed using a Biacore T200 system (GE Healthcare, Uppsala, Sweden). HSA was immobilized on a CM5 sensor chip (Cytiva, Uppsala, Sweden) using standard amine coupling chemistry. Briefly, HSA was diluted to 2 µM in 10 mM sodium acetate buffer (pH 5.0) and injected over an EDC/NHS-activated sensor surface for 60 s at a flow rate of 2 µL/min. To evaluate peptide binding to HSA, ABD-RM26′ was diluted to 37 µM in running buffer (PBS supplemented with 0.05% Tween 20), followed by a three-fold serial dilution. Semaglutide was prepared in the same manner and used as a positive control. The resulting concentration series were injected over the immobilized HSA surface with a contact time of 60 s and a dissociation time of 120 s at a flow rate of 30 µL/min. The equilibrium binding constant (KD) was determined using Biacore T200 Evaluation Software by fitting the sensorgrams to a 1:1 Langmuir binding model.

### Radiolabeling of the peptides

GK6 and Exenatide were radiolabeled with ^125^I using N-succinimidyl p-(trimethylstannyl)benzoate as a prosthetic group, following a previously published protocol with minor modifications^[82]^. Radioiodination was performed at pH 5.0, followed by conjugation to the peptide at pH 9.0. Radiolabeled products were purified by size-exclusion chromatography prior to use.

Because RM26 does not contain a free amine group for radioiodination, an RM26 derivative with an albumin-binding domain (ABD-RM26′) bearing a DOTA chelator was synthesized and radiolabeled with ^111^In for SPECT–CT imaging^[83]^ (Supplementary information method section). Radiolabeling was performed in an ammonium acetate buffer at an elevated temperature. Radiochemical purity exceeded 95% as determined by ITLC (Tec-Control, Biodex).

### Subcutaneous injection and SPECT/CT analysis

The in vivo experiments were carried out in accordance with European standards for laboratory animal welfare under the permit number 00473/21, which is approved by the regional ethics committee for animal research (Uppsala, Sweden).

SPECT/CT imaging was performed to evaluate the *in vivo* distribution of radiolabeled peptides (Supplementary Information Method section 15). Female NMRI mice, 6-7 weeks old at arrival, underwent a one-week quarantine with acclimatization. The mice were kept at a maximum of 5 mice/cage with free access to food pellets, water, nest-building material, and hiding places, and had a 12/12 h light/darkness regime. Mice that lost weight during acclimatization were excluded. Randomization and blinding were not applied. Before injection, animals were shaved at the injection site and anesthetized with sevoflurane. Mice (n = 2 per peptide*) were injected subcutaneously above the right hind limb with 50 µL of peptide solution corresponding to a dose of 125 µg kg^−1^. Imaging was performed using a nanoScan SPECT/CT system (Mediso Medical Imaging Systems, Budapest, Hungary) as described in the supplementary information.

Following injection, a whole-body SPECT scan was acquired immediately and repeated at 5, 15, 30, 45, 60, 90, and 120 min post-injection. Animals were then allowed to recover for 1 h with free access to food and water, after which both CT and SPECT scans were repeated at 240 min post-injection. In additional experiments, one mouse was injected with GK6 and one with Exe using the same procedure described above, and SPECT/CT imaging was performed immediately after injection and again after 1 h, during which the animals were awake.

SPECT images were reconstructed as described in the Supplementary Information, and volumes of interest (VOIs) were defined at the injection site and in organs exhibiting radiotracer accumulation. Activity and volume values were analyzed using Origin 2018 software.

\**In accordance with the 3Rs principles of animal welfare (Reduction), a sample size of n = 2 mice per group was selected for this exploratory pilot study. This sample size was chosen to capture qualitative spatial distribution and clearance dynamics from the subcutaneous tissue within the same subject over time using continuous non-invasive SPECT/CT imaging, rather than to conduct formal statistical hypothesis testing or establish quantitative pharmacokinetic parameters*.

### Statistical analysis

Statistical analyses were performed using GraphPad Prism 8 (GraphPad Software, San Diego, CA, USA). Data normality was assessed using the Shapiro-Wilk test. When all datasets followed a normal distribution, statistical multiple comparisons were performed using one-way analysis of variance (ANOVA) with Tukey’s multiple comparisons test. Homoscedasticity was evaluated using Brown-Forsythe and Bartlett’s tests. If these tests indicated unequal variances, Welch’s ANOVA was applied instead.

When at least one dataset deviated from normality, statistical analysis was performed using the nonparametric Kruskal-Wallis test. For comparisons between two groups with normally distributed data, an unpaired Student’s t-test was used. Statistical significance was defined as p < 0.05 (*), p < 0.01 (**), p < 0.001 (***), and p < 0.0001 (****). The number of replicates used in the different experiments is indicated in the statistical table in the Supplementary Information as follows: N_g_ represents the number of independent hydrogels used, while N_m_ represents the number of measurements taken in each hydrogel. Further statistical analysis can be seen in the Supplementary information.

## Supporting information

Supplementary informations

## Acknowledgements

The authors are thankful for the help of the following people: Krisztina Juriga-Tóth (Uppsala University): SEM microscopy and SEM-EDS, Jeremy Adler (Uppsala University): FRAP measurements, confocal microscopy, and image analysis, Ákos György Juhász (Fondazione Casa Sollievo della Sofferenza): FTIR measurements, Luca D’Amario (Uppsala University): RAMAN measurements, Jasper van der Rijst: Image analysis (SVI BV), and Veronika Wingstedt (Uppsala University): *In vivo* experiments. ChatGPT 5.2 (accessed April 2026) was used to improve the text flow and cohesion, polish the language, and generate ImageJ and Excel macros for image analysis and data handling. Google Gemini version 3.6 Flash (accessed June 2026) was used as a tool to produce the initial concept and composition for the Table of Contents graphic. The authors reviewed and verified all content for scientific accuracy.

The work was financially supported by the Faculty of Medicine and Pharmacy at Uppsala University and Vinnova (Dnr 2019-00048 and 2024-03851) through the science program of the Swedish Drug Delivery Center (SweDeliver). Furthermore, the project was supported by the Swedish Cancer Society (Cancerfonden 23 2645 PjF 01H and 24 3889 Pj), and the Swedish Research Council (Vetenskaprådet, 2022-00556, 2021-03293 and 2022-04831). The molecular imaging work in this study was performed at the Preclinical PET−MRI Platform, a Swedish nationally available imaging platform at Uppsala University, Sweden, supported by the Knut and Alice Wallenberg Foundation.

## Author contribution

**D.J.:** Conceptualization, Methodology, Validation, Formal analysis, Investigation, Data Curation, Writing-Original Draft, Visualization; **A.N.:** Methodology, Investigation, Data Curation; **E.O.H.:** Methodology, Investigation; **J.W.H.:** Methodology, Investigation; **J.G.:** Methodology, Software, Formal analysis, Writing-Review & Editing; **L.O.:** Resources, Writing-Review & Editing, Supervision, Project administration, Funding acquisition; **A.O.:** Resources, Methodology, Investigation, Writing-Review & Editing, Project administration, Funding acquisition; **P.H.:** Conceptualization, Methodology, Validation, Formal analysis, Writing-Original Draft, Supervision, Project administration, Funding acquisition

