## Supplementary informations for "Roles of Interstitial Albumin in the Subcutaneous Delivery of Therapeutic Peptides"

**Supplementary Information**
**for**
**Roles of Interstitial Albumin in the Subcutaneous Delivery of Therapeutic Peptides**

David Juriga<sup>a,†</sup>, Anton Norberg<sup>a</sup>, Esther Olaniran Håkansson<sup>b</sup>, Jeroen Wilco van den Heuvel<sup>b</sup>, Johan Gråsjö<sup>a</sup>, Luke Odell<sup>b</sup>, Anna Orlova<sup>c</sup>, Per Hansson<sup>a,†</sup>

<sup>a</sup>Pharmaceutical Physical Chemistry, Department of Medicinal Chemistry, Faculty of Pharmacy, Uppsala University, Biomedicinskt Centrum BMC, Husargatan 3, 751 23, Uppsala, Sweden

<sup>b</sup>Preparative Medicinal Chemistry, Department of Medicinal Chemistry, Faculty of Pharmacy, Uppsala University, Biomedicinskt Centrum BMC, Husargatan 3, 751 23, Uppsala, Sweden

<sup>c</sup>Theranostics Group, Department of Medicinal Chemistry, Faculty of Pharmacy, Uppsala University, Biomedicinskt Centrum BMC, Husargatan 3, 751 23, Uppsala, Sweden

<sup>†</sup>Corresponding Authors:

Per Hansson,;

David Juriga,

Peptide structure and labeling analysis by LC-MS

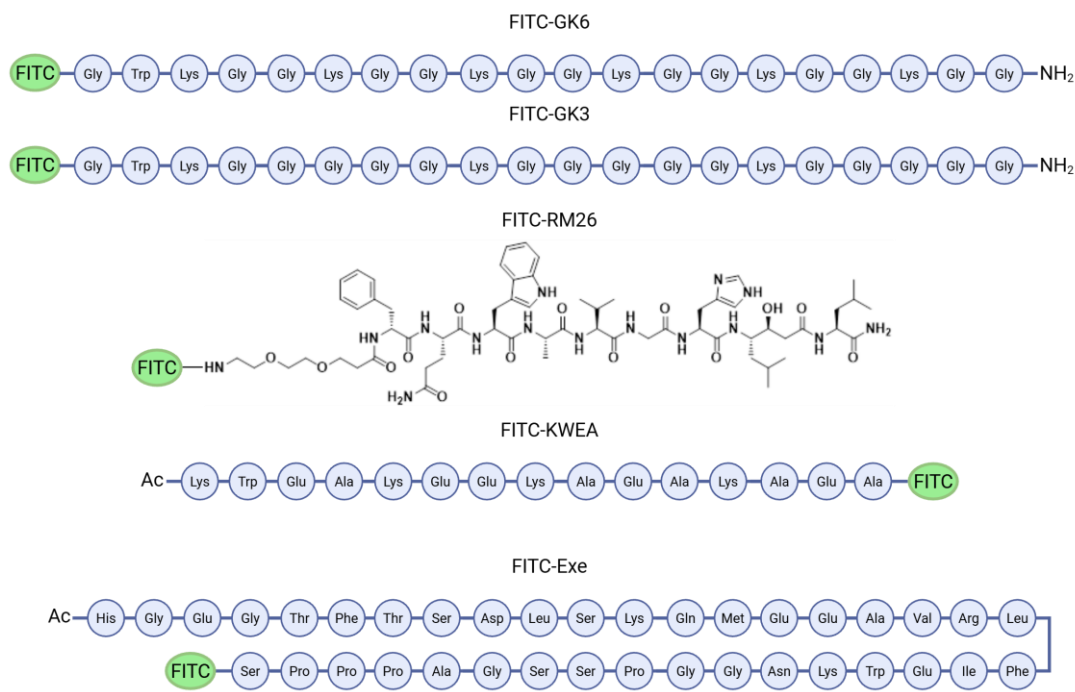

Figure S1: Structure of the FITC-labeled peptides

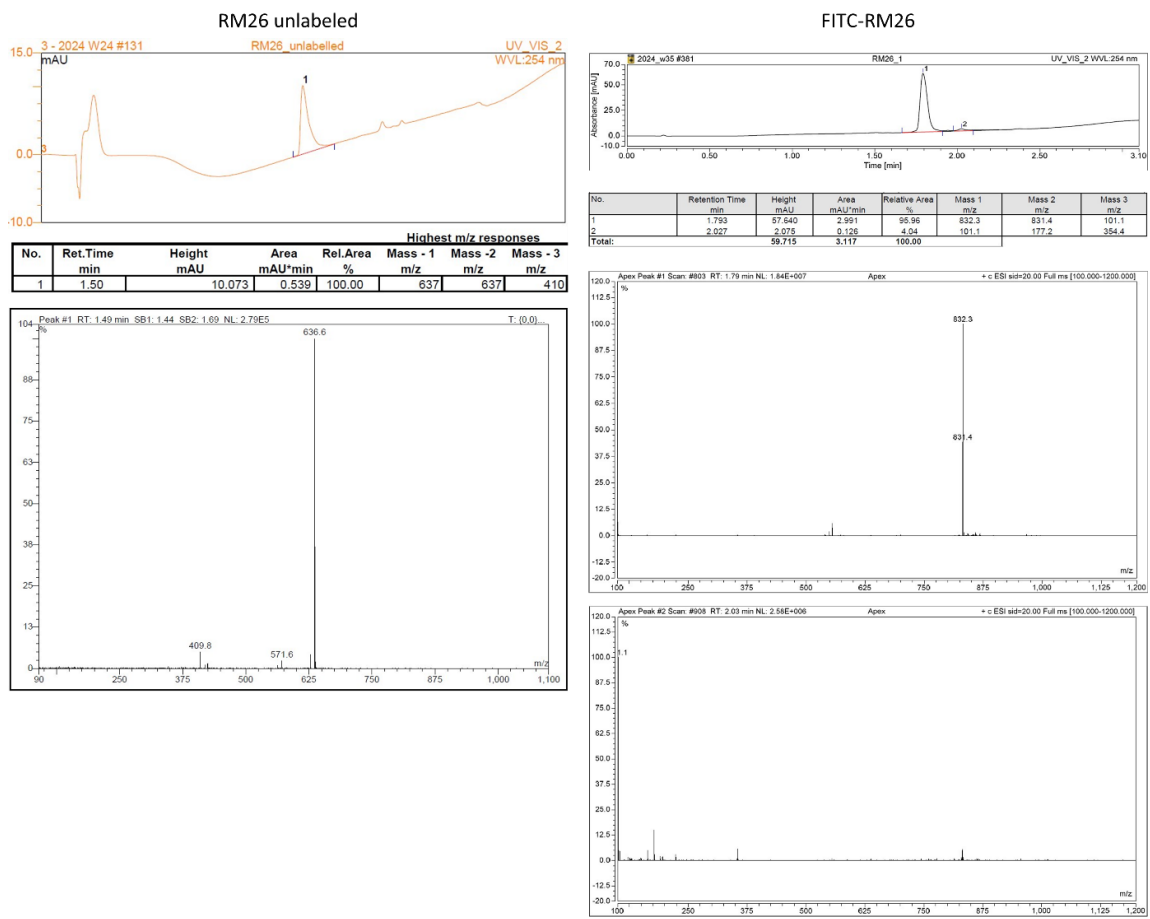

Figure S2: LC-MS spectra of unlabeled and FITC-labeled RM26

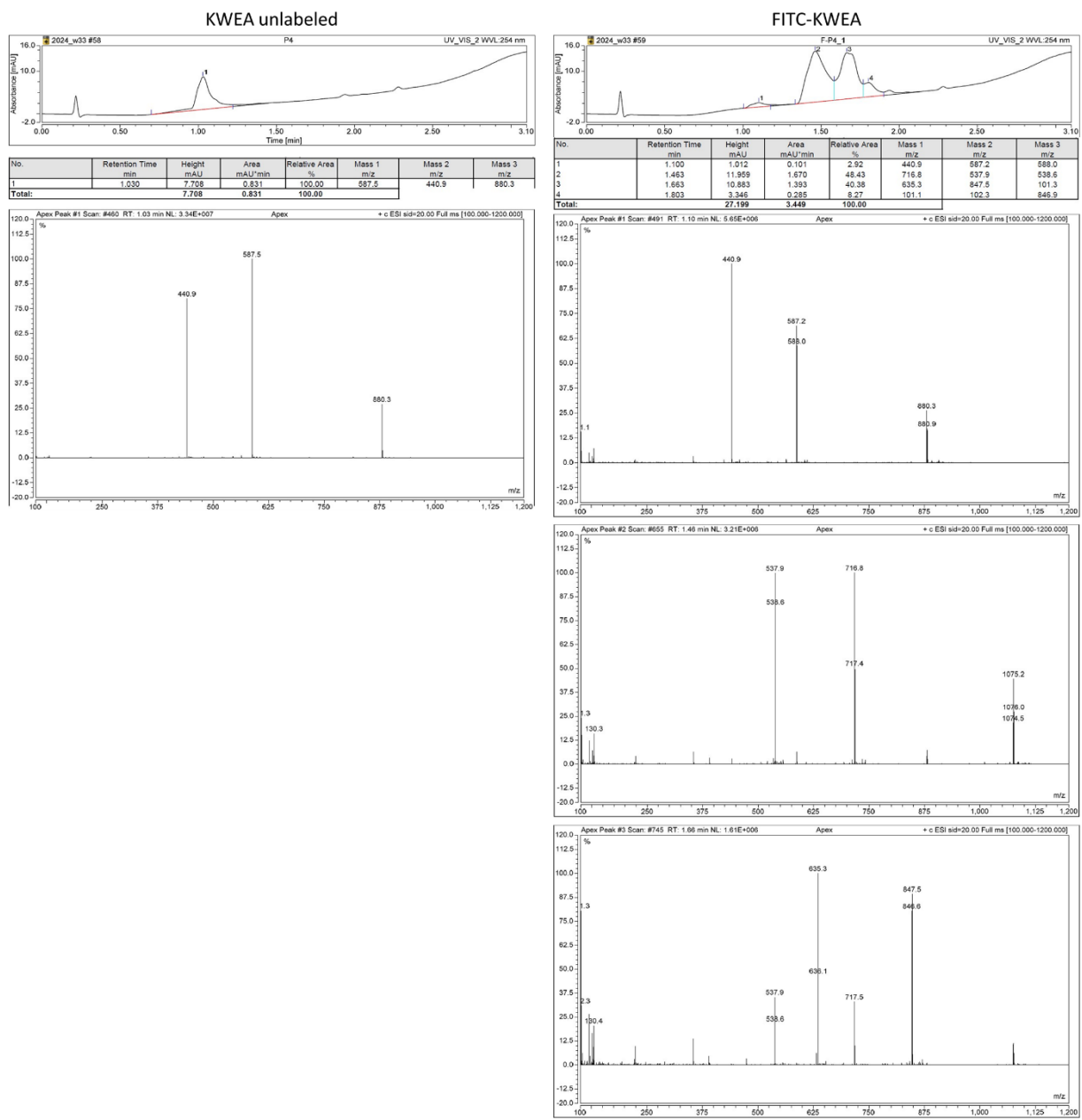

**Figure S3:** LC-MS spectra of unlabeled and FITC-labeled KWEA

#### Exe unlabeled

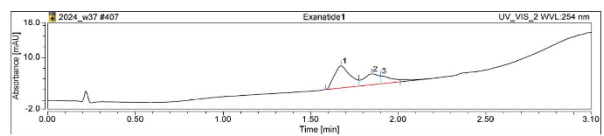

| No. | Retention Time<br>min | Height<br>mAU | Area<br>mAU*min | Relative Area<br>% | Mass 1<br>m/z | Mass 2<br>m/z | Mass 3<br>m/z |
| --- | --- | --- | --- | --- | --- | --- | --- |
| 1 | 1.673 | 5.144 | 0.539 | 59.14 | 1048.3 | 1047.4 | 838.8 |
| 2 | 1.850 | 2.518 | 0.250 | 27.46 | 1047.5 | 1048.3 | 838.9 |
| 3 | 1.900 | 1.800 | 0.122 | 13.40 | 1048.2 | 101.3 | 838.3 |
| Total: |  | 9.462 | 0.911 | 100.00 |  |  |  |

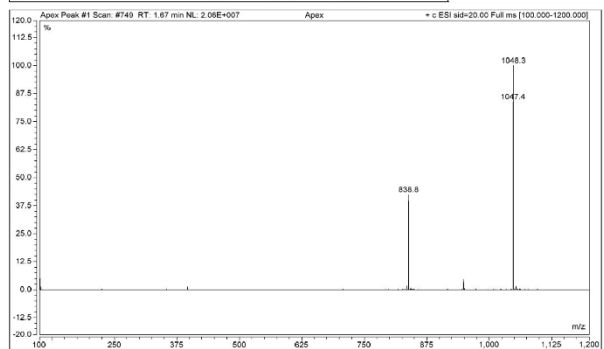

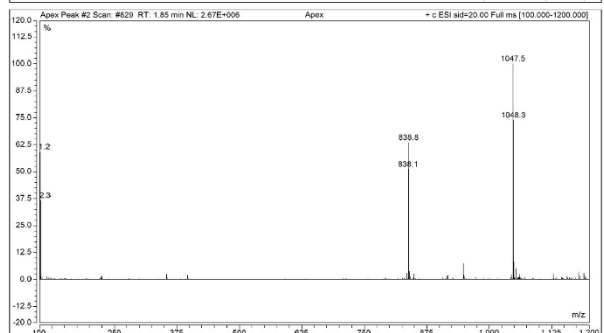

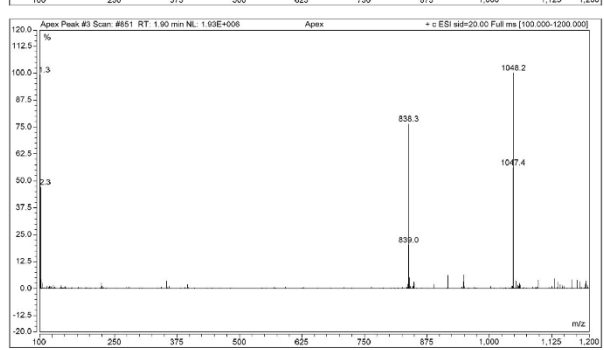

#### FITC-Exe

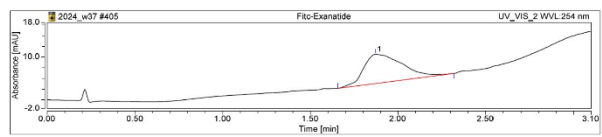

| No. | Retention Time<br>min | Height<br>mAU | Area<br>mAU*min | Relative Area<br>% | Mass 1<br>m/z | Mass 2<br>m/z | Mass 3<br>m/z |
| --- | --- | --- | --- | --- | --- | --- | --- |
| 1 | 1.673 | 6.848 | 1.708 | 100.00 | 916.1 | 1145.0 | 102.3 |
| Total: |  | 6.848 | 1.708 | 100.00 |  |  |  |

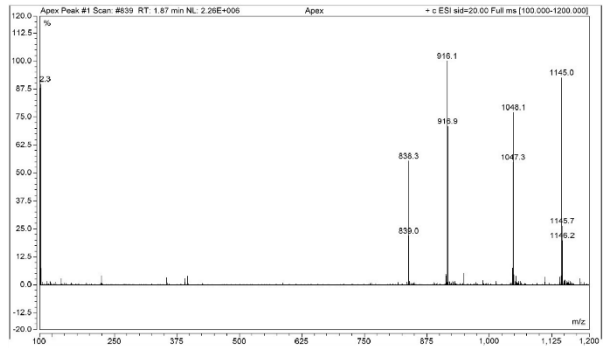

**Figure S4:** LC-MS spectra of unlabeled and FITC-labeled Exenatide

### Testing the albumin binding of the peptide by MST

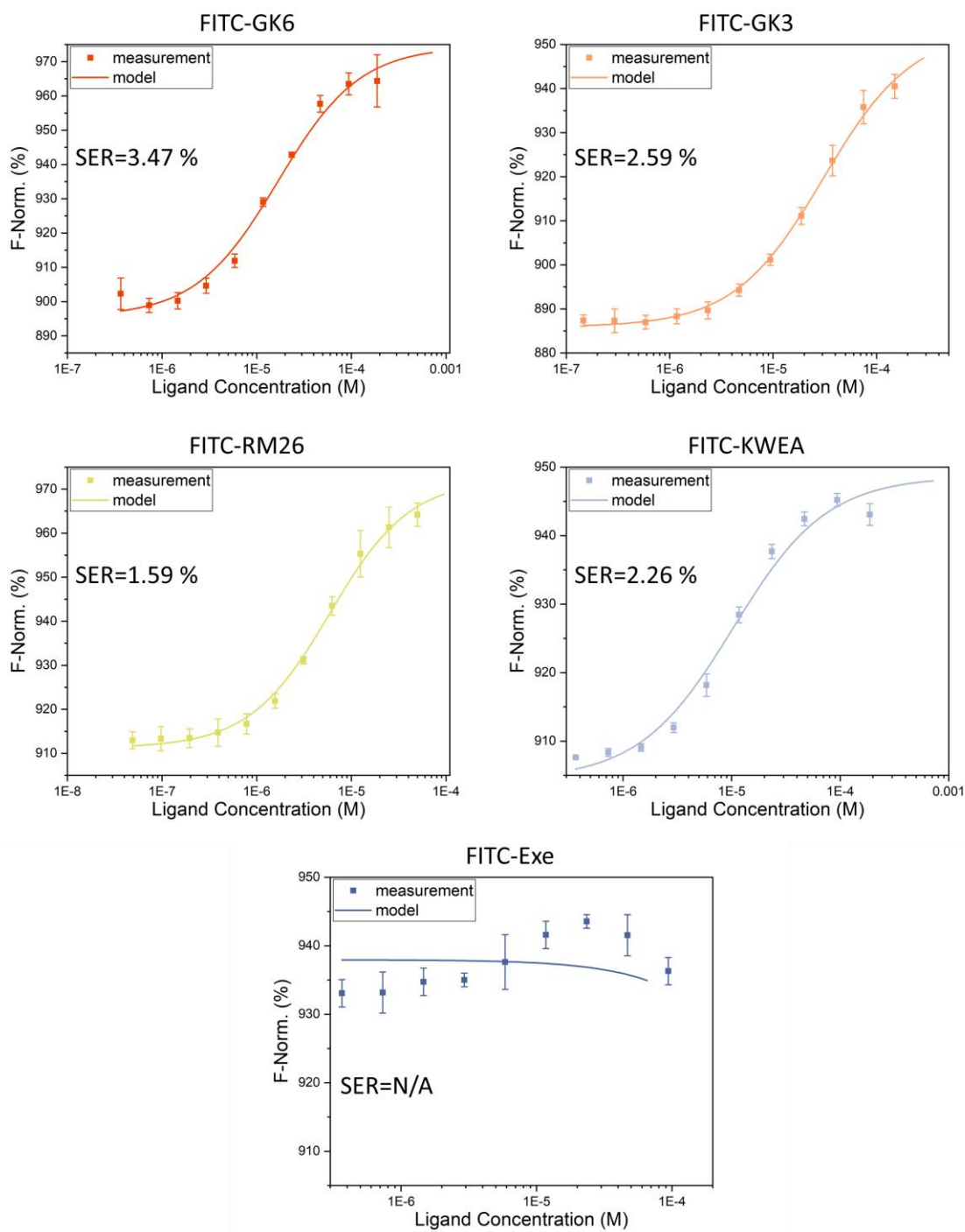

**Figure S5:** HSA binding isotherms of different peptides and the fitted model to determine the HSA binding constant

### Analysis of the functionalized hyaluronic acid (HASH) and chondroitin sulfate (CSSH)

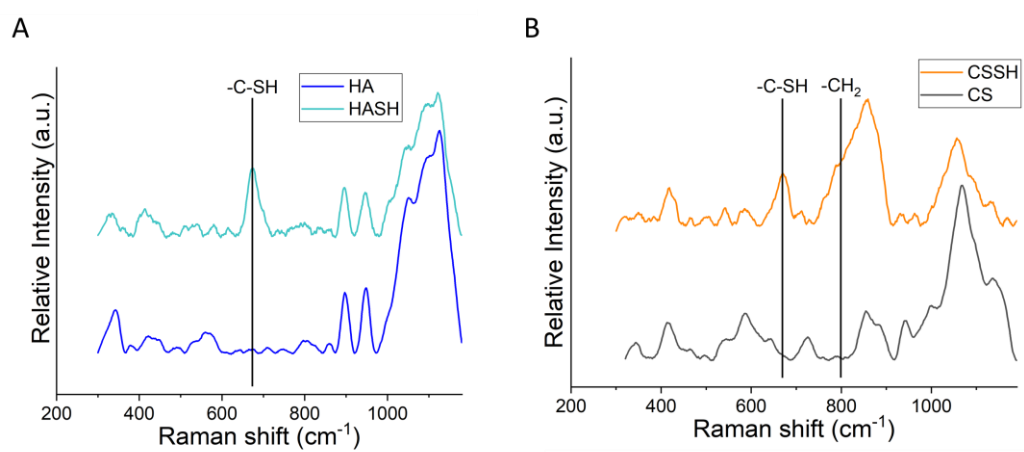

**Figure S6:** RAMAN spectra of unmodified and modified HA (A) and CS (B)

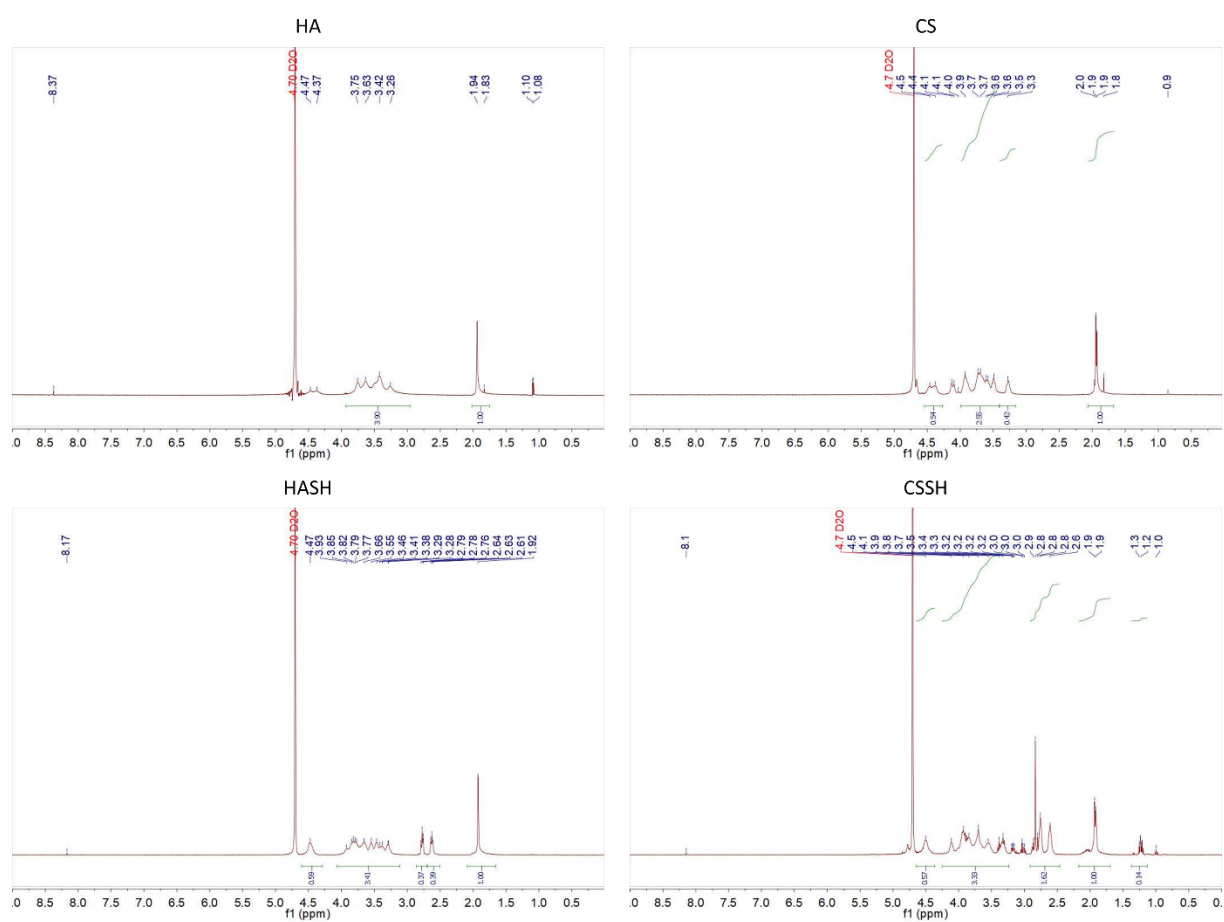

**Figure S7:** NMR spectra of the unmodified and modified HA and CS

The spectrum of HASH shows high similarity to that published by Rodler and coworkers<sup>[1]</sup>. The peaks at 2.63 and 2.78 ppm correspond to the protons of the methylene groups of DTP, while the peak at 1.85 ppm corresponds to the N-acetyl group of HA. Based on the ratio of the integrals, 38% of the carboxylic groups were modified. In the NMR spectrum of CSSH, the same peaks (2.6 and 2.8 ppm) appeared after modification. By comparing the integrals of these peaks with that at 1.9 ppm, approximately 80% modification of the carboxylic groups was determined. Since several peaks overlap in the NMR spectra, we also used the DTNB assay to quantify the degree of carboxylic group modification and to support the NMR results.

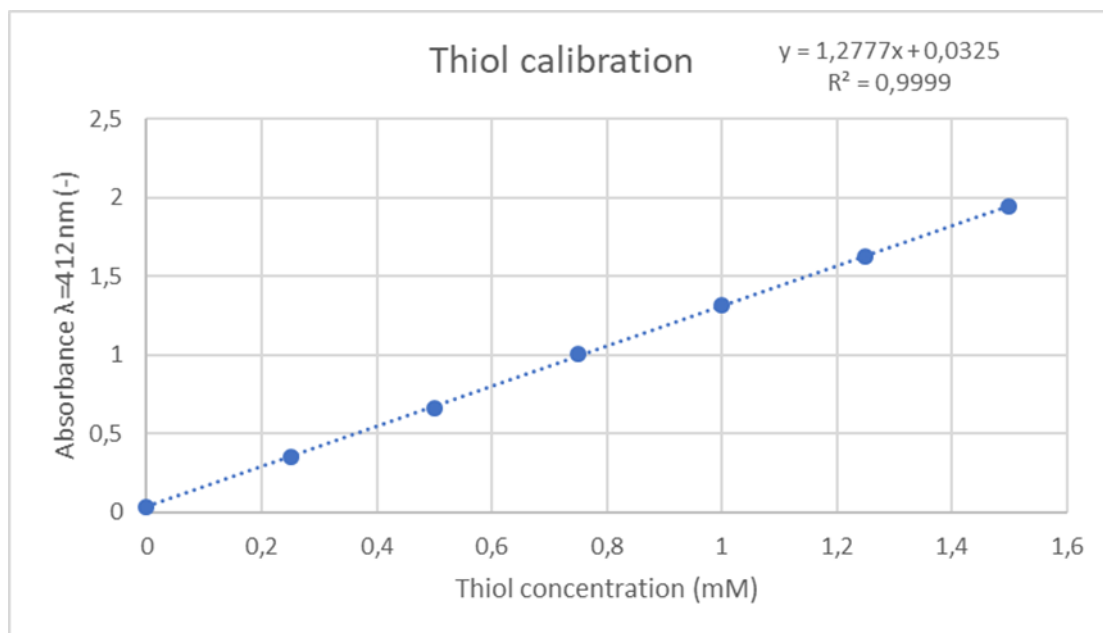

**Figure S8:** Thiol calibration used for DTNB measurement

##### FTIR and EDS spectra and EDS layered SEM images of the freeze-dried hydrogels

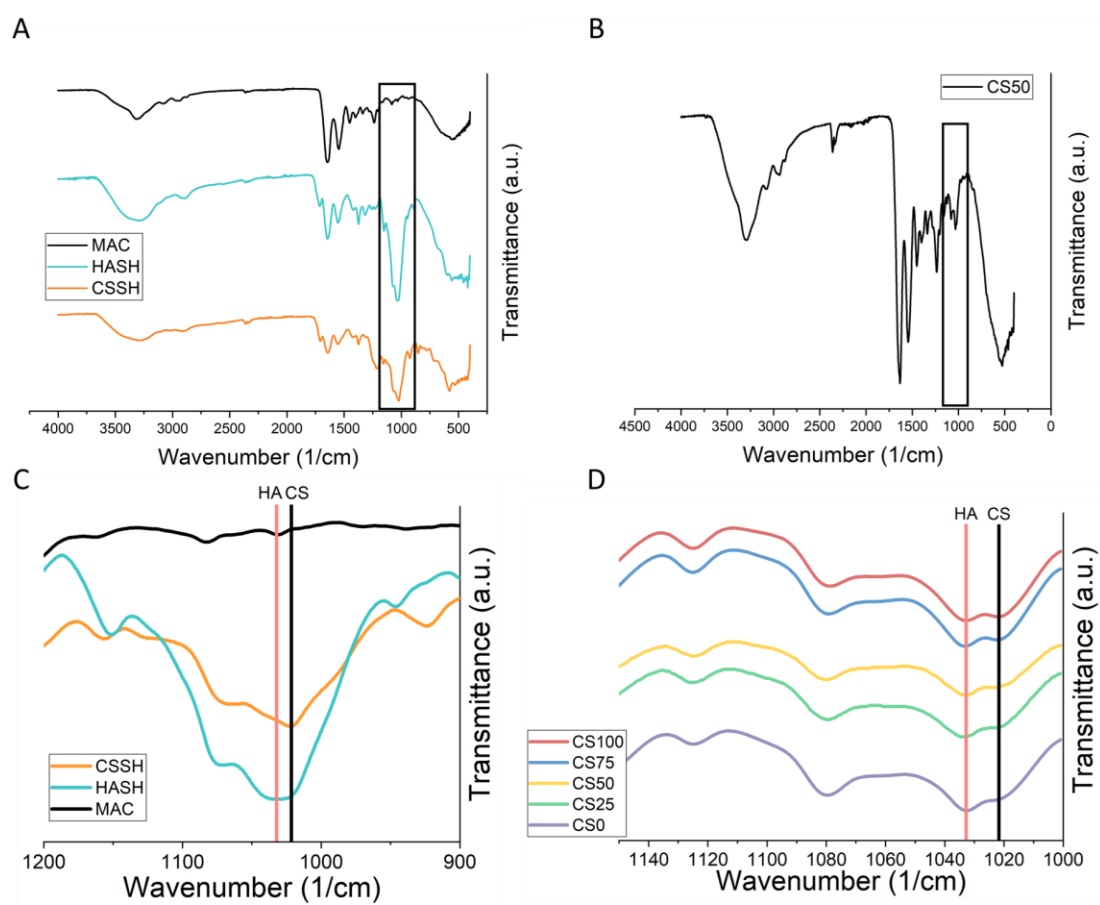

**Figure S9:** FTIR spectra of the different polymers (A) and a gel containing a 1:1 ratio of HASH and CSSH (B), and the zoomed-in spectra of the polymers (C) and the hydrogels (D)

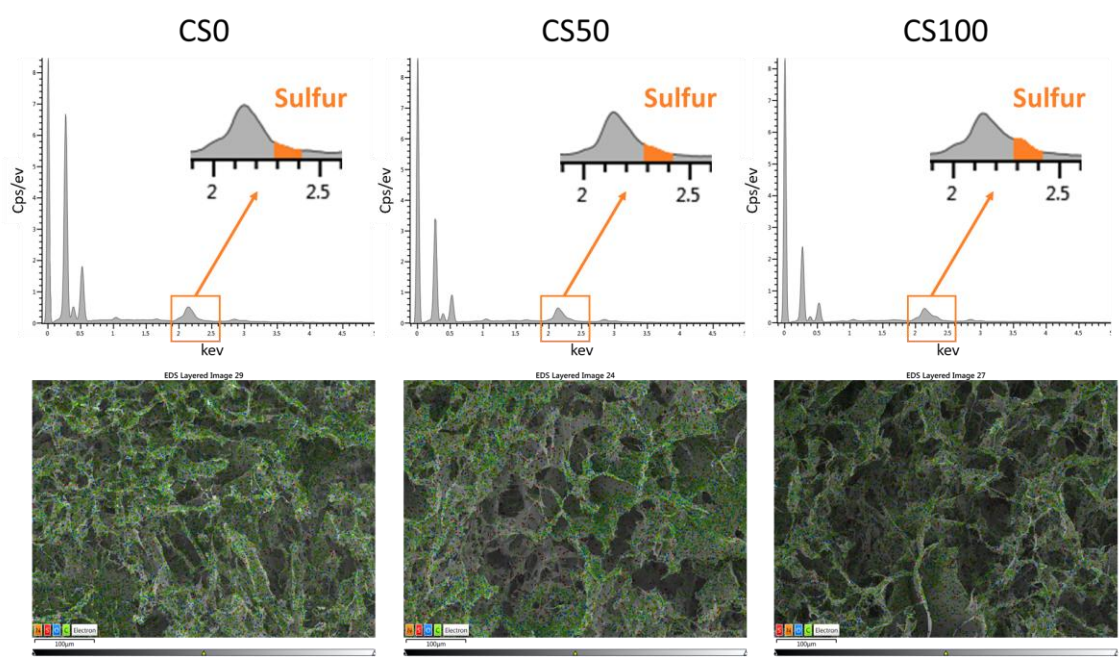

**Figure S10:** EDS spectra and EDS layered SEM images of the different hydrogels after freeze-drying

According to the atomic composition of the different hydrogels, CS0 contains 0.5 wt% sulfur, which belongs to the sulfur in the COL, while CS50 contains 1.5 wt%, and CS100 contains 3.2 wt%.

##### SEM images of the freeze-dried hydrogels and three-dimensional confocal images of the distribution of different dextrans in the CS50 hydrogel matrix

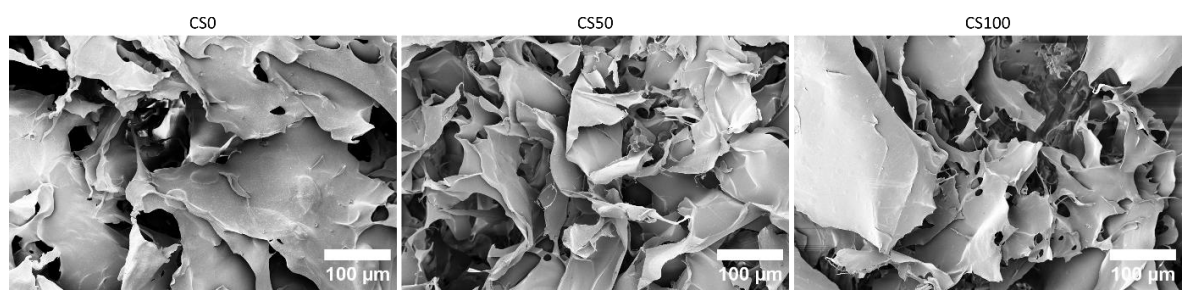

**Figure S11:** SEM images of the different hydrogels after freeze-drying

By using image analysis, we could quantify the volume fraction of the collagen-rich domains. While the volume fraction of the collagen-rich domains was measured as $0.54 \pm 0.027$ in the CS0, it was $0.62 \pm 0.014$ and $0.66 \pm 0.009$ in CS50 and CS100, respectively. These results proved to be statistically significant, indicating a 20% increase in the volume fraction of the collagen-rich domains.

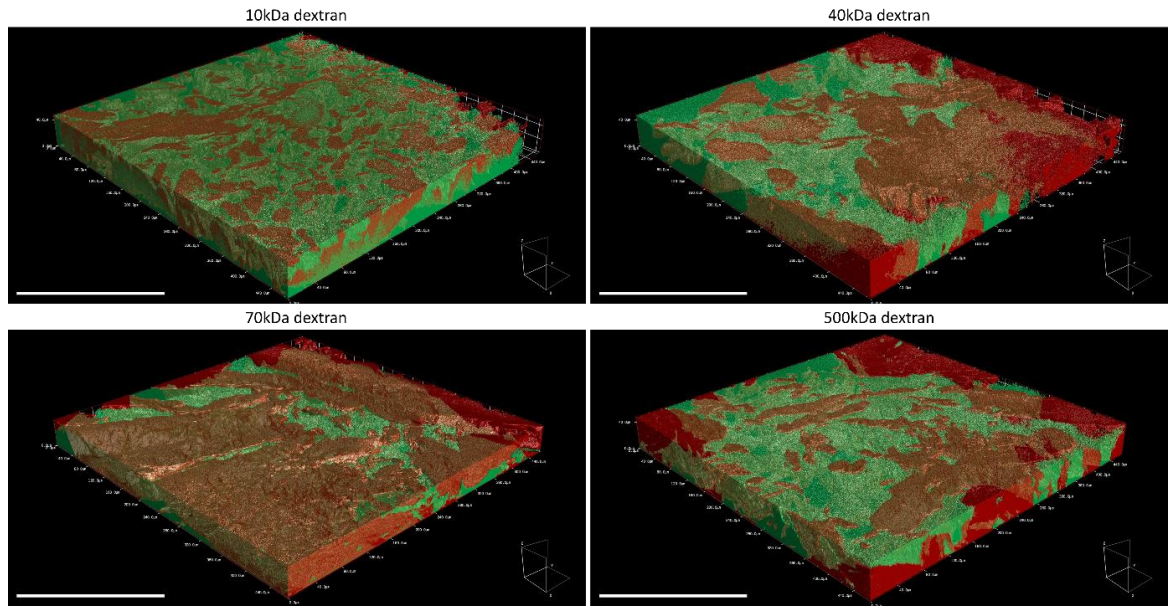

**Figure S12:** Three-dimensional distribution of the dextran probes in CS0 hydrogel

#### Analysis of the diffusion of the different dextran probes

**Table S1:** Statistical comparison of partitioning coefficients for the different dextran probes in each hydrogel (related to Figure 2C).

| Gel type CS0 |  |  |  |  | Gel type CS50 |  |  |  |  | Gel type CS100 |  |  |  |  |
| --- | --- | --- | --- | --- | --- | --- | --- | --- | --- | --- | --- | --- | --- | --- |
| Shapiro-Wilk test |  |  |  |  | Shapiro-Wilk test |  |  |  |  | Shapiro-Wilk test |  |  |  |  |
| P value | 0.8420 | 0.1298 | 0.3479 | 0.5643 | P value | 0.4515 | 0.4103 | 0.5726 | 0.0565 | P value | 0.7317 | 0.6956 | 0.0618 | 0.4222 |
| Passed normality test (alpha=0.05)? | Yes | Yes | Yes | Yes | Passed normality test (alpha=0.05)? | Yes | Yes | Yes | Yes | Passed normality test (alpha=0.05)? | Yes | Yes | Yes | Yes |
| P value summary | ns | ns | ns | ns | P value summary | ns | ns | ns | ns | P value summary | ns | ns | ns | ns |
| Brown-Forsythe test |  | Bartlett's test |  |  | Brown-Forsythe test |  | Bartlett's test |  |  | Brown-Forsythe test |  | Bartlett's test |  |  |
| 0.6317 (ns) |  | 0.4728 (ns) |  |  | 0.4054 (ns) |  | 0.0998 (ns) |  |  | 0.5981 (ns) |  | 0.2978 (ns) |  |  |
| Ordinary One-way Anova | 10kDa | 40kDa | 70kDa |  | Ordinary One-way Anova | 10kDa | 40kDa | 70kDa |  | Ordinary One-way Anova | 10kDa | 40kDa | 70kDa |  |
| 40kDa | 0.0004 (***) | X | X |  | 40kDa | 0.0007 (***) | X | X |  | 40kDa | 0.0002 (***) | X | X |  |
| 70kDa | <0.0001 (****) | 0.0036 (**) | X |  | 70kDa | <0.0001 (****) | 0.0002 (****) | X |  | 70kDa | <0.0001 (****) | 0.2051 (ns) | X |  |
| 500kDa | <0.0001 (****) | <0.0001 (****) | 0.0004 (***) |  | 500kDa | <0.0001 (****) | <0.0001 (****) | <0.0001 (****) |  | 500kDa | <0.0001 (****) | <0.0001 (****) | <0.0001 (****) |  |

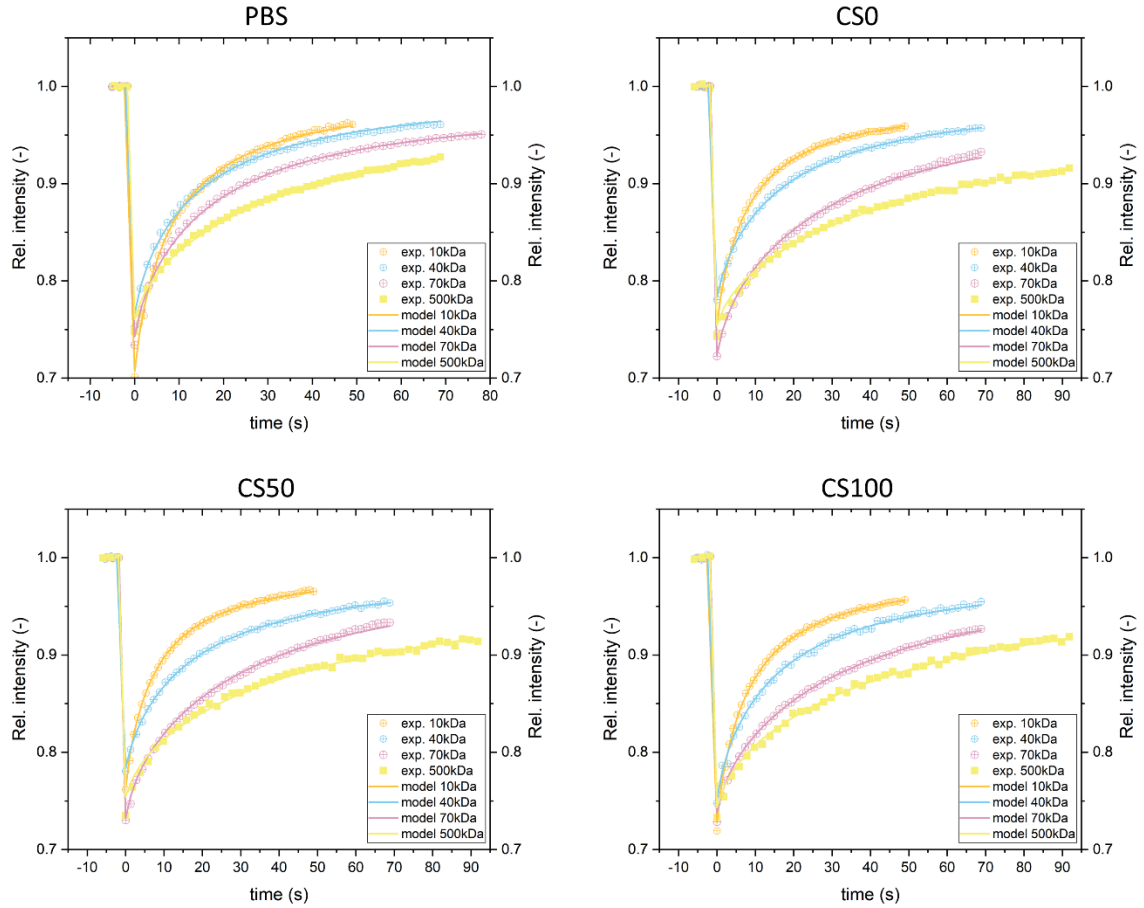

**Figure S13:** Examples of rFRAP model fitting on the measured recovery curves for each dextran probe

**Table S2:** Statistical comparison of diffusion coefficients for the different dextran probes in each medium (related to Figure 2D)

| $N_g=3$ $N_m=3/3/2$ | | | | | PBS | | | | |
| --- | --- | --- | --- | --- | --- | --- | --- | --- | --- |
| Shapiro-Wilk test | 10kDa | 40kDa | 70kDa | 500kDa | Shapiro-Wilk test | 10kDa | 40kDa | 70kDa | 500kDa |
| P value | 0.0576 | 0.8779 | 0.0762 | 0.0858 | P value | 0.5630 | 0.0911 | 0.9900 | 0.5079 |
| Passed normality test (alpha=0.05)? | Yes | Yes | Yes | Yes | Passed normality test (alpha=0.05)? | Yes | Yes | Yes | Yes |
| P value summary | ns | ns | ns | ns | P value summary | ns | ns | ns | ns |
| Brown-Forsythe test |  | Bartlett's test |  |  | Brown-Forsythe test |  | Bartlett's test |  |  |
| 0.1135 (ns) |  | 0.1008 (ns) |  |  | <0.0001 (****) |  | 0.0003 (***) |  |  |
| Ordinary one-way Anova | 10kDa | 40kDa | 70kDa |  | Anova test with Welch correction | 10kDa | 40kDa | 70kDa |  |
| 40kDa | <0.0001 (****) | X | X |  | 40kDa | <0.0001 (****) | X | X |  |
| 70kDa | <0.0001 (****) | <0.0001 (****) | X |  | 70kDa | <0.0001 (****) | <0.0001 (****) | X |  |
| 500kDa | <0.0001 (****) | <0.0001 (****) | 0.0001 (***) |  | 500kDa | <0.0001 (****) | <0.0001 (****) | <0.0001 (****) |  |

| Gel type CS50 |  |  |  |  | Gel type CS100 |  |  |  |  |
| --- | --- | --- | --- | --- | --- | --- | --- | --- | --- |
| Shapiro-Wilk test | 10kDa | 40kDa | 70kDa | 500kDa | Shapiro-Wilk test | 10kDa | 40kDa | 70kDa | 500kDa |
| P value | 0.3726 | 0.4341 | 0.1715 | 0.6272 | P value | 0.2556 | 0.9527 | 0.9985 | 0.2200 |
| Passed normality test (alpha=0.05)? | Yes | Yes | Yes | Yes | Passed normality test (alpha=0.05)? | Yes | Yes | Yes | Yes |
| P value summary | ns | ns | ns | ns | P value summary | ns | ns | ns | ns |
| Brown-Forsythe test |  | Bartlett's test |  |  | Brown-Forsythe test |  | Bartlett's test |  |  |
| 0.0017 (**) |  | <0.0001 (****) |  |  | 0.0162 (*) |  | 0.0199 (*) |  |  |
| Anova test with Welch correction | 10kDa | 40kDa | 70kDa |  | Anova test with Welch correction | 10kDa | 40kDa | 70kDa |  |
| 40kDa | <0.0001 (****) | X | X |  | 40kDa | <0.0001 (****) | X | X |  |
| 70kDa | <0.0001 (****) | <0.0001 (****) | X |  | 70kDa | <0.0001 (****) | <0.0001 (****) | X |  |
| 500kDa | <0.0001 (****) | <0.0001 (****) | <0.0001 (****) |  | 500kDa | <0.0001 (****) | <0.0001 (****) | <0.0001 (****) |  |

### HSA distribution in different hydrogels

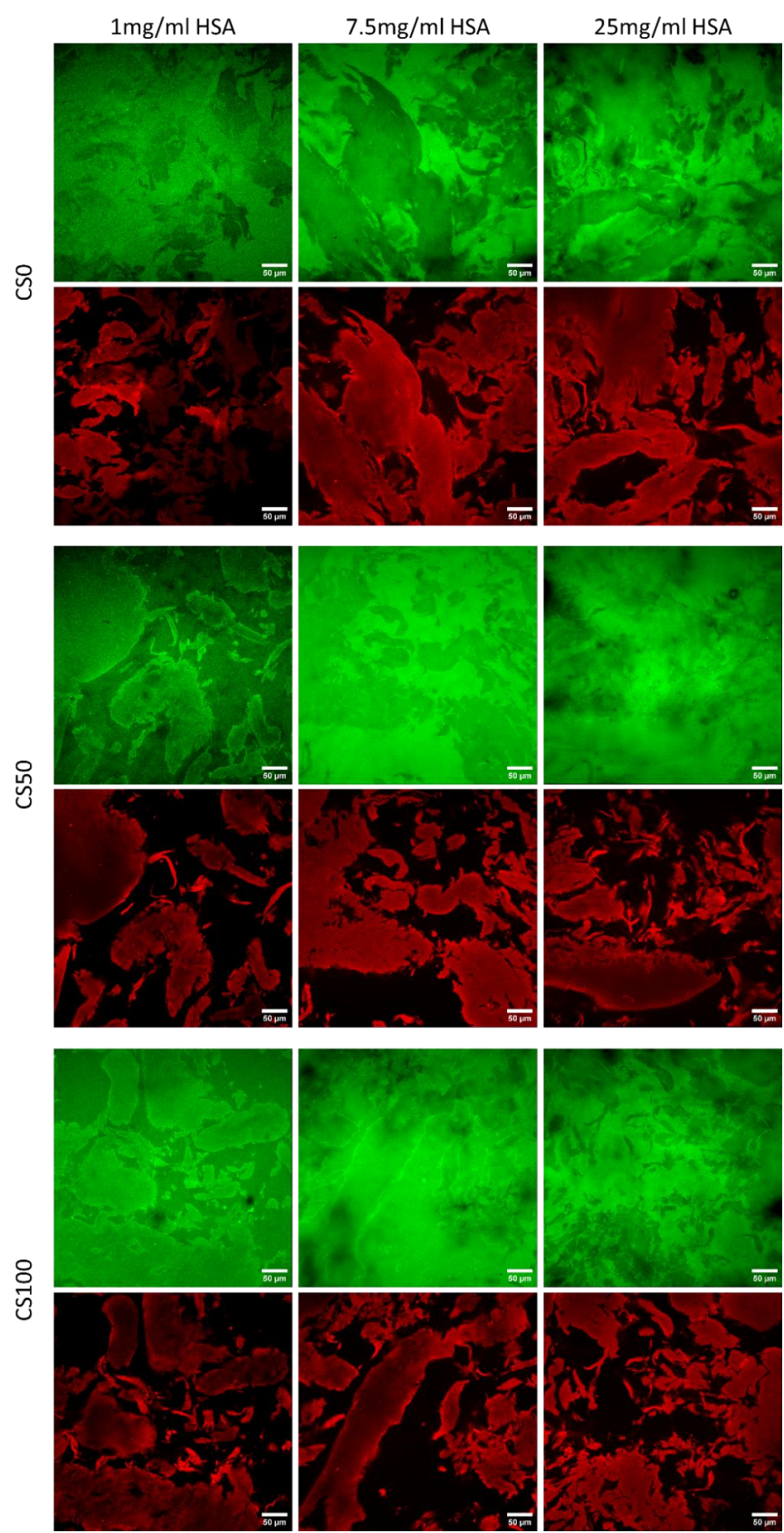

**Figure S14:** Concentration-dependent distribution of HSA in different hydrogels

**Table S3:** Statistical comparison of partitioning coefficients of HSA among hydrogel types at each concentration (related to Figure 3B)

| N<sub>0</sub>=3 N<sub>m</sub>=2/2/3 |  |  |  |  |  | 1mg/ml HSA |  |  |  |  |  | 7.5mg/ml HSA |  |  |  |  |  | 25mg/ml HSA |  |  |
| --- | --- | --- | --- | --- | --- | --- | --- | --- | --- | --- | --- | --- | --- | --- | --- | --- | --- | --- | --- | --- |
| Shapiro-Wilk test |  | CS0 | CS25 | CS50 | CS75 | CS100 | Shapiro-Wilk test |  | CS0 | CS25 | CS50 | CS75 | CS100 | Shapiro-Wilk test |  | CS0 | CS25 | CS50 | CS75 | CS100 |
| P value |  | 0.4259 | 0.0678 | 0.0561 | 0.0504 | 0.7536 | P value |  | 0.1529 | 0.1522 | 0.0717 | 0.2121 | 0.2023 | P value |  | 0.1583 | 0.0584 | 0.4473 | 0.1323 | 0.3821 |
| Passed normality test (alpha=0.05)? |  | Yes | Yes | Yes | Yes | Yes | Passed normality test (alpha=0.05)? |  | Yes | Yes | Yes | Yes | Yes | Passed normality test (alpha=0.05)? |  | Yes | Yes | Yes | Yes | Yes |
| P value summary |  | ns | ns | ns | ns | ns | P value summary |  | ns | ns | ns | ns | ns | P value summary |  | ns | ns | ns | ns | ns |
| Brown-Forsythe test |  |  | Bartlett's test |  |  | Brown-Forsythe test |  |  | Bartlett's test |  |  | Brown-Forsythe test |  |  | Bartlett's test |  |  |  |  |  |
| 0.6870 (ns) |  |  | 0.1889 (ns) |  |  | 0.2728 (ns) |  |  | 0.0142 (*) |  |  | 0.5740 (ns) |  |  | 0.3171 (ns) |  |  |  |  |  |
| Ordinary One-way Anova |  | CS0 | CS25 | CS50 | CS75 | Anova test with Welch correction |  | CS0 | CS25 | CS50 | CS75 | Ordinary One-way Anova |  | CS0 | CS25 | CS50 | CS75 |  |  |  |
| CS25 |  | 0.9442 (ns) | X | X | X | CS25 |  | 0.6003 (ns) | X | X | X | CS25 |  | 0.9018 (ns) | X | X | X |  |  |  |
| CS50 |  | 0.0237 (*) | 0.0036 (**) | X | X | CS50 |  | 0.4203 (ns) | 0.1150 (ns) | X | X | CS50 |  | 0.1155 (ns) | 0.0157 (*) | X | X |  |  |  |
| CS75 |  | 0.1667 (ns) | 0.0344 (*) | 0.8941 (ns) | X | CS75 |  | 0.0036 (**) | 0.0009 (***) | 0.157 (ns) | X | CS75 |  | 0.0023 (**) | 0.0002 (***) | 0.4964 (ns) | X |  |  |  |
| CS100 |  | 0.1137 (ns) | 0.0215 (*) | 0.9538 (ns) | 0.9996 (ns) | CS100 |  | <0.0001 (****) | 0.0003 (****) | 0.0286 (*) | 0.8906 (ns) | CS100 |  | 0.0376 (*) | 0.0042 (**) | 0.9850 (ns) | 0.8034 (ns) |  |  |  |

### Rheology

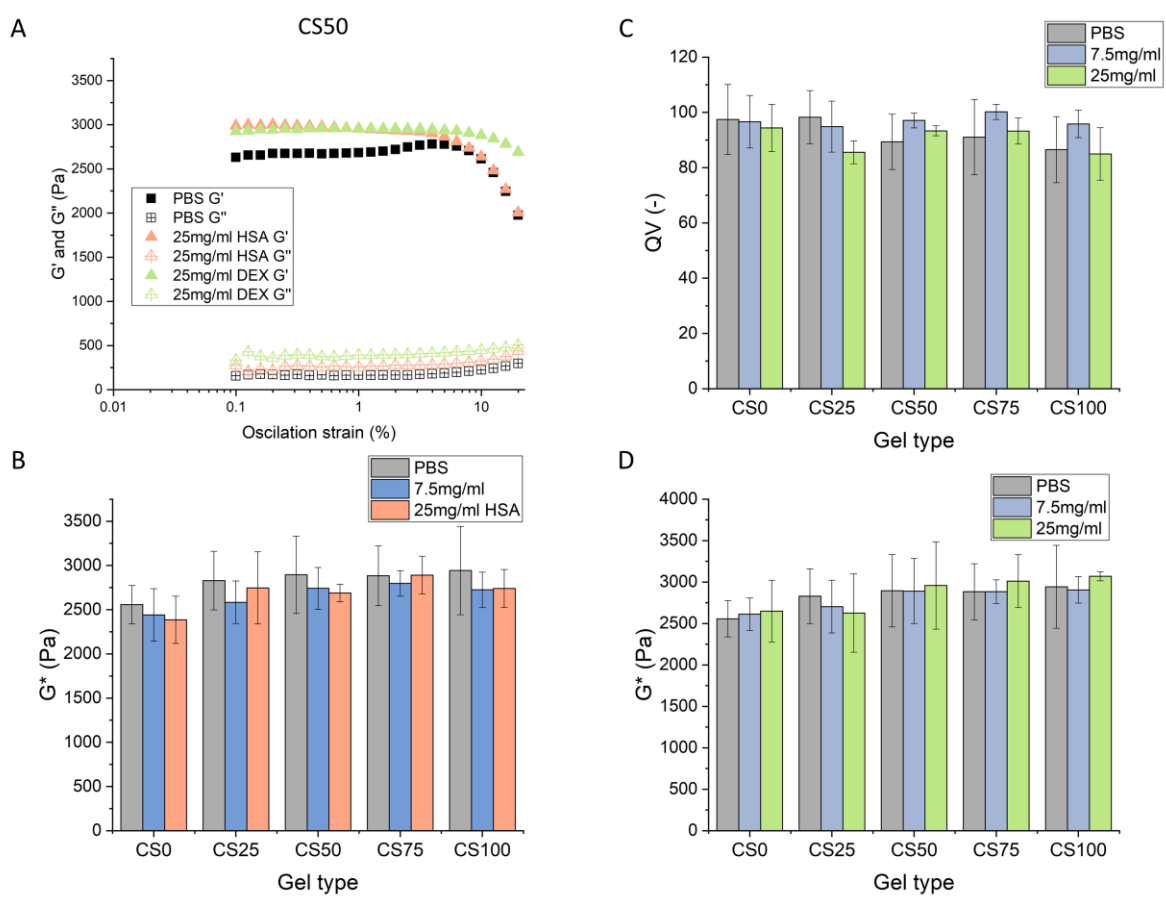

**Figure S15:** Oscillation strain curves of CS50 hydrogel after incubation in PBS, 25mg/ml dextran, and HSA solution (A), Complex modulus of the hydrogels measured at different HSA concentrations (B), Volumetric swelling degree (QV) (C), and complex modulus (D) of the hydrogels measured at different dextran concentrations

The oscillation strain measurement revealed no effect between 0.1-6% strain, while at higher deformation, the $G'$ started to decrease, while the $G''$ increased (Figure S15A). However, this effect was due to gel sliding between the rheometer geometries. To avoid this effect, we performed frequency oscillation measurements at 1% deformation (Figure S16). The measurements did not show any frequency-dependent behavior of the hydrogels, neither with nor without HSA and dextran. To compare the hydrogel types, we selected the complex modulus measured at 25 rad/s oscillation frequency (Figure 3C).

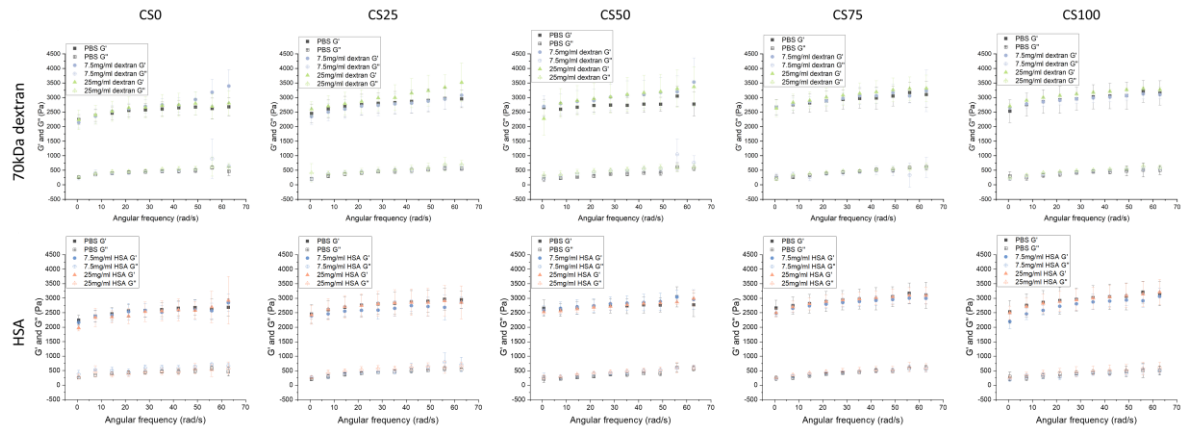

**Figure S16:** Frequency oscillation measurement of the different gels after incubation in HSA and dextran solutions with different concentrations

### Diffusion measurements of HSA and 70kDa dextran by FRAP

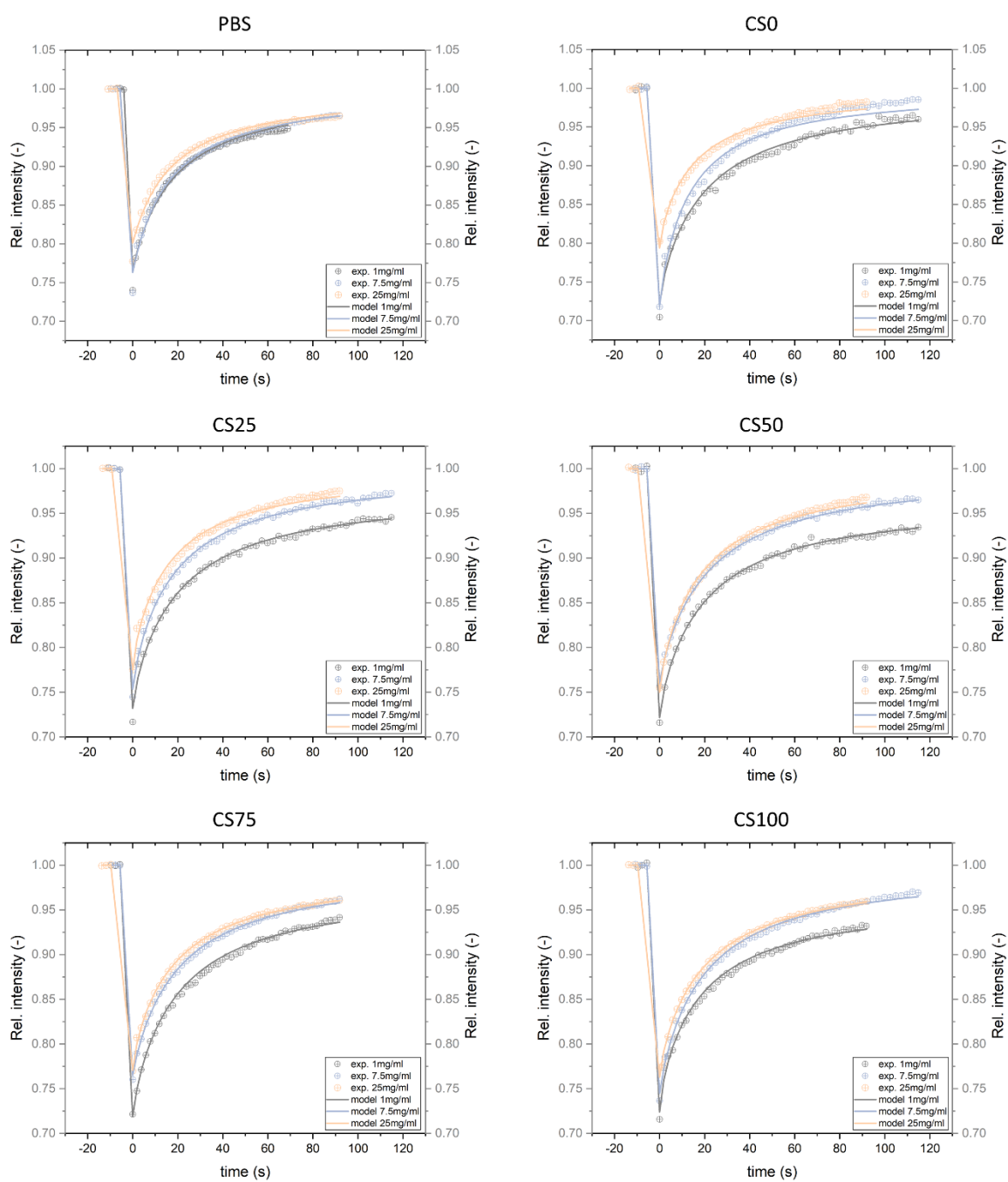

**Figure S17:** rFRAP model fitting on recovery curves of HSA measured at different concentrations in each medium

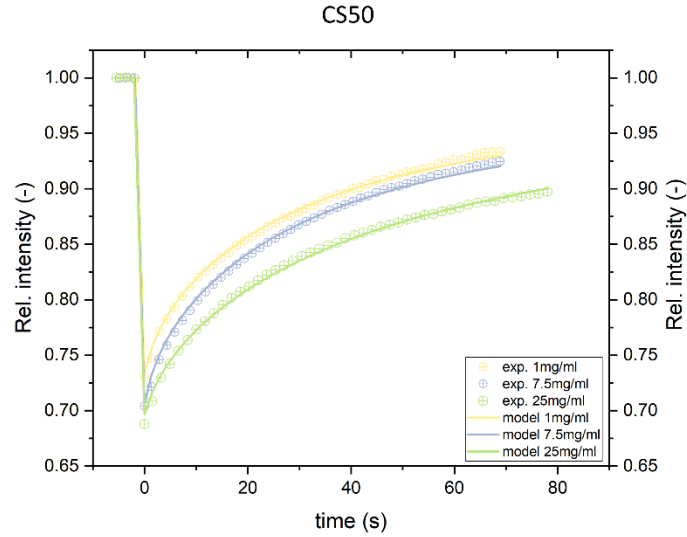

**Figure S18:** rFRAP model fitting on recovery curves of 70kDa dextran measured at different concentrations in the CS50 hydrogel

**Table S4:** Statistical comparison of diffusion coefficients of 70kDa dextran measured at different concentrations in each gel type (Related to Figure 3D)

|  |  |  |  |  |  |  |  |  |  |  |  |
| --- | --- | --- | --- | --- | --- | --- | --- | --- | --- | --- | --- |
| $N_g=3$ $N_m=2/3/3$ PBS | | | | Gel type CS0 | | | | Gel type CS25 | | | |
| Shapiro-Wilk test | 1mg/ml | 7.5mg/ml | 25mg/ml | Shapiro-Wilk test | 1mg/ml | 7.5mg/ml | 25mg/ml | Shapiro-Wilk test | 1mg/ml | 7.5mg/ml | 25mg/ml |
| P value | 0.2006 | 0.2840 | 0.5530 | P value | 0.9900 | 0.3065 | 0.0917 | P value | 0.8921 | 0.5891 | 0.5517 |
| Passed normality test (alpha=0.05)? | Yes | Yes | Yes | Passed normality test (alpha=0.05)? | Yes | Yes | Yes | Passed normality test (alpha=0.05)? | Yes | Yes | Yes |
| P value summary | ns | ns | ns | P value summary | ns | ns | ns | P value summary | ns | ns | ns |
| Brown-Forsythe test |  | Bartlett's test |  | Brown-Forsythe test |  | Bartlett's test |  | Brown-Forsythe test |  | Bartlett's test |  |
| 0.3742 (ns) |  | 0.1929 (ns) |  | 0.4549 (ns) |  | 0.2731 (ns) |  | 0.1330 (ns) |  | 0.2405 (ns) |  |
| Ordinary one-way Anova |  | 1mg/ml | 7.5mg/ml | Ordinary one-way Anova |  | 1mg/ml | 7.5mg/ml | Ordinary one-way Anova |  | 1mg/ml | 7.5mg/ml |
| 7.5mg/ml |  | 0.8975 (ns) | X | 7.5mg/ml |  | 0.8562 (ns) | X | 7.5mg/ml |  | 0.9989 (ns) | X |
| 25mg/ml |  | 0.0031 (**) | 0.0011 (**) | 25mg/ml |  | 0.0107 (*) | 0.0031 (**) | 25mg/ml |  | 0.0003 (***) | 0.0003 (***) |
| Gel type CS50 |  |  |  | Gel type CS75 |  |  |  | Gel type CS100 |  |  |  |
| Shapiro-Wilk test | 1mg/ml | 7.5mg/ml | 25mg/ml | Shapiro-Wilk test | 1mg/ml | 7.5mg/ml | 25mg/ml | Shapiro-Wilk test | 1mg/ml | 7.5mg/ml | 25mg/ml |
| P value | 0.1715 | 0.8346 | 0.7564 | P value | 0.3098 | 0.2000 | 0.7136 | P value | 0.9985 | 0.5347 | 0.6344 |
| Passed normality test (alpha=0.05)? | Yes | Yes | Yes | Passed normality test (alpha=0.05)? | Yes | Yes | Yes | Passed normality test (alpha=0.05)? | Yes | Yes | Yes |
| P value summary | ns | ns | ns | P value summary | ns | ns | ns | P value summary | ns | ns | ns |
| Brown-Forsythe test |  | Bartlett's test |  | Brown-Forsythe test |  | Bartlett's test |  | Brown-Forsythe test |  | Bartlett's test |  |
| 0.4968 (ns) |  | 0.2128 (ns) |  | 0.4861 (ns) |  | 0.1305 (ns) |  | 0.7695 (ns) |  | 0.6381 (ns) |  |
| Ordinary one-way Anova |  | 1mg/ml | 7.5mg/ml | Ordinary one-way Anova |  | 1mg/ml | 7.5mg/ml | Ordinary one-way Anova |  | 1mg/ml | 7.5mg/ml |
| 7.5mg/ml |  | 0.3240 (ns) | X | 7.5mg/ml |  | 0.9652 (ns) | X | 7.5mg/ml |  | 0.8386 (ns) | X |
| 25mg/ml |  | 0.0071 (**) | 0.0002 (***) | 25mg/ml |  | <0.0001 (****) | <0.0001 (****) | 25mg/ml |  | 0.0044 (**) | 0.0012 (**) |

### Distribution of different peptides in the hydrogel matrices

**Table S5:** Statistical comparison of partitioning coefficients of different peptides among hydrogel types (Related to Figure 4B)

| N<sub>g</sub>=2 N<sub>m</sub>=2/2 |  |  |  |  | GK6 |
| --- | --- | --- | --- | --- | --- |
| Brown-Forsythe test |  |  |  |  | Bartlett's test |
| 0.5926 (ns) |  |  |  |  | 0.3723 (ns) |
| Ordinary one-way Anova | CS0 | CS25 | CS50 | CS75 |  |
| CS25 | 0.9846 (ns) | X | X | X |  |
| CS50 | 0.2092 (ns) | 0.0876 (ns) | X | X |  |
| CS75 | 0.0385 (*) | 0.0144 (*) | 0.8683 (ns) | X |  |
| CS100 | <0.0001 (****) | <0.0001 (****) | 0.0013 (**) | 0.0082 (**) |  |

| GK3 |  |  |  |  | RM26 |  |  |  |  |
| --- | --- | --- | --- | --- | --- | --- | --- | --- | --- |
| Brown-Forsythe test |  |  |  |  | Bartlett's test |  |  |  |  |
| 0.6371 (ns) |  |  |  |  | 0.3808 (ns) |  |  |  |  |
| Ordinary one-way Anova | CS0 | CS25 | CS50 | CS75 | Anova test with Welch correction | CS0 | CS25 | CS50 | CS75 |
| CS25 | 0.5144 (ns) | X | X | X | CS25 | 0.9949 (ns) | X | X | X |
| CS50 | 0.3382 (ns) | 0.9970 (ns) | X | X | CS50 | 0.9722 (ns) | 0.9744 (ns) | X | X |
| CS75 | 0.9006 (ns) | 0.1461 (ns) | 0.0816 (ns) | X | CS75 | 0.8854 (ns) | 0.5069 (ns) | 0.8795 (ns) | X |
| CS100 | 0.8765 (ns) | 0.1313 (ns) | 0.0728 (ns) | >0.9999 (ns) | CS100 | 0.9426 (ns) | 0.9673 (ns) | >0.9999 (ns) | 0.7650 (ns) |

| KWEA |  |  |  |  | Exe |  |  |  |  |
| --- | --- | --- | --- | --- | --- | --- | --- | --- | --- |
| Brown-Forsythe test |  |  |  |  | Bartlett's test |  |  |  |  |
| 0.8823 (ns) |  |  |  |  | 0.7965 (ns) |  |  |  |  |
| Ordinary one-way Anova | CS0 | CS25 | CS50 | CS75 | Ordinary One-way Anova | CS0 | CS25 | CS50 | CS75 |
| CS25 | 0.4120 (ns) | X | X | X | CS25 | 0.2971 (ns) | X | X | X |
| CS50 | 0.6293 (ns) | 0.9950 (ns) | X | X | CS50 | >0.9999 (ns) | 0.3009 (ns) | X | X |
| CS75 | 0.7620 (ns) | 0.9707 (ns) | 0.9993 (ns) | X | CS75 | 0.3388 (ns) | 0.0091 (**) | 0.3347 (ns) | X |
| CS100 | 0.1387 (ns) | 0.9478 (ns) | 0.8017 (ns) | 0.6740 (ns) | CS100 | 0.7352 (ns) | 0.0359 (*) | 0.7302 (ns) | 0.9505 (ns) |

**Table S6:** Statistical comparison of the partitioning coefficient of GK6 (Related to Figure S18 and Figure 4C)

| Effect of the HSA concentration in each gel type |  |  |  |  |  |  |  |  |  |  |  |  |  |  |
| --- | --- | --- | --- | --- | --- | --- | --- | --- | --- | --- | --- | --- | --- | --- |
| N<sub>g</sub>=2 N<sub>m</sub>=2/2 |  |  |  |  | Gel type CS25 |  |  |  |  | Gel type CS50 |  |  |  |  |
| Gel type CS0 |  |  |  |  | Gel type CS25 |  |  |  |  | Gel type CS50 |  |  |  |  |
| Shapiro-Wilk test | PBS | 1mg/ml | 7.5mg/ml | 25mg/ml | Shapiro-Wilk test | PBS | 1mg/ml | 7.5mg/ml | 25mg/ml | Shapiro-Wilk test | PBS | 1mg/ml | 7.5mg/ml | 25mg/ml |
| P value | 0.6586 | 0.5184 | 0.9896 | 0.5176 | P value | 0.3187 | 0.5562 | 0.8919 | 0.0922 | P value | 0.7806 | 0.9983 | 0.7783 | 0.4865 |
| Passed normality test (alpha=0.05)? | Yes | Yes | Yes | Yes | Passed normality test (alpha=0.05)? | Yes | Yes | Yes | Yes | Passed normality test (alpha=0.05)? | Yes | Yes | Yes | Yes |
| P value summary | ns | ns | ns | ns | P value summary | ns | ns | ns | ns | P value summary | ns | ns | ns | ns |
| Brown-Forsythe test |  | Bartlett's test |  |  | Brown-Forsythe test |  | Bartlett's test |  |  | Brown-Forsythe test |  | Bartlett's test |  |  |
| 0.7304 (ns) |  | 0.8572 (ns) |  |  | 0.5592 (ns) |  | 0.3051 (ns) |  |  | 0.2618 (ns) |  | 0.0826 (ns) |  |  |
| Ordinary One-way Anova | PBS | 1mg/ml | 7.5mg/ml |  | Ordinary One-way Anova | PBS | 1mg/ml | 7.5mg/ml |  | Ordinary One-way Anova | PBS | 1mg/ml | 7.5mg/ml |  |
| 1mg/ml | 0.1321 (ns) | X | X |  | 1mg/ml | 0.9476 (ns) | X | X |  | 1mg/ml | 0.9982 (ns) | X | X |  |
| 7.5mg/ml | 0.0004 (***) | 0.0164 (*) | X |  | 7.5mg/ml | 0.8036 (ns) | 0.9756 (ns) | X |  | 7.5mg/ml | 0.0018 (**) | 0.0015 (**) | X |  |
| 25mg/ml | 0.0007 (***) | 0.0269 (*) | 0.9938 (ns) |  | 25mg/ml | 0.0060 (**) | 0.0112 (*) | 0.0362 (*) |  | 25mg/ml | 0.0007 (***) | 0.0006 (***) | 0.9550 (ns) |  |

| Gel type CS75 |  |  |  |  | Gel type CS100 |  |  |  |  |
| --- | --- | --- | --- | --- | --- | --- | --- | --- | --- |
| Shapiro-Wilk test | PBS | 1mg/ml | 7.5mg/ml | 25mg/ml | Shapiro-Wilk test | PBS | 1mg/ml | 7.5mg/ml | 25mg/ml |
| P value | 0.5004 | 0.4818 | 0.7819 | 0.3549 | P value | 0.4212 | 0.6469 | 0.6889 | 0.7728 |
| Passed normality test (alpha=0.05)? | Yes | Yes | Yes | Yes | Passed normality test (alpha=0.05)? | Yes | Yes | Yes | Yes |
| P value summary | ns | ns | ns | ns | P value summary | ns | ns | ns | ns |
| Brown-Forsythe test |  | Bartlett's test |  |  | Brown-Forsythe test |  | Bartlett's test |  |  |
| 0.0493 (*) |  | 0.0010 (**) |  |  | 0.8224 (ns) |  | 0.5788 (ns) |  |  |
| Anova test with Welch correction | PBS | 1mg/ml | 7.5mg/ml |  | Ordinary One-way Anova | PBS | 1mg/ml | 7.5mg/ml |  |
| 1mg/ml | 0.9985 (ns) | X | X |  | 1mg/ml | 0.2551 (ns) | X | X |  |
| 7.5mg/ml | 0.2928 (ns) | 0.3480 (ns) | X |  | 7.5mg/ml | 0.0193 (*) | 0.3644 (ns) | X |  |
| 25mg/ml | 0.0496 (*) | 0.0430 (*) | 0.0157 (*) |  | 25mg/ml | 0.0005 (***) | 0.0082 (**) | 0.1971 (ns) |  |

| Effect of the gel types at each HSA concentration |  |  |  |  |  |  |  |  |  |  |  |  |  |  |
| --- | --- | --- | --- | --- | --- | --- | --- | --- | --- | --- | --- | --- | --- | --- |
| 1mg/ml |  |  |  |  | 7.5mg/ml |  |  |  |  | 25mg/ml |  |  |  |  |
| Brown-Forsythe test |  | Bartlett's test |  |  | Brown-Forsythe test |  | Bartlett's test |  |  | Brown-Forsythe test |  | Bartlett's test |  |  |
| 0.1767 (ns) |  | 0.0610 (ns) |  |  | 0.1953 (ns) |  | 0.0189 (*) |  |  | 0.6104 (ns) |  | 0.2952 (ns) |  |  |
| Ordinary one-way Anova | CS0 | CS25 | CS50 | CS75 | Anova test with Welch correction | CS0 | CS25 | CS50 | CS75 | Ordinary one-way Anova | CS0 | CS25 | CS50 | CS75 |
| CS25 | 0.9974 (ns) | X | X | X | CS25 | 0.0308 (*) | X | X | X | CS25 | 0.9998 (ns) | X | X | X |
| CS50 | 0.0325 (*) | 0.0634 (ns) | X | X | CS50 | 0.0292 (*) | 0.9928 (ns) | X | X | CS50 | 0.1467 (ns) | 0.0869 (ns) | X | X |
| CS75 | 0.0056 (**) | 0.0115 (*) | 0.9262 (ns) | X | CS75 | 0.0049 (**) | 0.0019 (**) | 0.0378 (*) | X | CS75 | 0.2296 (ns) | 0.1446 (ns) | 0.9986 (ns) | X |
| CS100 | <0.0001 (****) | 0.0001 (****) | 0.0743 (ns) | 0.3060 (ns) | CS100 | 0.0020 (**) | 0.0171 (*) | 0.0114 (*) | 0.1183 (ns) | CS100 | 0.0003 (****) | 0.0001 (****) | 0.0444 (*) | 0.0261 (*) |

**Table S7:** Statistical comparison of the partitioning coefficient of GK3 (Connected to Figure S18 and Figure 4C)

| Effect of the HSA concentration in each gel type |  |  |  |  |  |  |  |  |  |  |  |  |  |  |
| --- | --- | --- | --- | --- | --- | --- | --- | --- | --- | --- | --- | --- | --- | --- |
| Gel type CS0 |  |  |  |  | Gel type CS25 |  |  |  |  | Gel type CS50 |  |  |  |  |
| Shapiro-Wilk test | PBS | 1mg/ml | 7.5mg/ml | 25mg/ml | Shapiro-Wilk test | PBS | 1mg/ml | 7.5mg/ml | 25mg/ml | Shapiro-Wilk test | PBS | 1mg/ml | 7.5mg/ml | 25mg/ml |
| P value | 0.7888 | 0.6212 | 0.1161 | 0.6803 | P value | 0.5541 | 0.7570 | 0.7002 | 0.3350 | P value | 0.7785 | 0.6499 | 0.4248 | 0.5649 |
| Passed normality test (alpha=0.05)? | Yes | Yes | Yes | Yes | Passed normality test (alpha=0.05)? | Yes | Yes | Yes | Yes | Passed normality test (alpha=0.05)? | Yes | Yes | Yes | Yes |
| P value summary | ns | ns | ns | ns | P value summary | ns | ns | ns | ns | P value summary | ns | ns | ns | ns |
| Brown-Forsythe test |  | Bartlett's test |  |  | Brown-Forsythe test |  | Bartlett's test |  |  | Brown-Forsythe test |  | Bartlett's test |  |  |
| 0.4567 (ns) |  | 0.0844 (ns) |  |  | 0.4113 (ns) |  | 0.3939 (ns) |  |  | 0.2314 (ns) |  | 0.0514 (ns) |  |  |
| Ordinary One-way Anova | PBS | 1mg/ml | 7.5mg/ml |  | Ordinary One-way Anova | PBS | 1mg/ml | 7.5mg/ml |  | Ordinary One-way Anova | PBS | 1mg/ml | 7.5mg/ml |  |
| 1mg/ml | 0.1321 (**) | X | X |  | 1mg/ml | 0.0083 (**) | X | X |  | 1mg/ml | 0.0464 (*) | X | X |  |
| 7.5mg/ml | <0.0001 (****) | 0.0272 (*) | X |  | 7.5mg/ml | <0.0001 (****) | <0.0001 (****) | X |  | 7.5mg/ml | <0.0001 (****) | 0.0103 (*) | X |  |
| 25mg/ml | <0.0001 (****) | 0.0001 (***) | 0.0338 (*) |  | 25mg/ml | <0.0001 (****) | <0.0001 (****) | 0.9792 (ns) |  | 25mg/ml | <0.0001 (****) | 0.0004 (***) | 0.2634 (ns) |  |

| Gel type CS75 |  |  |  |  | Gel type CS100 |  |  |  |  |
| --- | --- | --- | --- | --- | --- | --- | --- | --- | --- |
| Shapiro-Wilk test | PBS | 1mg/ml | 7.5mg/ml | 25mg/ml | Shapiro-Wilk test | PBS | 1mg/ml | 7.5mg/ml | 25mg/ml |
| P value | 0.5004 | 0.4818 | 0.7819 | 0.3549 | P value | 0.7975 | 0.4087 | 0.2981 | 0.6290 |
| Passed normality test (alpha=0.05)? | Yes | Yes | Yes | Yes | Passed normality test (alpha=0.05)? | Yes | Yes | Yes | Yes |
| P value summary | ns | ns | ns | ns | P value summary | ns | ns | ns | ns |
| Brown-Forsythe test |  | Bartlett's test |  |  | Brown-Forsythe test |  | Bartlett's test |  |  |
| 0.1255 (ns) |  | 0.0506 (ns) |  |  | 0.3039 (ns) |  | 0.7568 (ns) |  |  |
| Ordinary One-way Anova | PBS | 1mg/ml | 7.5mg/ml |  | Ordinary One-way Anova | PBS | 1mg/ml | 7.5mg/ml |  |
| 1mg/ml | 0.0013 (**) | X | X |  | 1mg/ml | 0.0052 (**) | X | X |  |
| 7.5mg/ml | <0.0001 (****) | 0.0628 (ns) | X |  | 7.5mg/ml | <0.0001 (****) | 0.0424 (**) | X |  |
| 25mg/ml | <0.0001 (****) | 0.0664 (*) | >0.9999 (ns) |  | 25mg/ml | <0.0001 (****) | 0.0001 (****) | 0.0284 (**) |  |

| 1mg/ml |  |  |  |  | 7.5mg/ml |  |  |  |  | 25mg/ml |  |  |  |  |
| --- | --- | --- | --- | --- | --- | --- | --- | --- | --- | --- | --- | --- | --- | --- |
| Brown-Forsythe test |  | Bartlett's test |  |  | Brown-Forsythe test |  | Bartlett's test |  |  | Brown-Forsythe test |  | Bartlett's test |  |  |
| 0.0309 (*) |  | 0.2305 (ns) |  |  | 0.3734 (ns) |  | 0.3061 (ns) |  |  | 0.0231 (*) |  | 0.0437 (*) |  |  |
| Anova test with Welch correction | CS0 | CS25 | CS50 | CS75 | Ordinary one-way Anova | CS0 | CS25 | CS50 | CS75 | Anova test with Welch correction | CS0 | CS25 | CS50 | CS75 |
| CS25 | 0.9989 (ns) | X | X | X | CS25 | 0.8596 (ns) | X | X | X | CS25 | 0.0128 (*) | X | X | X |
| CS50 | 0.9598 (ns) | 0.8668 (ns) | X | X | CS50 | 0.9559 (ns) | 0.9982 (ns) | X | X | CS50 | 0.3312 (ns) | 0.0407 (*) | X | X |
| CS75 | >0.9999 (ns) | 0.9990 (ns) | 0.9590 (ns) | X | CS75 | >0.9999 (ns) | 0.8406 (ns) | 0.9457 (ns) | X | CS75 | 0.0011 (**) | 0.0446 (*) | 0.0025 (**) | X |
| CS100 | 0.1491 (ns) | 0.1839 (ns) | 0.0397 (*) | 0.1501 (ns) | CS100 | 0.2250 (ns) | 0.0404 (*) | 0.0696 (ns) | 0.2403 (ns) | CS100 | 0.3798 (ns) | 0.9999 (ns) | 0.7372 (ns) | 0.6274 (ns) |

**Table S8:** Statistical comparison of the partitioning coefficient of RM26 (Connected to Figure S18 and Figure 4C)

| Effect of the HSA concentration in each gel type |  |  |  |  |  |  |  |  |  |  |  |  |  |  |
| --- | --- | --- | --- | --- | --- | --- | --- | --- | --- | --- | --- | --- | --- | --- |
| N<sub>g</sub>=2 N<sub>m</sub>=2/2 |  | Gel type CS0 |  |  |  | Gel type CS25 |  |  |  | Gel type CS50 |  |  |  |  |
| Shapiro-Wilk test | PBS | 1mg/ml | 7.5mg/ml | 25mg/ml | Shapiro-Wilk test | PBS | 1mg/ml | 7.5mg/ml | 25mg/ml | Shapiro-Wilk test | PBS | 1mg/ml | 7.5mg/ml | 25mg/ml |
| P value | 0.0866 | 0.7379 | 0.7898 | 0.1510 | P value | 0.7708 | 0.7835 | 0.9755 | 0.0751 | P value | 0.9753 | 0.6928 | 0.1458 | 0.1029 |
| Passed normality test (alpha=0.05)? | Yes | Yes | Yes | Yes | Passed normality test (alpha=0.05)? | Yes | Yes | Yes | Yes | Passed normality test (alpha=0.05)? | Yes | Yes | Yes | Yes |
| P value summary | ns | ns | ns | ns | P value summary | ns | ns | ns | ns | P value summary | ns | ns | ns | ns |
| Brown-Forsythe test |  | Bartlett's test |  |  | Brown-Forsythe test |  | Bartlett's test |  |  | Brown-Forsythe test |  | Bartlett's test |  |  |
| 0.2876 (ns) |  | 0.0031 (**) |  |  | 0.8879 (ns) |  | 0.7539 (ns) |  |  | 0.7623 (ns) |  | 0.8387 (ns) |  |  |
| Anova test with Welch correction | PBS | 1mg/ml | 7.5mg/ml |  | Ordinary One-way Anova | PBS | 1mg/ml | 7.5mg/ml |  | Ordinary One-way Anova | PBS | 1mg/ml | 7.5mg/ml |  |
| 1mg/ml | 0.9945 (ns) | X | X |  | 1mg/ml | 0.9950 (ns) | X | X |  | 1mg/ml | 0.0652 (ns) | X | X |  |
| 7.5mg/ml | 0.2229 (ns) | 0.0051 (**) | X |  | 7.5mg/ml | 0.04959 (*) | 0.0303 (*) | X |  | 7.5mg/ml | 0.0069 (**) | 0.5859 (**) | X |  |
| 25mg/ml | 0.1393 (ns) | 0.0067 (**) | 0.3759 (ns) |  | 25mg/ml | 0.0024 (**) | 0.0016 (**) | 0.3587 (ns) |  | 25mg/ml | <0.0001 (****) | 0.0039 (**) | 0.0364 (*) |  |

| Gel type CS75 |  |  |  |  | Gel type CS100 |  |  |  |  |
| --- | --- | --- | --- | --- | --- | --- | --- | --- | --- |
| Shapiro-Wilk test | PBS | 1mg/ml | 7.5mg/ml | 25mg/ml | Shapiro-Wilk test | PBS | 1mg/ml | 7.5mg/ml | 25mg/ml |
| P value | 0.3292 | 0.9321 | 0.7643 | 0.8832 | P value | 0.7925 | 0.7710 | 0.7395 | 0.0589 |
| Passed normality test (alpha=0.05)? | Yes | Yes | Yes | Yes | Passed normality test (alpha=0.05)? | Yes | Yes | Yes | Yes |
| P value summary | ns | ns | ns | ns | P value summary | ns | ns | ns | ns |
| Brown-Forsythe test |  | Bartlett's test |  |  | Brown-Forsythe test |  | Bartlett's test |  |  |
| 0.4642 (ns) |  | 0.1652 (ns) |  |  | 0.6390 (ns) |  | 0.0348 (*) |  |  |
| Ordinary One-way Anova | PBS | 1mg/ml | 7.5mg/ml |  | Anova test with Welch correction | PBS | 1mg/ml | 7.5mg/ml |  |
| 1mg/ml | 0.8949 (ns) | X | X |  | 1mg/ml | 0.5066 (ns) | X | X |  |
| 7.5mg/ml | 0.3247 (ns) | 0.1124 (ns) | X |  | 7.5mg/ml | 0.9734 (ns) | 0.3509 (ns) | X |  |
| 25mg/ml | 0.0030 (**) | 0.0009 (***) | 0.0654 (ns) |  | 25mg/ml | 0.0016 (**) | 0.0030 (**) | 0.0031 (**) |  |

| 1mg/ml |  |  |  |  | 7.5mg/ml |  |  |  |  | 25mg/ml |  |  |  |  |
| --- | --- | --- | --- | --- | --- | --- | --- | --- | --- | --- | --- | --- | --- | --- |
| Brown-Forsythe test |  | Bartlett's test |  |  | Brown-Forsythe test |  | Bartlett's test |  |  | Brown-Forsythe test |  | Bartlett's test |  |  |
| 0.2982 (ns) |  | 0.0094 (**) |  |  | 0.5321 (ns) |  | 0.2199 (ns) |  |  | 0.4956 (ns) |  | 0.0242 (*) |  |  |
| Anova test with Welch correction | CS0 | CS25 | CS50 | CS75 | Ordinary one-way Anova | CS0 | CS25 | CS50 | CS75 | Anova test with Welch correction | CS0 | CS25 | CS50 | CS75 |
| CS25 | 0.8969 (ns) | X | X | X | CS25 | 0.7594 (ns) | X | X | X | CS25 | 0.9903 (ns) | X | X | X |
| CS50 | 0.0465 (*) | 0.1905 (ns) | X | X | CS50 | 0.6608 (ns) | 0.9988 (ns) | X | X | CS50 | 0.8791 (ns) | >0.9999 (ns) | X | X |
| CS75 | 0.5993 (ns) | 0.9746 (ns) | 0.3600 (ns) | X | CS75 | 0.3838 (ns) | 0.9612 (ns) | 0.9871 (ns) | X | CS75 | 0.8637 (ns) | 0.9999 (ns) | 0.9996 (ns) | X |
| CS100 | 0.6950 (ns) | >0.9999 (ns) | 0.0180 (*) | 0.9116 (ns) | CS100 | 0.0026 (**) | 0.0253 (*) | 0.0352 (*) | 0.0871 (ns) | CS100 | 0.2752 (ns) | 0.9770 (ns) | 0.7900 (ns) | 0.5939 (ns) |

**Table S9:** Statistical comparison of the partitioning coefficient of KWEA (Connected to Figure S18 and Figure 4C)

| Effect of the HSA concentration in each gel type |  |  |  |  |  |  |  |  |  |  |  |  |  |  |
| --- | --- | --- | --- | --- | --- | --- | --- | --- | --- | --- | --- | --- | --- | --- |
| Gel type CS0 |  |  |  |  | Gel type CS25 |  |  |  |  | Gel type CS50 |  |  |  |  |
| Shapiro-Wilk test | PBS | 1mg/ml | 7.5mg/ml | 25mg/ml | Shapiro-Wilk test | PBS | 1mg/ml | 7.5mg/ml | 25mg/ml | Shapiro-Wilk test | PBS | 1mg/ml | 7.5mg/ml | 25mg/ml |
| P value | 0.7440 | 0.7985 | 0.7981 | 0.7960 | P value | 0.3395 | 0.6734 | 0.6944 | 0.7924 | P value | 0.7799 | 0.7491 | 0.3666 | 0.1642 |
| Passed normality test (alpha=0.05)? | Yes | Yes | Yes | Yes | Passed normality test (alpha=0.05)? | Yes | Yes | Yes | Yes | Passed normality test (alpha=0.05)? | Yes | Yes | Yes | Yes |
| P value summary | ns | ns | ns | ns | P value summary | ns | ns | ns | ns | P value summary | ns | ns | ns | ns |
| Brown-Forsythe test |  | Bartlett's test |  |  | Brown-Forsythe test |  | Bartlett's test |  |  | Brown-Forsythe test |  | Bartlett's test |  |  |
| 0.2758 (ns) |  | 0.0044 (**) |  |  | 0.5059 (ns) |  | 0.3448 (ns) |  |  | 0.4698 (ns) |  | 0.0688 (ns) |  |  |
| Anova test with Welch correction | PBS | 1mg/ml | 7.5mg/ml |  | Ordinary One-way Anova | PBS | 1mg/ml | 7.5mg/ml |  | Ordinary One-way Anova | PBS | 1mg/ml | 7.5mg/ml |  |
| 1mg/ml | 0.4789 (ns) | X | X |  | 1mg/ml | 0.9367 (ns) | X | X |  | 1mg/ml | 0.8766 (ns) | X | X |  |
| 7.5mg/ml | 0.4032 (ns) | 0.8809 (ns) | X |  | 7.5mg/ml | 0.9962 (ns) | 0.9829 (ns) | X |  | 7.5mg/ml | 0.1999 (ns) | 0.5342 (ns) | X |  |
| 25mg/ml | 0.0278 (*) | 0.0254 (*) | 0.1648 (ns) |  | 25mg/ml | 0.9989 (ns) | 0.9714 (ns) | 0.9999 (ns) |  | 25mg/ml | 0.0361 (*) | 0.1279 (ns) | 0.7357 (ns) |  |
| Gel type CS75 |  |  |  |  | Gel type CS100 |  |  |  |  |  |  |  |  |  |
| Shapiro-Wilk test | PBS | 1mg/ml | 7.5mg/ml | 25mg/ml | Shapiro-Wilk test | PBS | 1mg/ml | 7.5mg/ml | 25mg/ml |  |  |  |  |  |
| P value | 0.6038 | 0.6030 | 0.7589 | 0.7963 | P value | 0.0284 | 0.4703 | 0.6447 | 0.2496 |  |  |  |  |  |
| Passed normality test (alpha=0.05)? | Yes | Yes | Yes | Yes | Passed normality test (alpha=0.05)? | Yes | Yes | Yes | Yes |  |  |  |  |  |
| P value summary | ns | ns | ns | ns | P value summary | ns | ns | ns | ns |  |  |  |  |  |
| Brown-Forsythe test |  | Bartlett's test |  |  | Kruskal-Wallis test |  | Exact P value |  |  |  |  |  |  |  |
| 0.3972 (ns) |  | 0.1404 (ns) |  |  | 0.0768 (ns) |  | - |  |  |  |  |  |  |  |
| Ordinary One-way Anova | PBS | 1mg/ml | 7.5mg/ml |  | Ordinary One-way Anova | PBS | 1mg/ml | 7.5mg/ml |  |  |  |  |  |  |
| 1mg/ml | 0.8652 (ns) | X | X |  | 1mg/ml | 0.1876 (ns) | X | X |  |  |  |  |  |  |
| 7.5mg/ml | 0.2263 (ns) | 0.0650 (ns) | X |  | 7.5mg/ml | >0.9999 (ns) | 0.3211 (ns) | X |  |  |  |  |  |  |
| 25mg/ml | 0.9246 (ns) | 0.5283 (ns) | 0.5080 (ns) |  | 25mg/ml | >0.9999 (ns) | 0.1876 (ns) | >0.9999 (ns) |  |  |  |  |  |  |
| Effect of the gel types at each HSA concentration |  |  |  |  |  |  |  |  |  |  |  |  |  |  |
| 1mg/ml |  |  |  |  | 7.5mg/ml |  |  |  |  | 25mg/ml |  |  |  |  |
| Brown-Forsythe test |  | Bartlett's test |  |  | Brown-Forsythe test |  | Bartlett's test |  |  | Brown-Forsythe test |  | Bartlett's test |  |  |
| 0.1554 (ns) |  | 0.0053 (**) |  |  | 0.3686 (ns) |  | 0.1469 (ns) |  |  | 0.3131 (ns) |  | 0.0383 (*) |  |  |
| Anova test with Welch correction | CS0 | CS25 | CS50 | CS75 | Ordinary one-way Anova | CS0 | CS25 | CS50 | CS75 | Anova test with Welch correction | CS0 | CS25 | CS50 | CS75 |
| CS25 | 0.9991 (ns) | X | X | X | CS25 | 0.9984 (ns) | X | X | X | CS25 | 0.0939 (ns) | X | X | X |
| CS50 | 0.8813 (ns) | 0.9304 (ns) | X | X | CS50 | 0.2502 (ns) | 0.1590 (ns) | X | X | CS50 | 0.9652 (ns) | 0.0678 (ns) | X | X |
| CS75 | 0.4299 (ns) | 0.5075 (ns) | 0.5316 (ns) | X | CS75 | 0.44714 (ns) | 0.3258 (ns) | 0.9893 (ns) | X | CS75 | 0.2611 (ns) | >0.9999 (ns) | 0.1926 (ns) | X |
| CS100 | 0.2646 (ns) | 0.2542 (ns) | 0.2682 (ns) | 0.7632 (ns) | CS100 | 0.9850 (ns) | 0.9279 (ns) | 0.4963 (ns) | 0.7655 (ns) | CS100 | 0.5121 (ns) | 0.9999 (ns) | 0.4044 (ns) | 0.9999 (ns) |

**Table S10:** Statistical comparison of the partitioning coefficient of Exe (Connected to Figure S18 and Figure 4C)

| Effect of the HSA concentration in each gel type |  |  |  |  |  |  |  |  |  |  |  |  |  |  |
| --- | --- | --- | --- | --- | --- | --- | --- | --- | --- | --- | --- | --- | --- | --- |
| Gel type CS0 |  |  |  |  | Gel type CS25 |  |  |  |  | Gel type CS50 |  |  |  |  |
| Shapiro-Wilk test | PBS | 1mg/ml | 7.5mg/ml | 25mg/ml | Shapiro-Wilk test | PBS | 1mg/ml | 7.5mg/ml | 25mg/ml | Shapiro-Wilk test | PBS | 1mg/ml | 7.5mg/ml | 25mg/ml |
| P value | 0.5630 | 0.4082 | 0.7157 | 0.1105 | P value | 0.7067 | 0.3693 | 0.6430 | 0.1930 | P value | 0.1029 | 0.6484 | 0.3861 | 0.5290 |
| Passed normality test (alpha=0.05)? | Yes | Yes | Yes | Yes | Passed normality test (alpha=0.05)? | Yes | Yes | Yes | Yes | Passed normality test (alpha=0.05)? | Yes | Yes | Yes | Yes |
| P value summary | ns | ns | ns | ns | P value summary | ns | ns | ns | ns | P value summary | ns | ns | ns | ns |
| Brown-Forsythe test |  | Bartlett's test |  |  | Brown-Forsythe test |  | Bartlett's test |  |  | Brown-Forsythe test |  | Bartlett's test |  |  |
| 0.1512 (ns) |  | 0.5215 (ns) |  |  | 0.4992 (ns) |  | 0.1312 (ns) |  |  | 0.5123 (ns) |  | 0.8826 (ns) |  |  |
| Ordinary One-way Anova | PBS | 1mg/ml | 7.5mg/ml |  | Ordinary One-way Anova | PBS | 1mg/ml | 7.5mg/ml |  | Ordinary One-way Anova | PBS | 1mg/ml | 7.5mg/ml |  |
| 1mg/ml | 0.3998 (ns) | X | X |  | 1mg/ml | 0.9996 (ns) | X | X |  | 1mg/ml | 0.8395 (ns) | X | X |  |
| 7.5mg/ml | 0.3871 (ns) | >0.9999 (ns) | X |  | 7.5mg/ml | 0.9972 (ns) | 0.9997 (ns) | X |  | 7.5mg/ml | 0.0320 (*) | 0.1309 (ns) | X |  |
| 25mg/ml | 0.0227 (*) | 0.3228 (ns) | 0.3341 (ns) |  | 25mg/ml | 0.9631 (ns) | 0.9814 (ns) | 0.9919 (ns) |  | 25mg/ml | 0.1784 (ns) | 0.5394 (ns) | 0.7386 (ns) |  |

| Gel type CS75 |  |  |  |  | Gel type CS100 |  |  |  |  |
| --- | --- | --- | --- | --- | --- | --- | --- | --- | --- |
| Shapiro-Wilk test | PBS | 1mg/ml | 7.5mg/ml | 25mg/ml | Shapiro-Wilk test | PBS | 1mg/ml | 7.5mg/ml | 25mg/ml |
| P value | 0.8639 | 0.7533 | 0.9378 | 0.7879 | P value | 0.6316 | 0.6077 | 0.5066 | 0.6845 |
| Passed normality test (alpha=0.05)? | Yes | Yes | Yes | Yes | Passed normality test (alpha=0.05)? | Yes | Yes | Yes | Yes |
| P value summary | ns | ns | ns | ns | P value summary | ns | ns | ns | ns |
| Brown-Forsythe test |  | Bartlett's test |  |  | Brown-Forsythe test |  | Bartlett's test |  |  |
| 0.0004 (**) |  | 0.3434 (ns) |  |  | 0.2451 (ns) |  | 0.1990 (ns) |  |  |
| Anova test with Welch correction | PBS | 1mg/ml | 7.5mg/ml |  | Ordinary One-way Anova | PBS | 1mg/ml | 7.5mg/ml |  |
| 1mg/ml | 0.2492 (ns) | X | X |  | 1mg/ml | 0.0680 (ns) | X | X |  |
| 7.5mg/ml | 0.0011 (**) | 0.0043 (**) | X |  | 7.5mg/ml | 0.0794 (ns) | 0.9997 (ns) | X |  |
| 25mg/ml | 0.5290 (ns) | 0.9848 (ns) | 0.0964 (ns) |  | 25mg/ml | 0.1080 (ns) | 0.9923 (ns) | 0.9976 (ns) |  |

| Effect of the gel types at each HSA concentration |  |  |  |  |  |  |  |  |  |  |  |  |  |  |
| --- | --- | --- | --- | --- | --- | --- | --- | --- | --- | --- | --- | --- | --- | --- |
| 1mg/ml |  |  |  |  | 7.5mg/ml |  |  |  |  | 25mg/ml |  |  |  |  |
| Brown-Forsythe test |  | Bartlett's test |  |  | Brown-Forsythe test |  | Bartlett's test |  |  | Brown-Forsythe test |  | Bartlett's test |  |  |
| 0.3449 (ns) |  | 0.7140 (ns) |  |  | 0.6268 (ns) |  | 0.2707 (ns) |  |  | 0.8573 (ns) |  | 0.9315 (ns) |  |  |
| Ordinary one-way Anova | CS0 | CS25 | CS50 | CS75 | Ordinary one-way Anova | CS0 | CS25 | CS50 | CS75 | Ordinary one-way Anova | CS0 | CS25 | CS50 | CS75 |
| CS25 | 0.9962 (ns) | X | X | X | CS25 | 0.9900 (ns) | X | X | X | CS25 | 0.9119 (ns) | X | X | X |
| CS50 | 0.7982 (ns) | 0.5972 (ns) | X | X | CS50 | 0.3912 (ns) | 0.6506 (ns) | X | X | CS50 | 0.8568 (ns) | 0.9999 (ns) | X | X |
| CS75 | 0.1507 (ns) | 0.0811 (ns) | 0.6622 (ns) | X | CS75 | 0.0128 (*) | 0.0308 (*) | 0.3266 (ns) | X | CS75 | 0.1916 (ns) | 0.5920 (ns) | 0.6755 (ns) | X |
| CS100 | 0.9994 (ns) | >0.9999 (ns) | 0.6752 (ns) | 0.1027 (ns) | CS100 | >0.9999 (ns) | 0.9948 (*) | 0.4287 (*) | 0.0147 (*) | CS100 | 0.6609 (ns) | 0.9847 (ns) | 0.9955 (ns) | 0.8675 (ns) |

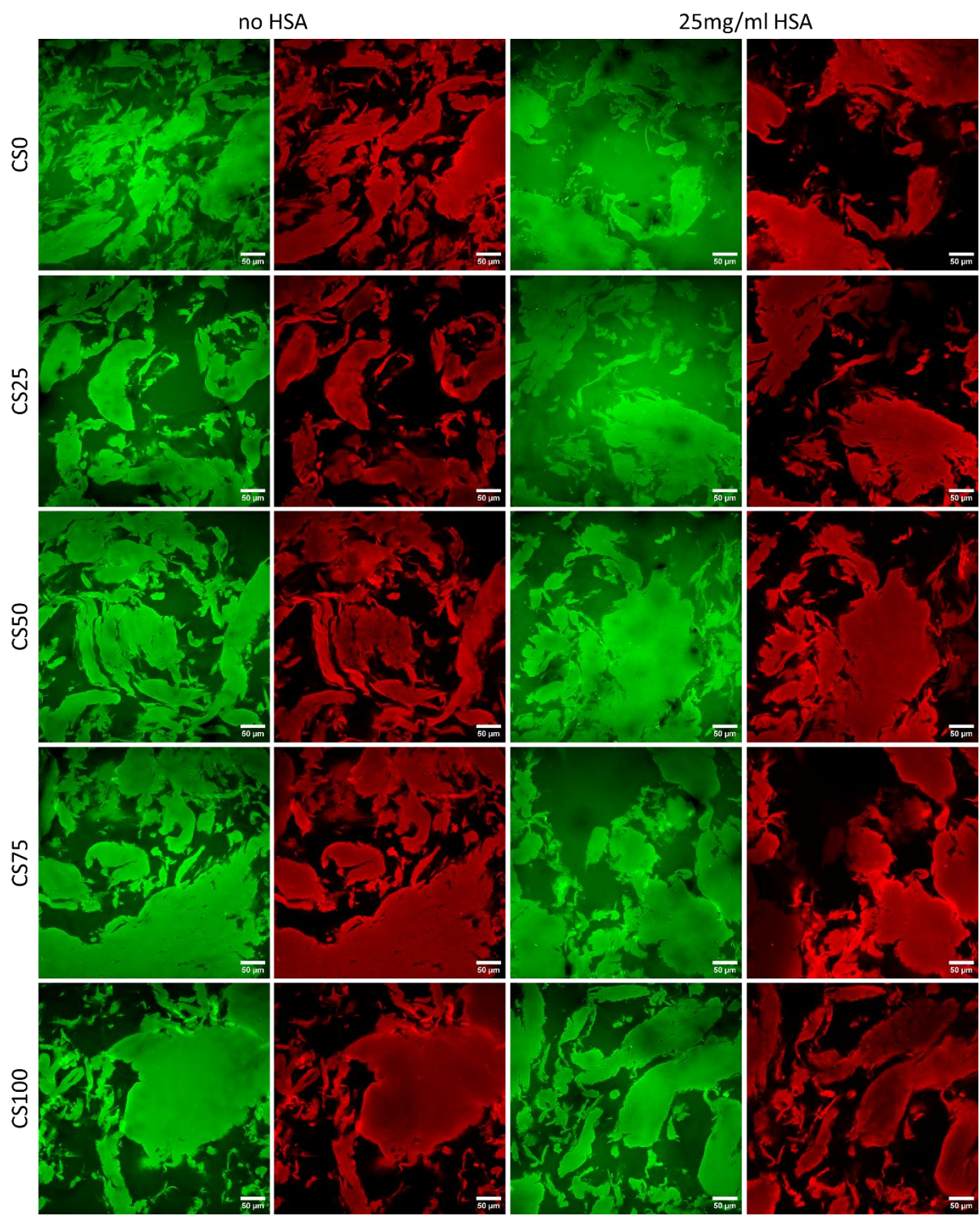

**Figure S19:** Effect of 25mg/ml HSA on the distribution of GK6 in each hydrogel type

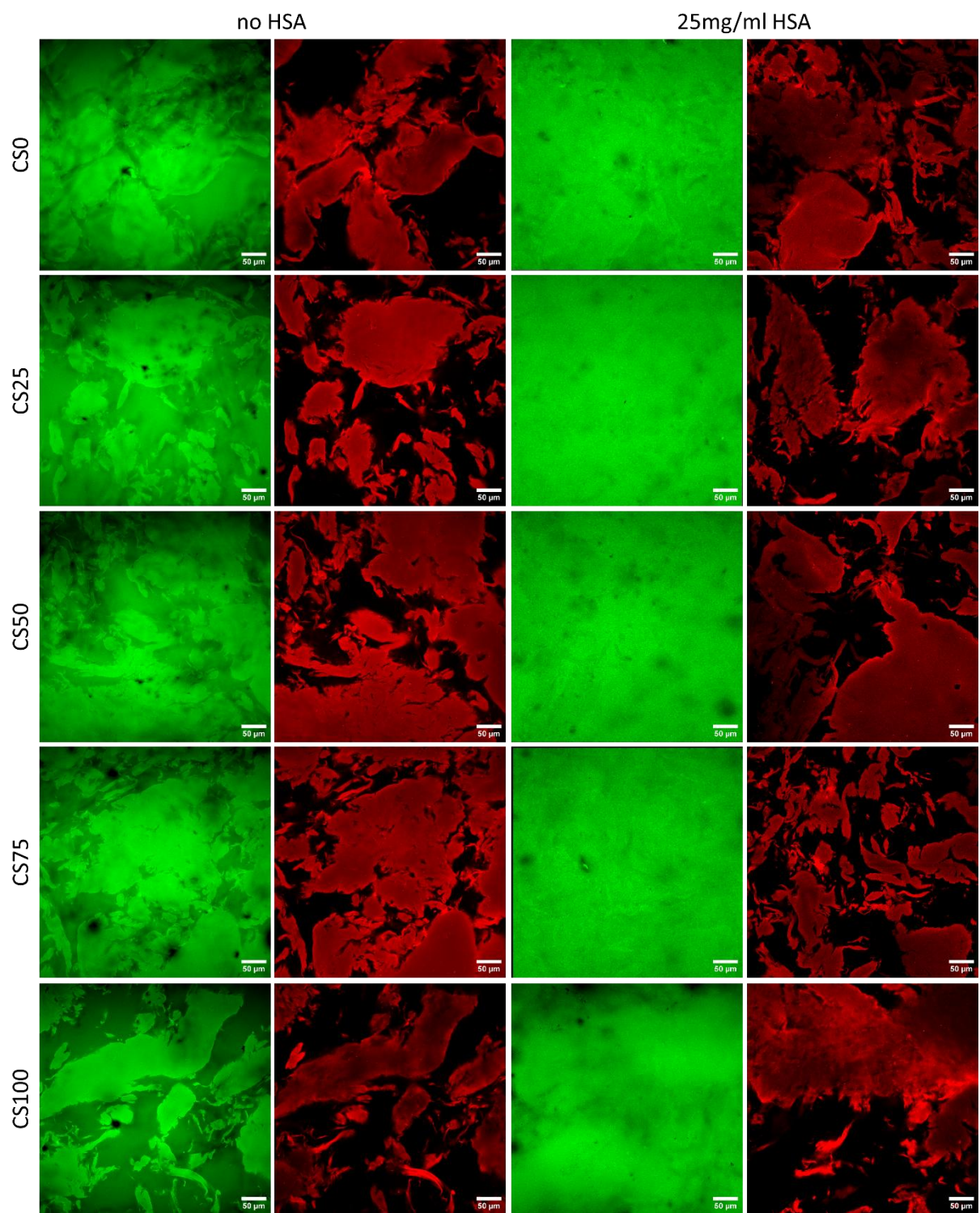

**Figure S20:** Effect of 25mg/ml HSA on the distribution of GK3 in each hydrogel type

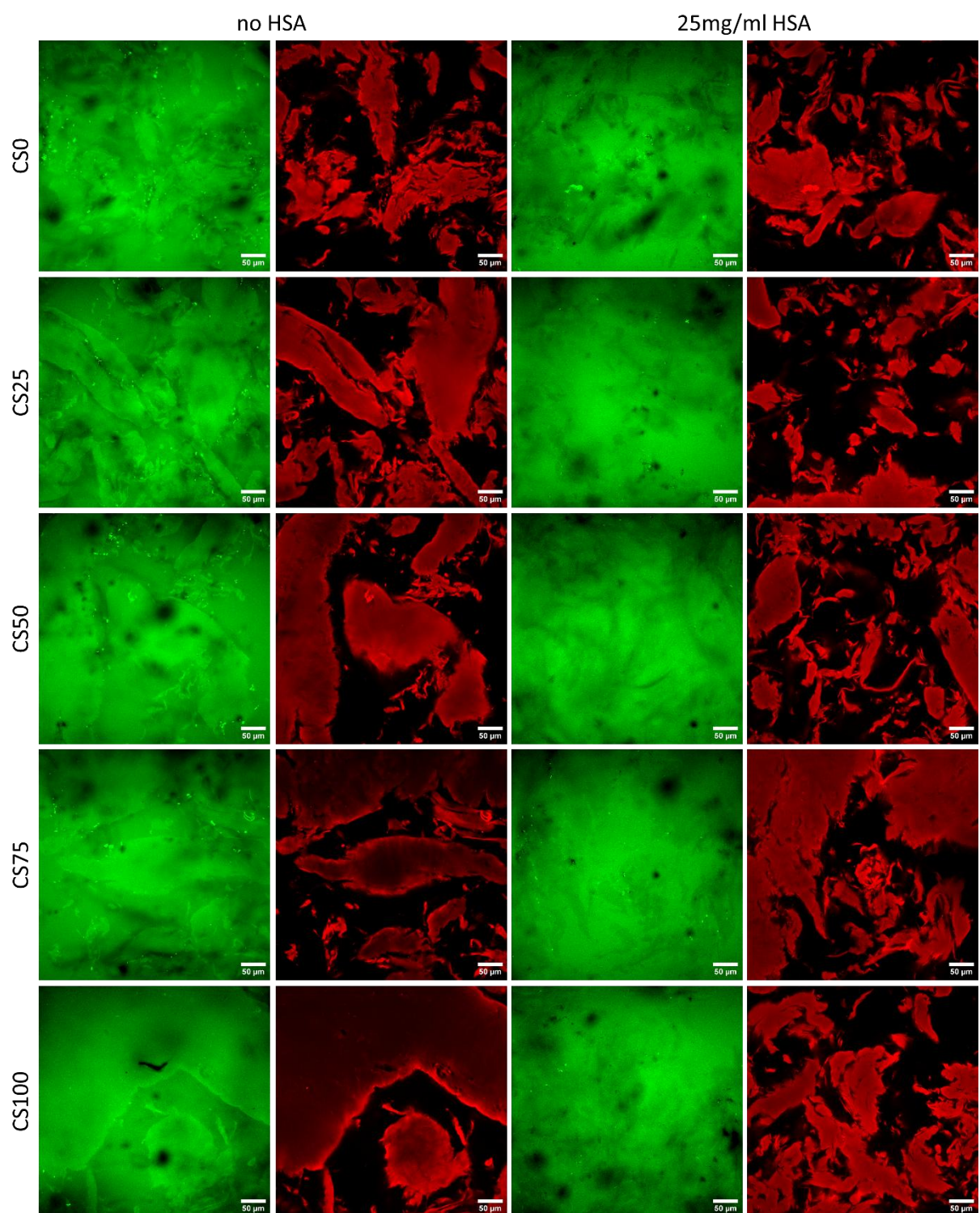

**Figure S21:** Effect of 25mg/ml HSA on the distribution of RM26 in each hydrogel type

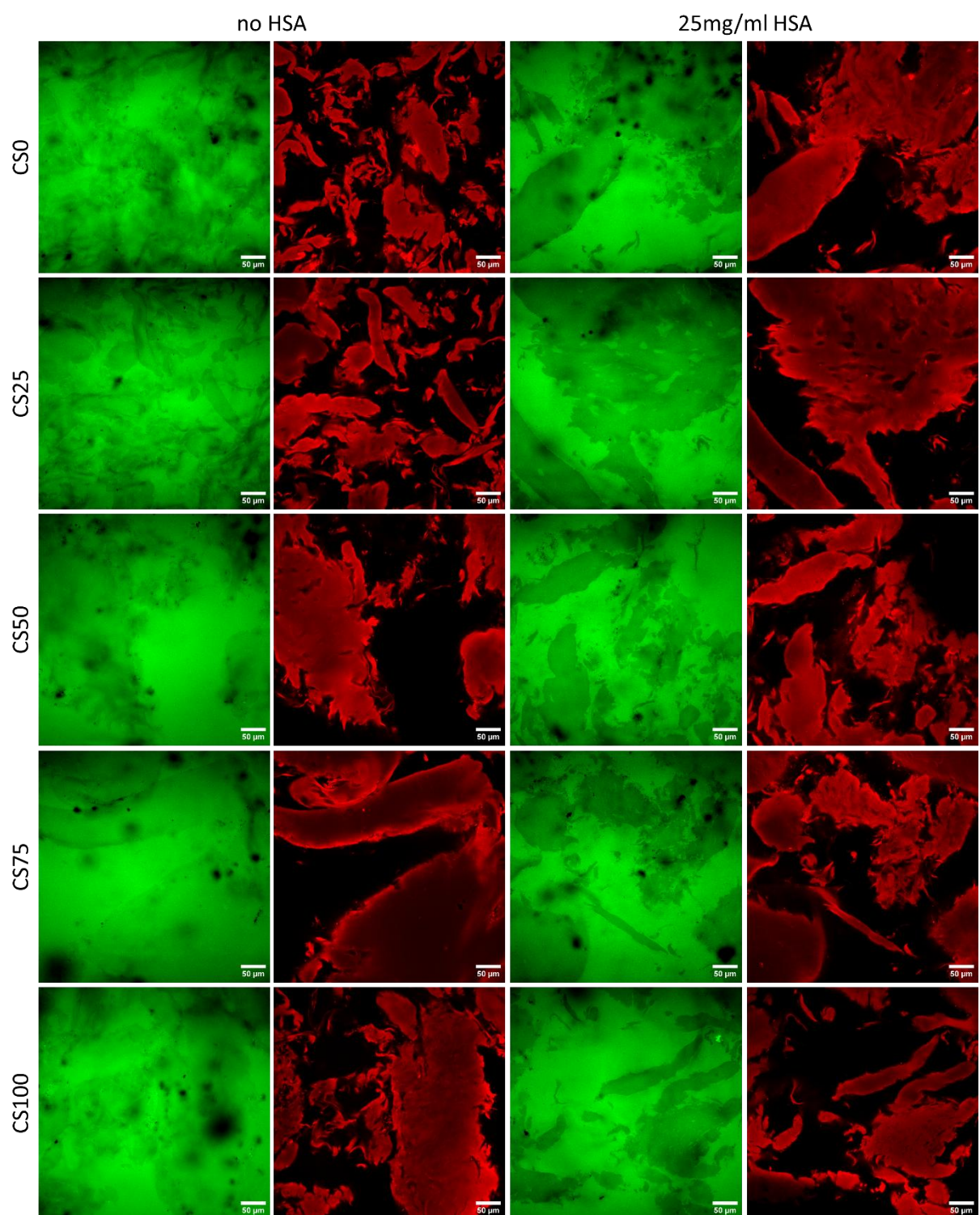

**Figure S22:** Effect of 25mg/ml HSA on the distribution of KWEA in each hydrogel type

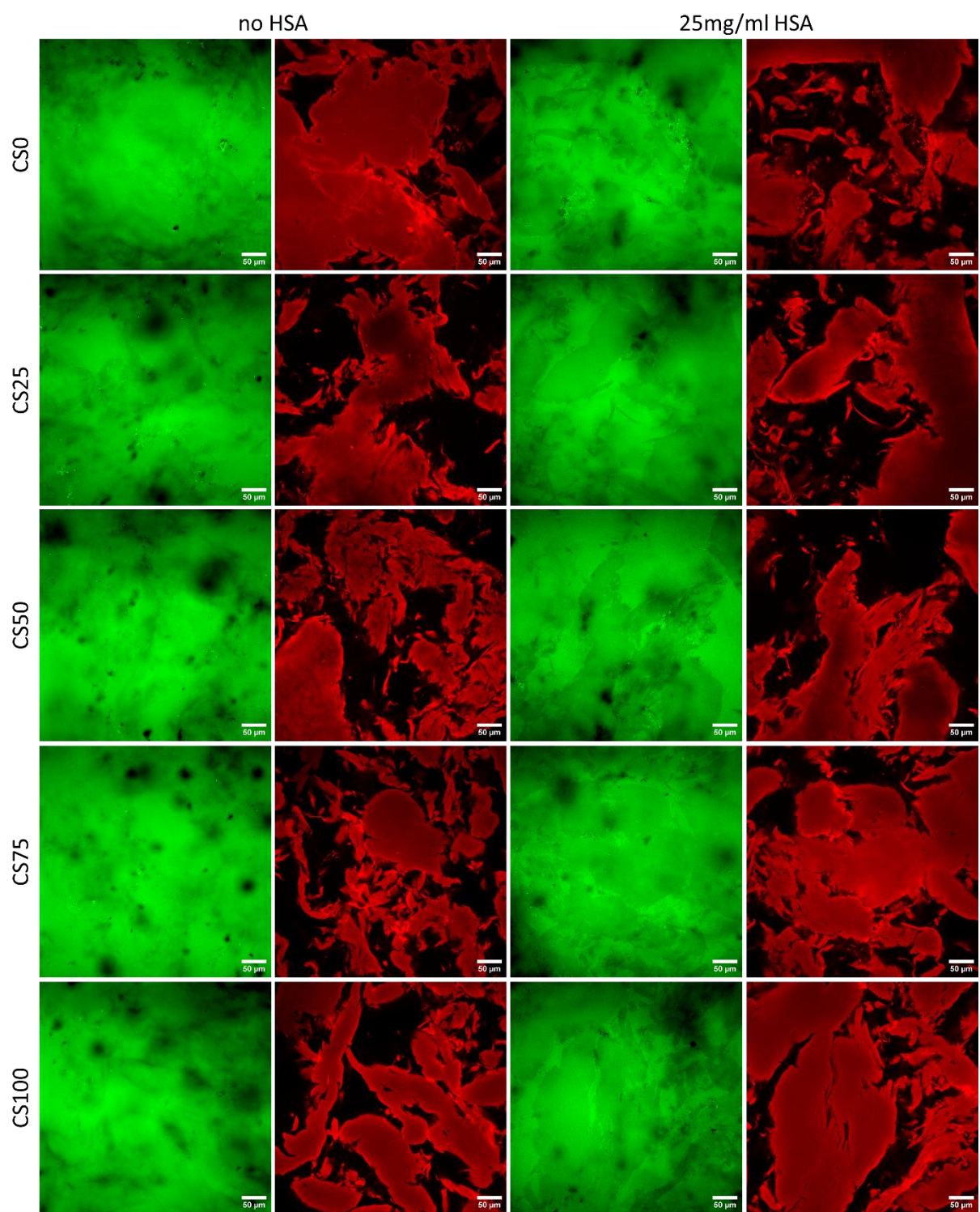

**Figure S23:** Effect of 25mg/ml HSA on the distribution of Exe in each hydrogel type

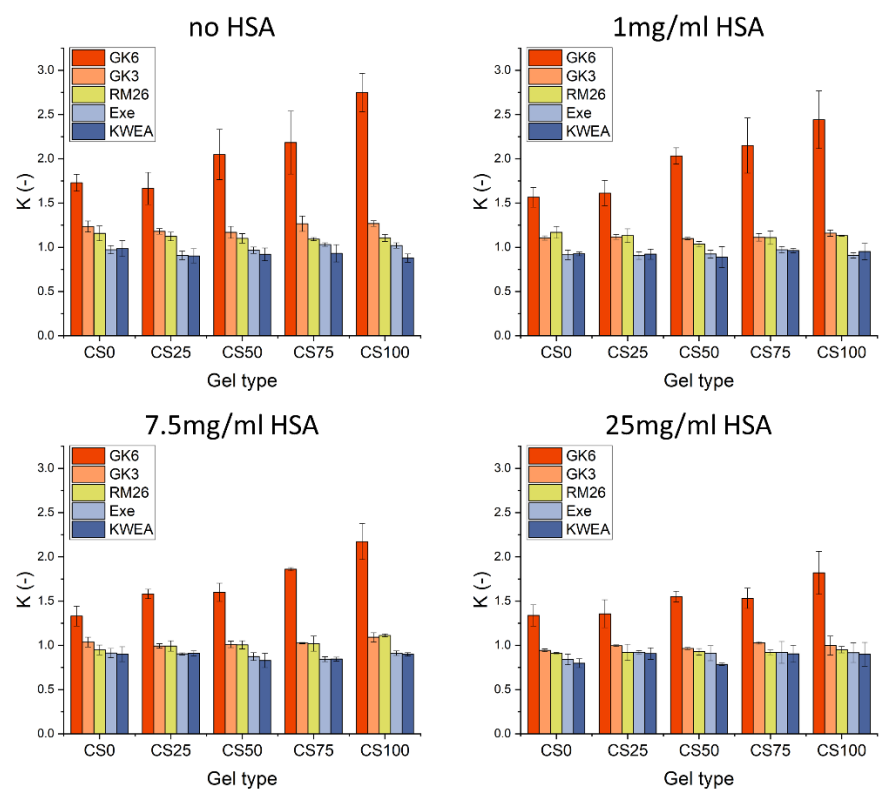

**Figure S24:** Partitioning coefficient of the different peptides in different hydrogels at each HSA concentration

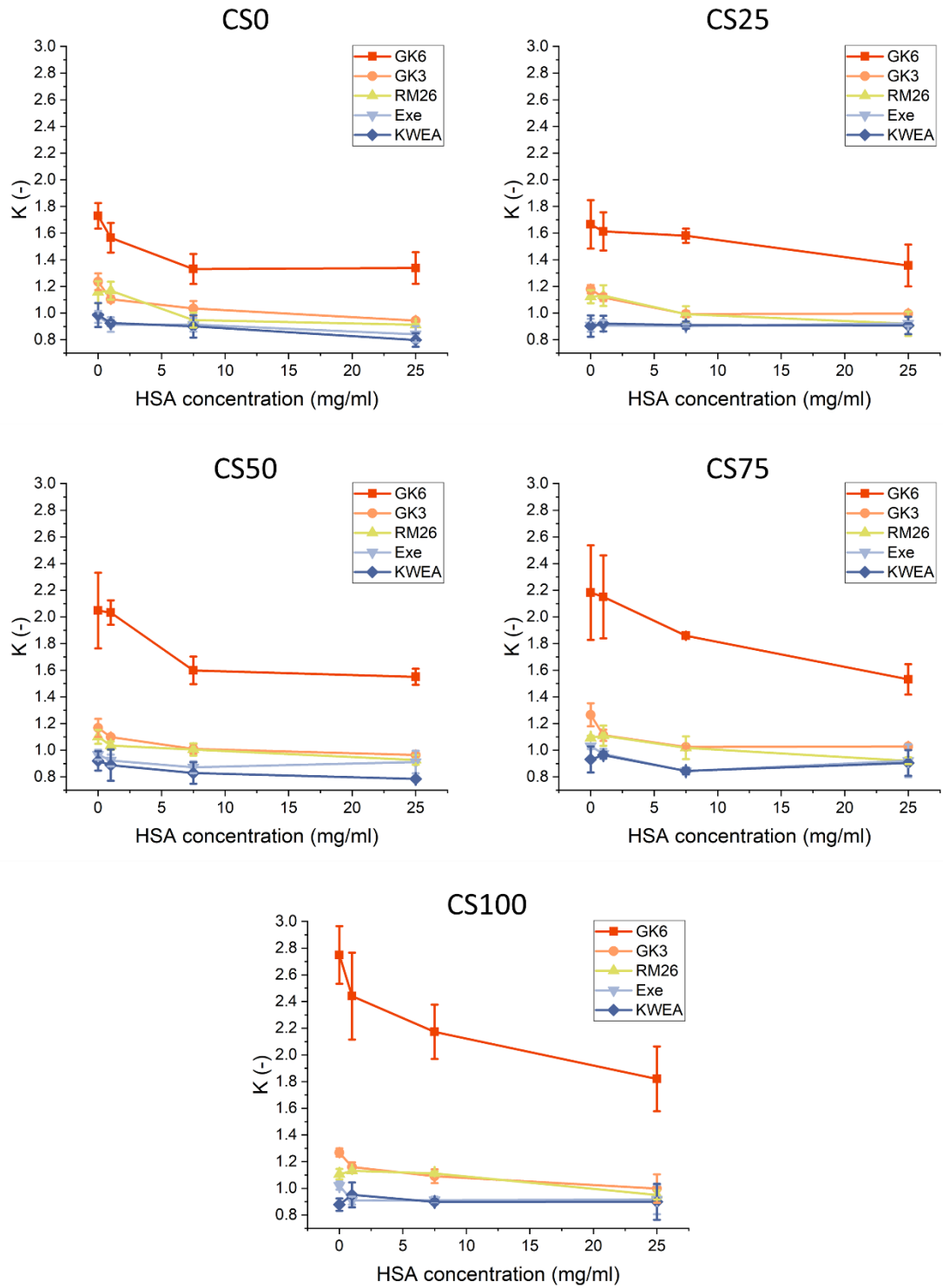

**Figure S25:** HSA concentration-dependent partitioning of the peptides in each hydrogel

### Effect of incubation time on the partitioning coefficient of different peptides

HSA concentrations of 7.5 and 25 mg/mL were used for peptides whose partitioning coefficient showed strong dependence on HSA concentration (GK6 and GK3), whereas 25 mg/mL was used for peptides whose partitioning coefficient showed weak or no dependence (RM26 and Exe). Because hydrogel composition significantly affected only GK6 distribution, the number of hydrogel types was reduced for the time-dependent measurements for other peptides.

**Table S11:** Statistical comparison of the partitioning coefficient of GK6 among hydrogel types after 21 hours of incubation (Related to Figure 4E)

| GK6 partitioning kinetics |  |  |  |  |  | GK6 partitioning kinetics |  |  |  |  |  |
| --- | --- | --- | --- | --- | --- | --- | --- | --- | --- | --- | --- |
| Different gel types after 21h. 7.5mg/ml HSA |  |  |  |  |  | Different gel types after 21h. 25mg/ml HSA |  |  |  |  |  |
| Shapiro-Wilk test | CS0 | CS25 | CS50 | CS75 | CS100 | Shapiro-Wilk test | CS0 | CS25 | CS50 | CS75 | CS100 |
| P value | 0.5578 | 0.7742 | 0.7737 | 0.7650 | 0.7903 | P value | 0.5278 | 0.1966 | 0.3052 | 0.7822 | 0.7891 |
| Passed normality test (alpha=0.05)? | Yes | Yes | Yes | Yes | Yes | Passed normality test (alpha=0.05)? | Yes | Yes | Yes | Yes | Yes |
| P value summary | ns | ns | ns | ns | ns | P value summary | ns | ns | ns | ns | ns |
| Brown-Forsythe test |  |  | Bartlett's test |  |  | Brown-Forsythe test |  |  | Bartlett's test |  |  |
| 0.1671 (ns) |  |  | 0.4539 (ns) |  |  | 0.2110 (ns) |  |  | 0.0302 (*) |  |  |
| Ordinary One-way Anova | CS0 | CS25 | CS50 | CS75 |  | Anova test with Welch correction | CS0 | CS25 | CS50 | CS75 |  |
| CS25 | 0.3370 (ns) | X | X | X |  | CS25 | 0.5964 (ns) | X | X | X |  |
| CS50 | 0.0060 (**) | 0.2152 (ns) | X | X |  | CS50 | 0.9212 (ns) | 0.9638 (ns) | X | X |  |
| CS75 | 0.0200 (*) | 0.5056 (ns) | 0.9687 (ns) | X |  | CS75 | 0.0594 (ns) | 0.5619 (ns) | 0.2401 (ns) | X |  |
| CS100 | 0.0016 (**) | 0.0645 (ns) | 0.9530 (ns) | 0.6802 (ns) |  | CS100 | 0.2222 (ns) | 0.9383 (ns) | 0.6307 (ns) | 0.9361 (ns) |  |

**Table S12:** Statistical comparison of the partitioning coefficient of GK3, RM26, and Exe among hydrogel types after 21 hours of incubation (Related to Figure 4E)

| GK3 |  |  |  | RM26 |  |  |  | Exe |  |  |  |
| --- | --- | --- | --- | --- | --- | --- | --- | --- | --- | --- | --- |
| Different gel types after 21h. 7.5mg/ml HSA |  |  |  | Different gel types after 21h. 25mg/ml HSA |  |  |  | Different gel types after 21h. 25mg/ml HSA |  |  |  |
| Shapiro-Wilk test | CS0 | CS50 | CS100 | Shapiro-Wilk test | CS0 | CS50 | CS100 | Shapiro-Wilk test | CS0 | CS50 | CS100 |
| P value | 0.6308 | 0.7286 | 0.7881 | P value | 0.5819 | 0.3914 | 0.7424 | P value | 0.2308 | 0.6732 | 0.7876 |
| Passed normality test (alpha=0.05)? | Yes | Yes | Yes | Passed normality test (alpha=0.05)? | Yes | Yes | Yes | Passed normality test (alpha=0.05)? | Yes | Yes | Yes |
| P value summary | ns | ns | ns | P value summary | ns | ns | ns | P value summary | ns | ns | ns |
| Brown-Forsythe test |  | Bartlett's test |  | Brown-Forsythe test |  | Bartlett's test |  | Brown-Forsythe test |  | Bartlett's test |  |
| 0.2509 (ns) |  | 0.0955 (ns) |  | 0.8927 (ns) |  | 0.8835 (ns) |  | 0.2684 (ns) |  | 0.0897 (ns) |  |
| 0.6875 (ns) |  | 0.1864 (ns) |  | 0.6875 (ns) |  | 0.1864 (ns) |  | 0.6875 (ns) |  | 0.1864 (ns) |  |
| Ordinary One-way Anova | CS0 | CS50 |  | Ordinary One-way Anova | CS0 | CS50 |  | Ordinary One-way Anova | CS0 | CS50 |  |
| CS50 | 0.1182 (ns) | X |  | CS50 | 0.8242 (ns) | X |  | CS50 | 0.3714 (ns) | X |  |
| CS100 | 0.0191 (*) | 0.4945 (ns) |  | CS100 | 0.2444 (ns) | 0.5161 (ns) |  | CS100 | 0.6486 (ns) | 0.1176 (ns) |  |

**Figure S26:** Calculated partition coefficients of GK6 and HSA in CS100 hydrogels as a function of HSA concentration, assuming no HSA-ECM binding (A) and no peptide binding to HSA (B). Calculated peptide concentrations in the different states for the same system as in (B).

### Diffusion of different peptides measured by FRAP

**Figure S27:** rFRAP model fitting on recovery curves of GK6 measured in each medium

**Table S13:** Statistical comparison of the diffusion coefficients of GK6 measured in different hydrogel types at different HSA concentrations (Related to Figure 5A)

| Effect of the HSA concentration in each gel type |  |  |  |  |  |  |  |  |  |  |  |  |  |  |
| --- | --- | --- | --- | --- | --- | --- | --- | --- | --- | --- | --- | --- | --- | --- |
| PBS |  |  |  |  | Gel type CS0 |  |  |  |  | Gel type CS25 |  |  |  |  |
| Shapiro-Wilk test | PBS | 1mg/ml | 7.5mg/ml | 25mg/ml | Shapiro-Wilk test | PBS | 1mg/ml | 7.5mg/ml | 25mg/ml | Shapiro-Wilk test | PBS | 1mg/ml | 7.5mg/ml | 25mg/ml |
| P value | 0.6300 | 0.1805 | 0.4842 | 0.1941 | P value | 0.6177 | 0.3128 | 0.5549 | 0.9013 | P value | 0.9053 | 0.3379 | 0.0797 | 0.9338 |
| Passed normality test (alpha=0.05)? | Yes | Yes | Yes | Yes | Passed normality test (alpha=0.05)? | Yes | Yes | Yes | Yes | Passed normality test (alpha=0.05)? | Yes | Yes | Yes | Yes |
| P value summary | ns | ns | ns | ns | P value summary | ns | ns | ns | ns | P value summary | ns | ns | ns | ns |
| Brown-Forsythe test |  | Bartlett's test |  |  | Brown-Forsythe test |  | Bartlett's test |  |  | Brown-Forsythe test |  | Bartlett's test |  |  |
| 0.4874 (ns) |  | 0.2472 (ns) |  |  | <0.0001 (****) |  | 0.0002 (***) |  |  | 0.0680 (ns) |  | 0.0016 (**) |  |  |
| Ordinary One-way Anova | PBS | 1mg/ml | 7.5mg/ml |  | Anova test with Welch correction | PBS | 1mg/ml | 7.5mg/ml |  | Anova test with Welch correction | PBS | 1mg/ml | 7.5mg/ml |  |
| 1mg/ml | 0.9949 (ns) | X | X |  | 1mg/ml | 0.9577 (ns) | X | X |  | 1mg/ml | 0.5219 (ns) | X | X |  |
| 7.5mg/ml | <0.0001 (****) | <0.0001 (****) | X |  | 7.5mg/ml | 0.0437 (*) | 0.0018 (**) | X |  | 7.5mg/ml | 0.0035 (**) | 0.0075 (**) | X |  |
| 25mg/ml | <0.0001 (****) | <0.0001 (****) | <0.0001 (****) |  | 25mg/ml | 0.0003 (***) | <0.0001 (****) | <0.0001 (****) |  | 25mg/ml | <0.0001 (****) | <0.0001 (****) | <0.0001 (****) |  |

| Gel type CS50 |  |  |  |  | Gel type CS75 |  |  |  |  | Gel type CS100 |  |  |  |  |
| --- | --- | --- | --- | --- | --- | --- | --- | --- | --- | --- | --- | --- | --- | --- |
| Shapiro-Wilk test | PBS | 1mg/ml | 7.5mg/ml | 25mg/ml | Shapiro-Wilk test | PBS | 1mg/ml | 7.5mg/ml | 25mg/ml | Shapiro-Wilk test | PBS | 1mg/ml | 7.5mg/ml | 25mg/ml |
| P value | 0.6677 | 0.8365 | 0.0525 | 0.1972 | P value | 0.6562 | 0.2078 | 0.0555 | 0.5234 | P value | 0.8132 | 0.0541 | 0.6504 | 0.9042 |
| Passed normality test (alpha=0.05)? | Yes | Yes | Yes | Yes | Passed normality test (alpha=0.05)? | Yes | Yes | Yes | Yes | Passed normality test (alpha=0.05)? | Yes | Yes | Yes | Yes |
| P value summary | ns | ns | ns | ns | P value summary | ns | ns | ns | ns | P value summary | ns | ns | ns | ns |
| Brown-Forsythe test |  | Bartlett's test |  |  | Brown-Forsythe test |  | Bartlett's test |  |  | Brown-Forsythe test |  | Bartlett's test |  |  |
| 0.4212 (ns) |  | 0.0903 (ns) |  |  | 0.4569 (ns) |  | 0.2757 (ns) |  |  | 0.1110 (ns) |  | <0.0001 (****) |  |  |
| Ordinary One-way Anova | PBS | 1mg/ml | 7.5mg/ml |  | Ordinary One-way Anova | PBS | 1mg/ml | 7.5mg/ml |  | Anova test with Welch correction | PBS | 1mg/ml | 7.5mg/ml |  |
| 1mg/ml | 0.9631 (ns) | X | X |  | 1mg/ml | 0.8291 (ns) | X | X |  | 1mg/ml | 0.9962 (ns) | X | X |  |
| 7.5mg/ml | 0.0095 (**) | 0.0028 (**) | X |  | 7.5mg/ml | 0.0768 (ns) | 0.0108 (*) | X |  | 7.5mg/ml | 0.5967 (*) | 0.8622 (ns) | X |  |
| 25mg/ml | <0.0001 (****) | <0.0001 (****) | <0.0001 (****) |  | 25mg/ml | <0.0001 (****) | <0.0001 (****) | <0.0001 (****) |  | 25mg/ml | <0.0001 (****) | 0.0258 (*) | <0.0001 (****) |  |

| Effect of the gel types at each HSA concentration |  |  |  |  |  |  |  |  |  |
| --- | --- | --- | --- | --- | --- | --- | --- | --- | --- |
| PBS |  |  |  |  | 1mg/ml |  |  |  |  |
| Brown-Forsythe test |  | Bartlett's test |  |  | Brown-Forsythe test |  | Bartlett's test |  |  |
| 0.0004 (***) |  | 0.0054 (**) |  |  | 0.3455 (ns) |  | 0.0241 (*) |  |  |
| Anova test with Welch correction | CS0 | CS25 | CS50 | CS75 | Anova test with Welch correction | CS0 | CS25 | CS50 | CS75 |
| CS25 | 0.6505 (ns) | X | X | X | CS25 | 0.0133 (*) | X | X | X |
| CS50 | 0.0090 (**) | 0.0059 (**) | X | X | CS50 | 0.0003 (****) | 0.0482 (*) | X | X |
| CS75 | 0.0019 (**) | 0.0003 (****) | 0.0554 (ns) | X | CS75 | <0.0001 (****) | 0.0005 (****) | 0.0943 (ns) | X |
| CS100 | 0.0009 (***) | <0.0001 (****) | 0.0233 (*) | 0.8299 (ns) | CS100 | 0.0005 (****) | 0.0348 (*) | 0.4098 (ns) | 0.9815 (ns) |

| 7.5mg/ml |  |  |  |  | 25mg/ml |  |  |  |  |
| --- | --- | --- | --- | --- | --- | --- | --- | --- | --- |
| Brown-Forsythe test |  | Bartlett's test |  |  | Brown-Forsythe test |  | Bartlett's test |  |  |
| 0.8671 (ns) |  | 0.3666 (ns) |  |  | 0.9142 (ns) |  | 0.8865 (ns) |  |  |
| Ordinary one-way Anova | CS0 | CS25 | CS50 | CS75 | Ordinary one-way Anova | CS0 | CS25 | CS50 | CS75 |
| CS25 | 0.0194 (*) | X | X | X | CS25 | 0.0001 (****) | X | X | X |
| CS50 | <0.0001 (****) | 0.0030 (**) | X | X | CS50 | <0.0001 (****) | 0.0159 (*) | X | X |
| CS75 | <0.0001 (****) | <0.0001 (****) | 0.6027 (ns) | X | CS75 | <0.0001 (****) | <0.0001 (****) | 0.3404 (ns) | X |
| CS100 | <0.0001 (****) | <0.0001 (****) | 0.4513 (ns) | 0.9991 (ns) | CS100 | <0.0001 (****) | <0.0001 (****) | 0.0193 (*) | 0.6679 (ns) |

**Table S14:** Statistical comparison of the diffusion coefficients of GK6 measured in different hydrogel types at different dextran concentrations (Related to Figure 5A)

| Effect of the dextran concentration in each gel type |  |  |  |  |  |  |  |  |  |  |  |  |  |  |
| --- | --- | --- | --- | --- | --- | --- | --- | --- | --- | --- | --- | --- | --- | --- |
| PBS |  |  |  |  | Gel type CS0 |  |  |  |  | Gel type CS25 |  |  |  |  |
| Shapiro-Wilk test | PBS | 1mg/ml | 7.5mg/ml | 25mg/ml | Shapiro-Wilk test | PBS | 1mg/ml | 7.5mg/ml | 25mg/ml | Shapiro-Wilk test | PBS | 1mg/ml | 7.5mg/ml | 25mg/ml |
| P value | 0.1545 | 0.1490 | 0.1626 | 0.6452 | P value | 0.6177 | 0.3358 | 0.8134 | 0.8014 | P value | 0.9053 | 0.5769 | 0.2898 | 0.1773 |
| Passed normality test (alpha=0.05)? | Yes | Yes | Yes | Yes | Passed normality test (alpha=0.05)? | Yes | Yes | Yes | Yes | Passed normality test (alpha=0.05)? | Yes | Yes | Yes | Yes |
| P value summary | ns | ns | ns | ns | P value summary | ns | ns | ns | ns | P value summary | ns | ns | ns | ns |
| Brown-Forsythe test |  | Bartlett's test |  |  | Brown-Forsythe test |  | Bartlett's test |  |  | Brown-Forsythe test |  | Bartlett's test |  |  |
| 0.1571 (ns) |  | 0.0723 (ns) |  |  | 0.1825 (ns) |  | 0.0587 (ns) |  |  | 0.4344 (ns) |  | 0.2395 (ns) |  |  |
| Ordinary One-way Anova | PBS | 1mg/ml | 7.5mg/ml |  | Ordinary One-way Anova | PBS | 1mg/ml | 7.5mg/ml |  | Ordinary One-way Anova | PBS | 1mg/ml | 7.5mg/ml |  |
| 1mg/ml | 0.1765 (ns) | X | X |  | 1mg/ml | 0.5448 (ns) | X | X |  | 1mg/ml | 0.0114 (*) | X | X |  |
| 7.5mg/ml | 0.00631 (**) | 0.0003 (***) | X |  | 7.5mg/ml | 0.0837 (ns) | 0.7038 (ns) | X |  | 7.5mg/ml | 0.0850 (ns) | 0.8179 (ns) | X |  |
| 25mg/ml | <0.0001 (****) | <0.0001 (****) | <0.0001 (****) |  | 25mg/ml | 0.0101 (*) | 0.2221 (ns) | 0.7950 (ns) |  | 25mg/ml | 0.0033 (**) | 0.9565 (ns) | 0.5257 (ns) |  |
| Gel type CS50 |  |  |  |  | Gel type CS75 |  |  |  |  | Gel type CS100 |  |  |  |  |
| Shapiro-Wilk test | PBS | 1mg/ml | 7.5mg/ml | 25mg/ml | Shapiro-Wilk test | PBS | 1mg/ml | 7.5mg/ml | 25mg/ml | Shapiro-Wilk test | PBS | 1mg/ml | 7.5mg/ml | 25mg/ml |
| P value | 0.6677 | 0.9803 | 0.8888 | 0.0093 | P value | 0.6562 | 0.5227 | 0.4060 | 0.8043 | P value | 0.8132 | 0.7422 | 0.2565 | 0.5666 |
| Passed normality test (alpha=0.05)? | Yes | Yes | Yes | no | Passed normality test (alpha=0.05)? | Yes | Yes | Yes | Yes | Passed normality test (alpha=0.05)? | Yes | Yes | Yes | Yes |
| P value summary | ns | ns | ns | ** | P value summary | ns | ns | ns | ns | P value summary | ns | ns | ns | ns |
| Kruskal-Wallis |  | Approximate P value |  |  | Brown-Forsythe test |  | Bartlett's test |  |  | Brown-Forsythe test |  | Bartlett's test |  |  |
| 0.0132 (*) |  |  |  |  | 0.4011 (ns) |  | 0.2197 (ns) |  |  | 0.2231 (ns) |  | 0.5207 (ns) |  |  |
| Kruskal-Wallis test | PBS | 1mg/ml | 7.5mg/ml |  | Ordinary One-way Anova | PBS | 1mg/ml | 7.5mg/ml |  | Ordinary One-way Anova | PBS | 1mg/ml | 7.5mg/ml |  |
| 1mg/ml | >0.9999 (ns) | X | X |  | 1mg/ml | 0.2512 (ns) | X | X |  | 1mg/ml | 0.9986 (ns) | X | X |  |
| 7.5mg/ml | >0.9999 (ns) | >0.9999 (ns) | X |  | 7.5mg/ml | 0.2912 (ns) | 0.9998 (ns) | X |  | 7.5mg/ml | 0.8899 (ns) | 0.8599 (ns) | X |  |
| 25mg/ml | 0.0066 (**) | 0.2413 (ns) | 0.4140 (ns) |  | 25mg/ml | 0.4684 (ns) | 0.9775 (ns) | 0.9889 (ns) |  | 25mg/ml | 0.9505 (ns) | 0.9245 (ns) | 0.9983 (ns) |  |
| Effect of the gel types at each dextran concentration |  |  |  |  |  |  |  |  |  |  |  |  |  |  |
| 1mg/ml |  |  |  |  | 7.5mg/ml |  |  |  |  | 25mg/ml |  |  |  |  |
| Brown-Forsythe test |  | Bartlett's test |  |  | Brown-Forsythe test |  | Bartlett's test |  |  | Kruskal-Wallis |  | Approximate P value |  |  |
| 0.1877 (ns) |  | 0.0507 (ns) |  |  | 0.0651 (ns) |  | 0.0508 (ns) |  |  | 0.0007 (***) |  |  |  |  |
| Ordinary one-way Anova | CS0 | CS25 | CS50 | CS75 | Ordinary one-way Anova | CS0 | CS25 | CS50 | CS75 | Kruskal-Wallis test | CS0 | CS25 | CS50 | CS75 |
| CS25 | 0.2133 (ns) | X | X | X | CS25 | >0.9999 (ns) | X | X | X | CS25 | >0.9999 (ns) | X | X | X |
| CS50 | 0.0282 (*) | 0.8305 (ns) | X | X | CS50 | 0.0314 (*) | 0.0353 (*) | X | X | CS50 | 0.0043 (**) | 0.0094 (**) | X | X |
| CS75 | 0.0027 (**) | 0.2562 (ns) | 0.8251 (ns) | X | CS75 | 0.0008 (***) | 0.0009 (***) | 0.4943 (ns) | X | CS75 | 0.2843 (ns) | 0.4810 (ns) | >0.9999 (ns) | X |
| CS100 | 0.0040 (**) | 0.3307 (ns) | 0.8971 (ns) | 0.9998 (ns) | CS100 | 0.0003 (***) | 0.0004 (***) | 0.3895 (ns) | 0.9945 (ns) | CS100 | 0.0472 (*) | 0.1432 (ns) | >0.9999 (ns) | >0.9999 (ns) |

**Table S15:** Statistical comparison of the diffusion coefficients of GK6 measured in the void and the collagen-rich domains of different hydrogel types, and Ds measured in PBS compared to those measured in the voids of the different hydrogels (Related to Figure 5A)

| Collagen domain vs void in different gels |  |  |  |  |  | PBS vs void |  |  |  |
| --- | --- | --- | --- | --- | --- | --- | --- | --- | --- |
|  |  |  |  |  |  | No HSA |  | No HSA |  |
| Unpaired t test | CS0 | CS25 | CS50 | CS75 | CS100 | Ordinary one-way Anova | PBS | Ordinary one-way Anova | PBS |
| No HSA | 0.4940 (ns) | 0.8656 (ns) | <0.0001 (****) | <0.0001 (****) | <0.0001 (****) |  |  |  |  |
| 1mg/ml HSA | 0.3575 (ns) | 0.0219 (*) | 0.0006 (***) | 0.0001 (***) | 0.0036 (**) | CS25 | 0.0371 (*) | CS75 | 0.0231 (*) |
| 7.5mg/ml HSA | 0.1187 (ns) | 0.0033 (**) | 0.0020 (**) | <0.0001 (****) | <0.0001 (****) | CS75 | 0.0140 (*) | CS100 | 0.0357 (*) |
| 25mg/ml HSA | 0.0003 (***) | 0.0443 (*) | 0.0006 (***) | <0.0001 (****) | <0.0001 (****) | CS100 | 0.0383 (*) |  |  |

**Table S16:** Statistical comparison of the diffusion coefficients of RM26 measured in different hydrogel types at different HSA and dextran concentrations (Related to Figure 5C and Figure S30)

| Effect of the HSA concentration in each gel type |  |  |  |  |  |  |  |  |  |  |  |  |  |  |
| --- | --- | --- | --- | --- | --- | --- | --- | --- | --- | --- | --- | --- | --- | --- |
| N<sub>g</sub>=2 N<sub>m</sub>=3/3 |  |  |  |  | Gel type CS0 |  |  |  |  | Gel type CS25 |  |  |  |  |
| Shapiro-Wilk test | PBS | 1mg/ml | 7.5mg/ml | 25mg/ml | Shapiro-Wilk test | PBS | 1mg/ml | 7.5mg/ml | 25mg/ml | Shapiro-Wilk test | PBS | 1mg/ml | 7.5mg/ml | 25mg/ml |
| P value | 0.1619 | 0.0556 | 0.2423 | 0.5413 | P value | 0.4115 | 0.5692 | 0.0893 | 0.3785 | P value | 0.6211 | 0.4017 | 0.7194 | 0.7295 |
| Passed normality test (alpha=0.05)? | Yes | Yes | Yes | Yes | Passed normality test (alpha=0.05)? | Yes | Yes | Yes | Yes | Passed normality test (alpha=0.05)? | Yes | Yes | Yes | Yes |
| P value summary | ns | ns | ns | ns | P value summary | ns | ns | ns | ns | P value summary | ns | ns | ns | ns |
| Brown-Forsythe test |  | Bartlett's test |  |  | Brown-Forsythe test |  | Bartlett's test |  |  | Brown-Forsythe test |  | Bartlett's test |  |  |
| 0.4897 (ns) |  | 0.5226 (ns) |  |  | 0.5882 (ns) |  | 0.0770 (ns) |  |  | 0.4057 (ns) |  | 0.1769 (ns) |  |  |
| Ordinary One-way Anova | PBS | 1mg/ml | 7.5mg/ml |  | Ordinary One-way Anova | PBS | 1mg/ml | 7.5mg/ml |  | Ordinary One-way Anova | PBS | 1mg/ml | 7.5mg/ml |  |
| 1mg/ml | 0.4758 (ns) | X | X |  | 1mg/ml | >0.9999 (ns) | X | X |  | 1mg/ml | 0.4839 (ns) | X | X |  |
| 7.5mg/ml | <0.0001 (****) | 0.0013 (**) | X |  | 7.5mg/ml | <0.0001 (****) | <0.0001 (****) | X |  | 7.5mg/ml | <0.0001 (****) | <0.0001 (****) | X |  |
| 25mg/ml | <0.0001 (****) | <0.0001 (****) | <0.0001 (****) |  | 25mg/ml | <0.0001 (****) | <0.0001 (****) | <0.0001 (****) |  | 25mg/ml | <0.0001 (****) | <0.0001 (****) | <0.0001 (****) |  |

| Gel type CS50 |  |  |  |  | Gel type CS75 |  |  |  |  | Gel type CS100 |  |  |  |  |
| --- | --- | --- | --- | --- | --- | --- | --- | --- | --- | --- | --- | --- | --- | --- |
| Shapiro-Wilk test | PBS | 1mg/ml | 7.5mg/ml | 25mg/ml | Shapiro-Wilk test | PBS | 1mg/ml | 7.5mg/ml | 25mg/ml | Shapiro-Wilk test | PBS | 1mg/ml | 7.5mg/ml | 25mg/ml |
| P value | 0.9103 | 0.6629 | 0.4848 | 0.1123 | P value | 0.1060 | 0.2576 | 0.6819 | 0.9735 | P value | 0.7931 | 0.5215 | 0.9960 | 0.3192 |
| Passed normality test (alpha=0.05)? | Yes | Yes | Yes | Yes | Passed normality test (alpha=0.05)? | Yes | Yes | Yes | Yes | Passed normality test (alpha=0.05)? | Yes | Yes | Yes | Yes |
| P value summary | ns | ns | ns | ns | P value summary | ns | ns | ns | ns | P value summary | ns | ns | ns | ns |
| Brown-Forsythe test |  | Bartlett's test |  |  | Brown-Forsythe test |  | Bartlett's test |  |  | Brown-Forsythe test |  | Bartlett's test |  |  |
| 0.6809 (ns) |  | 0.4896 (ns) |  |  | 0.1351 (ns) |  | 0.0052 (*) |  |  | 0.1513 (ns) |  | 0.1582 (ns) |  |  |
| Ordinary One-way Anova | PBS | 1mg/ml | 7.5mg/ml |  | Anova test with Welch correction | PBS | 1mg/ml | 7.5mg/ml |  | Ordinary One-way Anova | PBS | 1mg/ml | 7.5mg/ml |  |
| 1mg/ml | 0.1222 (ns) | X | X |  | 1mg/ml | 0.7989 (ns) | X | X |  | 1mg/ml | 0.9403 (ns) | X | X |  |
| 7.5mg/ml | <0.0001 (****) | <0.0001 (****) | X |  | 7.5mg/ml | 0.0005 (***) | 0.0007 (***) | X |  | 7.5mg/ml | <0.0001 (****) | <0.0001 (****) | X |  |
| 25mg/ml | <0.0001 (****) | <0.0001 (****) | <0.0001 (****) |  | 25mg/ml | <0.0001 (****) | 0.0001 (***) | <0.0001 (****) |  | 25mg/ml | <0.0001 (****) | <0.0001 (****) | 0.0020 (**) |  |

| Effect of the gel types at each HSA concentration |  |  |  |  | Effect of dextran concentration |  |  |  |  |
| --- | --- | --- | --- | --- | --- | --- | --- | --- | --- |
| N<sub>g</sub>=2 N<sub>m</sub>=3/3 |  |  |  |  | N<sub>g</sub>=2 N<sub>m</sub>=2/3 |  |  |  |  |
| 7.5mg/ml |  |  |  |  | PBS |  |  |  |  |
| Brown-Forsythe test |  | Bartlett's test |  |  | Shapiro-Wilk test | PBS | 1mg/ml | 7.5mg/ml | 25mg/ml |
| 0.6690 (ns) |  | 0.1478 (ns) |  |  | P value | 0.1619 | 0.0953 | 0.5593 | 0.0883 |
| Ordinary One-way Anova | CS0 | CS25 | CS50 | CS75 | Passed normality test (alpha=0.05)? | Yes | Yes | Yes | Yes |
| CS25 | 0.5330 (ns) | X | X | X | P value summary | ns | ns | ns | ns |
| CS50 | 0.1805 (ns) | 0.9421 (ns) | X | X | Brown-Forsythe test |  | Bartlett's test |  |  |
| CS75 | <0.0001 (****) | 0.0025 (**) | 0.0136 (*) | X | 0.5938 (ns) |  | 0.7034 (ns) |  |  |
| CS100 | <0.0001 (****) | <0.0001 (****) | <0.0001 (****) | 0.1225 (ns) | Ordinary One-way Anova | PBS | 1mg/ml | 7.5mg/ml |  |

| Hydrogel vs PBS in different gels |  |  |  |  |  |
| --- | --- | --- | --- | --- | --- |
| Unpaired t test | CS0 | CS25 | CS50 | CS75 | CS100 |
| 7.5mg/ml HSA | <0.0001 (****) | <0.0001 (****) | <0.0001 (****) | <0.0001 (****) | <0.0001 (****) |
| 25mg/ml HSA | <0.0001 (****) | <0.0001 (****) | <0.0001 (****) | <0.0001 (****) | <0.0001 (****) |

| Dextran concentration has no significant effect in gels |  |  |  |  |
| --- | --- | --- | --- | --- |
| Shapiro-Wilk test | PBS | 1mg/ml | 7.5mg/ml | 25mg/ml |
| P value | 0.1619 | 0.0953 | 0.5593 | 0.0883 |
| Passed normality test (alpha=0.05)? | Yes | Yes | Yes | Yes |
| P value summary | ns | ns | ns | ns |
| Brown-Forsythe test |  | Bartlett's test |  |  |
| 0.5938 (ns) |  | 0.7034 (ns) |  |  |
| Ordinary One-way Anova | PBS | 1mg/ml | 7.5mg/ml |  |
| 1mg/ml | 0.9457 (ns) | X | X |  |
| 7.5mg/ml | 0.4876 (ns) | 0.2291 (ns) | X |  |
| 25mg/ml | <0.0001 (****) | <0.0001 (****) | 0.0005 (***) |  |

**Table S17:** Statistical comparison of the diffusion coefficients of GK3 measured in different hydrogel types at different HSA and dextran concentrations (Related to Figure 5C and Figure S30)

| Effect of the HSA concentration in each gel type |  |  |  |  |  |  |  |  |  |  |  |  |  |  |
| --- | --- | --- | --- | --- | --- | --- | --- | --- | --- | --- | --- | --- | --- | --- |
| N<sub>g</sub>=2 N<sub>m</sub>=3 |  |  |  |  | Gel type CS0 |  |  |  |  | Gel type CS25 |  |  |  |  |
| Shapiro-Wilk test | PBS | 1mg/ml | 7.5mg/ml | 25mg/ml | Shapiro-Wilk test | PBS | 1mg/ml | 7.5mg/ml | 25mg/ml | Shapiro-Wilk test | PBS | 1mg/ml | 7.5mg/ml | 25mg/ml |
| P value | 0.0866 | 0.9981 | 0.0564 | 0.8316 | P value | 0.9089 | 0.1337 | 0.2762 | 0.5389 | P value | 0.0536 | 0.2158 | 0.0522 | 0.7063 |
| Passed normality test (alpha=0.05)? | Yes | Yes | Yes | Yes | Passed normality test (alpha=0.05)? | Yes | Yes | Yes | Yes | Passed normality test (alpha=0.05)? | Yes | Yes | Yes | Yes |
| P value summary | ns | ns | ns | ns | P value summary | ns | ns | ns | ns | P value summary | ns | ns | ns | ns |
| Brown-Forsythe test |  | Bartlett's test |  |  | Brown-Forsythe test |  | Bartlett's test |  |  | Brown-Forsythe test |  | Bartlett's test |  |  |
| 0.3267 (ns) |  | 0.1038 (ns) |  |  | 0.0391 (*) |  | 0.0683 (ns) |  |  | 0.3079 (ns) |  | 0.0869 (ns) |  |  |
| Ordinary One-way Anova | PBS | 1mg/ml | 7.5mg/ml |  | Anova test with Welch correction | PBS | 1mg/ml | 7.5mg/ml |  | Ordinary One-way Anova | PBS | 1mg/ml | 7.5mg/ml |  |
| 1mg/ml | 0.0031 (**) | X | X |  | 1mg/ml | 0.6646 (ns) | X | X |  | 1mg/ml | 0.8667 (ns) | X | X |  |
| 7.5mg/ml | <0.0001 (****) | <0.0001 (****) | X |  | 7.5mg/ml | 0.0034 (**) | 0.0043 (**) | X |  | 7.5mg/ml | <0.0001 (****) | 0.0003 (****) | X |  |
| 25mg/ml | <0.0001 (****) | <0.0001 (****) | <0.0001 (****) |  | 25mg/ml | <0.0001 (****) | <0.0001 (****) | <0.0001 (****) |  | 25mg/ml | <0.0001 (****) | <0.0001 (****) | 0.0003 (****) |  |

| Gel type CS50 |  |  |  |  | Gel type CS75 |  |  |  |  | Gel type CS100 |  |  |  |  |
| --- | --- | --- | --- | --- | --- | --- | --- | --- | --- | --- | --- | --- | --- | --- |
| Shapiro-Wilk test | PBS | 1mg/ml | 7.5mg/ml | 25mg/ml | Shapiro-Wilk test | PBS | 1mg/ml | 7.5mg/ml | 25mg/ml | Shapiro-Wilk test | PBS | 1mg/ml | 7.5mg/ml | 25mg/ml |
| P value | 0.1852 | 0.2291 | 0.8556 | 0.2589 | P value | 0.2768 | 0.6211 | 0.3280 | 0.3599 | P value | 0.7898 | 0.3421 | 0.9013 | 0.4418 |
| Passed normality test (alpha=0.05)? | Yes | Yes | Yes | Yes | Passed normality test (alpha=0.05)? | Yes | Yes | Yes | Yes | Passed normality test (alpha=0.05)? | Yes | Yes | Yes | Yes |
| P value summary | ns | ns | ns | ns | P value summary | ns | ns | ns | ns | P value summary | ns | ns | ns | ns |
| Brown-Forsythe test |  | Bartlett's test |  |  | Brown-Forsythe test |  | Bartlett's test |  |  | Brown-Forsythe test |  | Bartlett's test |  |  |
| 0.1140 (ns) |  | 0.0921 (ns) |  |  | 0.0642 (ns) |  | 0.0291 (*) |  |  | 0.0042 (**) |  | 0.0128 (*) |  |  |
| Ordinary One-way Anova | PBS | 1mg/ml | 7.5mg/ml |  | Anova test with Welch correction | PBS | 1mg/ml | 7.5mg/ml |  | Anova test with Welch correction | PBS | 1mg/ml | 7.5mg/ml |  |
| 1mg/ml | 0.9330 (ns) | X | X |  | 1mg/ml | 0.5677 (ns) | X | X |  | 1mg/ml | 0.9965 (ns) | X | X |  |
| 7.5mg/ml | <0.0001 (****) | <0.0001 (****) | X |  | 7.5mg/ml | 0.3285 (ns) | 0.1016 (ns) | X |  | 7.5mg/ml | 0.2043 (ns) | 0.0028 (**) | X |  |
| 25mg/ml | <0.0001 (****) | <0.0001 (****) | <0.0001 (****) |  | 25mg/ml | <0.0001 (****) | <0.0001 (****) | 0.0024 (**) |  | 25mg/ml | 0.0013 (**) | <0.0001 (****) | <0.0001 (****) |  |

| Effect of the gel types at each HSA concentration |  |  |  |  |  |  |  |  |  |  |  |  |  |  |
| --- | --- | --- | --- | --- | --- | --- | --- | --- | --- | --- | --- | --- | --- | --- |
| PBS |  |  |  |  | 1mg/ml |  |  |  |  | 7.5mg/ml |  |  |  |  |
| Brown-Forsythe test |  | Bartlett's test |  |  | Brown-Forsythe test |  | Bartlett's test |  |  | Brown-Forsythe test |  | Bartlett's test |  |  |
| 0.0184 (*) |  | 0.0008 (***) |  |  | 0.6707 (ns) |  | 0.7598 (ns) |  |  | 0.0070 (**) |  | 0.0075 (**) |  |  |
| Anova test with Welch correction | CS0 | CS25 | CS50 | CS75 | Ordinary One-way Anova | CS0 | CS25 | CS50 | CS75 | Anova test with Welch correction | CS0 | CS25 | CS50 | CS75 |
| CS25 | >0.9999 (ns) | X | X | X | CS25 | 0.9967 (ns) | X | X | X | CS25 | >0.9999 (ns) | X | X | X |
| CS50 | 0.9031 (ns) | 0.9622 (ns) | X | X | CS50 | >0.9999 (ns) | 0.9994 (ns) | X | X | CS50 | 0.9906 (ns) | 0.9807 (ns) | X | X |
| CS75 | 0.3534 (ns) | 0.5454 (ns) | 0.4144 (ns) | X | CS75 | 0.9621 (ns) | 0.9976 (ns) | 0.9820 (ns) | X | CS75 | 0.0060 (**) | 0.0047 (**) | 0.0178 (*) | X |
| CS100 | 0.1373 (ns) | 0.1923 (ns) | 0.2033 (ns) | 0.6393 (ns) | CS100 | 0.1380 (ns) | 0.0490 (*) | 0.1088 (ns) | 0.0346 (*) | CS100 | 0.0002 (****) | 0.0002 (****) | 0.0006 (****) | 0.6665 (ns) |

| Effect of the dextran concentration |  |  |  |  |  |  |  |  |  |  |  |  |  |  |
| --- | --- | --- | --- | --- | --- | --- | --- | --- | --- | --- | --- | --- | --- | --- |
| N<sub>g</sub>=2 N<sub>m</sub>=2 and 3 |  |  |  |  | PBS |  |  |  |  | 7.5mg/ml |  |  |  |  |
| Shapiro-Wilk test | PBS | 1mg/ml | 7.5mg/ml | 25mg/ml | Shapiro-Wilk test | PBS | 1mg/ml | 7.5mg/ml | 25mg/ml | Shapiro-Wilk test | PBS | 1mg/ml | 7.5mg/ml | 25mg/ml |
| P value | 0.8404 | 0.9458 | 0.9839 | 0.9819 | P value | 0.8404 | 0.9458 | 0.9839 | 0.9819 | P value | 0.8404 | 0.9458 | 0.9839 | 0.9819 |
| Passed normality test (alpha=0.05)? | Yes | Yes | Yes | Yes | Passed normality test (alpha=0.05)? | Yes | Yes | Yes | Yes | Passed normality test (alpha=0.05)? | Yes | Yes | Yes | Yes |
| P value summary | ns | ns | ns | ns | P value summary | ns | ns | ns | ns | P value summary | ns | ns | ns | ns |
| Brown-Forsythe test |  | Bartlett's test |  |  | Brown-Forsythe test |  | Bartlett's test |  |  | Brown-Forsythe test |  | Bartlett's test |  |  |
| 0.4657 (ns) |  | 0.4978 (ns) |  |  | 0.4657 (ns) |  | 0.4978 (ns) |  |  | 0.4657 (ns) |  | 0.4978 (ns) |  |  |
| Ordinary One-way Anova | PBS | 1mg/ml | 7.5mg/ml |  | Ordinary One-way Anova | PBS | 1mg/ml | 7.5mg/ml |  | Ordinary One-way Anova | PBS | 1mg/ml | 7.5mg/ml |  |
| 1mg/ml | 0.0192 (*) | X | X |  | 1mg/ml | 0.0192 (*) | X | X |  | 1mg/ml | 0.0192 (*) | X | X |  |
| 7.5mg/ml | 0.6374 (ns) | 0.0037 (**) | X |  | 7.5mg/ml | 0.6374 (ns) | 0.0037 (**) | X |  | 7.5mg/ml | 0.6374 (ns) | 0.0037 (**) | X |  |
| 25mg/ml | 0.0026 (**) | <0.0001 (****) | 0.0623 (ns) |  | 25mg/ml | 0.0026 (**) | <0.0001 (****) | 0.0623 (ns) |  | 25mg/ml | 0.0026 (**) | <0.0001 (****) | 0.0623 (ns) |  |

Dextran concentration has no significant effect in the gels

**Table S18:** Statistical comparison of the diffusion coefficients of KWEA measured in different hydrogel types at different HSA and dextran concentrations (Related to Figure 5C and Figure S30)

| Effect of the HSA concentration in each gel type |  |  |  |  |  |  |  |  |  |  |  |  |  |  |
| --- | --- | --- | --- | --- | --- | --- | --- | --- | --- | --- | --- | --- | --- | --- |
| N<sub>g</sub>=2 N<sub>m</sub>=2/3 |  |  |  | PBS |  |  |  | Gel type CS0 |  |  |  | Gel type CS25 |  |  |
| Shapiro-Wilk test | PBS | 1mg/ml | 7.5mg/ml | 25mg/ml | Shapiro-Wilk test | PBS | 1mg/ml | 7.5mg/ml | 25mg/ml | Shapiro-Wilk test | PBS | 1mg/ml | 7.5mg/ml | 25mg/ml |
| P value | 0.4393 | 0.8256 | 0.3966 | 0.5627 | P value | 0.7919 | 0.1509 | 0.7778 | 0.0516 | P value | 0.8592 | 0.9983 | 0.6275 | 0.9974 |
| Passed normality test (alpha=0.05)? | Yes | Yes | Yes | Yes | Passed normality test (alpha=0.05)? | Yes | Yes | Yes | Yes | Passed normality test (alpha=0.05)? | Yes | Yes | Yes | Yes |
| P value summary | ns | ns | ns | ns | P value summary | ns | ns | ns | ns | P value summary | ns | ns | ns | ns |
| Brown-Forsythe test |  | Bartlett's test |  | Brown-Forsythe test |  | Bartlett's test |  | Brown-Forsythe test |  | Bartlett's test |  |  |  |  |
| 0.2020 (ns) |  | 0.0491 (*) |  | 0.1921 (ns) |  | 0.0215 (*) |  | 0.1548 (ns) |  | 0.0004 (***) |  |  |  |  |
| Anova test with Welch correction | PBS | 1mg/ml | 7.5mg/ml | Anova test with Welch correction | PBS | 1mg/ml | 7.5mg/ml | Anova test with Welch correction | PBS | 1mg/ml | 7.5mg/ml |  |  |  |
| 1mg/ml | 0.0556 (ns) | X | X | 1mg/ml | 0.0532 (ns) | X | X | 1mg/ml | 0.3397 (ns) | X | X |  |  |  |
| 7.5mg/ml | <0.0001 (****) | <0.0001 (****) | X | 7.5mg/ml | 0.0032 (**) | 0.0219 (*) | X | 7.5mg/ml | 0.0131 (*) | 0.0548 (ns) | X |  |  |  |
| 25mg/ml | <0.0001 (****) | <0.0001 (****) | <0.0001 (****) | 25mg/ml | <0.0001 (****) | <0.0001 (****) | <0.0001 (****) | 25mg/ml | 0.0006 (***) | <0.0001 (****) | 0.0076 (**) |  |  |  |
| Gel type CS50 |  |  |  | Gel type CS75 |  |  |  | Gel type CS100 |  |  |  |  |  |  |
| Shapiro-Wilk test | PBS | 1mg/ml | 7.5mg/ml | 25mg/ml | Shapiro-Wilk test | PBS | 1mg/ml | 7.5mg/ml | 25mg/ml | Shapiro-Wilk test | PBS | 1mg/ml | 7.5mg/ml | 25mg/ml |
| P value | 0.7210 | 0.6915 | 0.0673 | 0.1532 | P value | 0.6652 | 0.6555 | 0.2481 | 0.5088 | P value | 0.2106 | 0.8241 | 0.3513 | 0.5273 |
| Passed normality test (alpha=0.05)? | Yes | Yes | Yes | Yes | Passed normality test (alpha=0.05)? | Yes | Yes | Yes | Yes | Passed normality test (alpha=0.05)? | Yes | Yes | Yes | Yes |
| P value summary | ns | ns | ns | ns | P value summary | ns | ns | ns | ns | P value summary | ns | ns | ns | ns |
| Brown-Forsythe test |  | Bartlett's test |  | Brown-Forsythe test |  | Bartlett's test |  | Brown-Forsythe test |  | Bartlett's test |  |  |  |  |
| 0.4529 (ns) |  | 0.3761 (ns) |  | 0.5176 (ns) |  | 0.2224 (ns) |  | 0.3177 (ns) |  | 0.0006 (***) |  |  |  |  |
| Ordinary One-way Anova | PBS | 1mg/ml | 7.5mg/ml | Ordinary One-way Anova | PBS | 1mg/ml | 7.5mg/ml | Anova test with Welch correction | PBS | 1mg/ml | 7.5mg/ml |  |  |  |
| 1mg/ml | 0.1250 (ns) | X | X | 1mg/ml | 0.5144 (ns) | X | X | 1mg/ml | 0.9780 (ns) | X | X |  |  |  |
| 7.5mg/ml | <0.0001 (****) | 0.0031 (**) | X | 7.5mg/ml | 0.0014 (**) | 0.0237 (*) | X | 7.5mg/ml | 0.2260 (ns) | 0.0180 (*) | X |  |  |  |
| 25mg/ml | <0.0001 (****) | <0.0001 (****) | <0.0001 (****) | 25mg/ml | <0.0001 (****) | 0.0001 (****) | <0.0001 (****) | 25mg/ml | 0.0007 (****) | <0.0001 (****) | 0.0001 (****) |  |  |  |
| Effect of the gel types at each HSA concentration |  |  |  | Effect of dextran concentration |  |  |  |  |  |  |  |  |  |  |
| 25mg/ml |  |  |  | PBS |  |  |  | Gel type CS0 |  |  |  |  |  |  |
| Brown-Forsythe test |  | Bartlett's test |  | Shapiro-Wilk test | PBS | 1mg/ml | 7.5mg/ml | 25mg/ml | Shapiro-Wilk test | PBS | 1mg/ml | 7.5mg/ml | 25mg/ml |  |
| 0.1948 (ns) |  | 0.0008 (**) |  | P value | 0.4393 | 0.1285 | 0.9596 | 0.2624 | P value | 0.7919 | 0.5245 | 0.1335 | 0.1696 |  |
| Passed normality test (alpha=0.05)? |  | Yes |  | Yes | Yes | Yes | Yes | Yes | Passed normality test (alpha=0.05)? | Yes | Yes | Yes | Yes |  |
| P value summary |  | ns |  | ns | ns | ns | ns | ns | P value summary | ns | ns | ns | ns |  |
| Brown-Forsythe test |  | Bartlett's test |  | Brown-Forsythe test |  | Bartlett's test |  | Brown-Forsythe test |  | Bartlett's test |  |  |  |  |
| 0.8610 (ns) |  | 0.8376 (ns) |  | 0.1276 (ns) |  | 0.1374 (ns) |  | 0.1276 (ns) |  | 0.1374 (ns) |  |  |  |  |
| Ordinary One-way Anova | PBS | 1mg/ml | 7.5mg/ml | Ordinary One-way Anova | PBS | 1mg/ml | 7.5mg/ml |  |  |  |  |  |  |  |
| 1mg/ml | 0.9977 (ns) | X | X | 1mg/ml | 0.2426 (ns) | X | X |  |  |  |  |  |  |  |
| 7.5mg/ml | 0.1329 (ns) | 0.1483 (ns) | X | 7.5mg/ml | <0.0001 (****) | 0.0003 (****) | X |  |  |  |  |  |  |  |
| 25mg/ml | <0.0001 (****) | 0.0002 (****) | 0.0215 (*) | 25mg/ml | 0.0369 (*) | 0.7203 (ns) | 0.0028 (**) |  |  |  |  |  |  |  |
| PBS, 1mg/ml, and 7.5mg/ml HSA are not significant among the gels |  |  |  | Gel type CS25 |  |  |  | Gel type CS50 |  |  |  |  |  |  |
| Shapiro-Wilk test |  | PBS | 1mg/ml | 7.5mg/ml | 25mg/ml | Shapiro-Wilk test |  | PBS | 1mg/ml | 7.5mg/ml | 25mg/ml |  |  |  |
| P value |  | 0.8592 | 0.3045 | 0.5286 | 0.7059 | P value |  | 0.7210 | 0.7998 | 0.0545 | 0.9653 |  |  |  |
| Passed normality test (alpha=0.05)? |  | Yes | Yes | Yes | Yes | Passed normality test (alpha=0.05)? |  | Yes | Yes | Yes | Yes |  |  |  |
| P value summary |  | ns | ns | ns | ns | P value summary |  | ns | ns | ns | ns |  |  |  |
| Brown-Forsythe test |  | Bartlett's test |  | Brown-Forsythe test |  | Bartlett's test |  |  |  |  |  |  |  |  |
| 0.1708 (ns) |  | 0.0224 (*) |  | 0.6429 (ns) |  | 0.5235 (ns) |  |  |  |  |  |  |  |  |
| Anova test with Welch correction | PBS | 1mg/ml | 7.5mg/ml | Ordinary One-way Anova | PBS | 1mg/ml | 7.5mg/ml |  |  |  |  |  |  |  |
| 1mg/ml | 0.5777 (ns) | X | X | 1mg/ml | 0.9039 (ns) | X | X |  |  |  |  |  |  |  |
| 7.5mg/ml | 0.5112 (ns) | 0.0061 (**) | X | 7.5mg/ml | 0.8668 (ns) | 0.4896 (ns) | X |  |  |  |  |  |  |  |
| 25mg/ml | 0.2376 (ns) | 0.2874 (ns) | 0.0013 (**) | 25mg/ml | 0.0326 (*) | 0.0084 (**) | 0.1369 (ns) |  |  |  |  |  |  |  |
| Dextran concentration has no significant effect in CS75 and CS100 hydrogels |  |  |  |  |  |  |  |  |  |  |  |  |  |  |

PBS, 1mg/ml, and 7.5mg/ml HSA are not significant among the gels

Dextran concentration has no significant effect in CS75 and CS100 hydrogels

**Figure S28:** Effect of the dextran concentration on the diffusion of GK3, RM26, KWEA, and Exe in different media

**Table S19:** Statistical comparison of the $D$ of Exe measured in PBS with those measured in 25 mg/ml dextran solution

$N_g=1$ $N_m=4$

| Unpaired t test | PBS | CS0 | CS25 | CS50 | CS75 | CS100 |
| --- | --- | --- | --- | --- | --- | --- |
| P-value | <0.0001 (***) | <0.0001 (***) | 0.0196 (*) | <0.0001 (***) | <0.0001 (***) | 0.0026 (**) |

**Figure S29:** rFRAP model fitting on the recovery curves of GK3 measured in each medium

**Figure S30:** rFRAP model fitting on the recovery curves of RM26 measured in each medium

**Figure S31:** rFRAP model fitting on the recovery curves of KWEA measured in each medium

**Figure S32:** rFRAP model fitting on the recovery curves of Exe measured in each medium

| N<sub>g</sub>=2 N<sub>m</sub>=3/3 |  |  |  |  |  | N<sub>g</sub>=2 N<sub>m</sub>=3 |  |  |  |  |  | RM26 |  |  |  |  |  |
| --- | --- | --- | --- | --- | --- | --- | --- | --- | --- | --- | --- | --- | --- | --- | --- | --- | --- |
| GK3 |  |  |  |  |  | N<sub>g</sub>=2 N<sub>m</sub>=3 |  |  |  |  |  | RM26 |  |  |  |  |  |
| Unpaired t test | CS0 | CS25 | CS50 | CS75 | CS100 | Unpaired t test | CS0 | CS25 | CS50 | CS75 | CS100 | Unpaired t test | CS0 | CS25 | CS50 | CS75 | CS100 |
| No HSA | 0.3995 (ns) | 0.5304 (ns) | 0.0011 (**) | 0.0009 (***) | 0.0012 (**) | No HSA | 0.4341 (ns) | 0.0196 (*) | 0.0592 (ns) | 0.1192 (ns) | 0.4936 (ns) | No HSA | 0.4341 (ns) | 0.0196 (*) | 0.0592 (ns) | 0.1192 (ns) | 0.4936 (ns) |
| 1mg/ml HSA | 0.3575 (ns) | 0.0219 (*) | 0.0006 (***) | 0.0001 (***) | 0.0036 (**) | 1mg/ml HSA | 0.9320 (ns) | 0.7480 (ns) | 0.1101 (ns) | 0.5347 (ns) | 0.3542 (ns) | 1mg/ml HSA | 0.9320 (ns) | 0.7480 (ns) | 0.1101 (ns) | 0.5347 (ns) | 0.3542 (ns) |
| 7.5mg/ml HSA | 0.0087 (**) | 0.0053 (*) | 0.0042 (**) | 0.6187 (ns) | <0.0001 (***) | 7.5mg/ml HSA | <0.0001 (***) | <0.0001 (***) | <0.0001 (***) | <0.0001 (***) | <0.0001 (***) | 7.5mg/ml HSA | <0.0001 (***) | <0.0001 (***) | <0.0001 (***) | <0.0001 (***) | <0.0001 (***) |
| 25mg/ml HSA | 0.0102 (*) | 0.0167 (*) | 0.0047 (**) | <0.0001 (***) | <0.0001 (***) | 25mg/ml HSA | <0.0001 (***) | <0.0001 (***) | 0.001 (***) | <0.0001 (***) | 0.001 (***) | 25mg/ml HSA | <0.0001 (***) | <0.0001 (***) | 0.001 (***) | <0.0001 (***) | 0.001 (***) |
| N<sub>g</sub>=2 N<sub>m</sub>=2 and 3 |  |  |  |  |  | N<sub>g</sub>=2 N<sub>m</sub>=2 and 3 |  |  |  |  |  | Exe |  |  |  |  |  |
| KWEA |  |  |  |  |  | N<sub>g</sub>=2 N<sub>m</sub>=2 and 3 |  |  |  |  |  | Exe |  |  |  |  |  |
| Unpaired t test | CS0 | CS25 | CS50 | CS75 | CS100 | Unpaired t test | CS0 | CS25 | CS50 | CS75 | CS100 | Unpaired t test | CS0 | CS25 | CS50 | CS75 | CS100 |
| No HSA | 0.1914 (ns) | 0.6830 (ns) | 0.5988 (ns) | 0.1834 (ns) | 0.0574 (ns) | No HSA | 0.0145 (*) | 0.0167 (*) | 0.0039 (**) | 0.0012 (**) | 0.0007 (***) | No HSA | 0.0145 (*) | 0.0167 (*) | 0.0039 (**) | 0.0012 (**) | 0.0007 (***) |
| 1mg/ml HSA | 0.0082 (**) | 0.6118 (ns) | 0.4104 (ns) | 0.4257 (ns) | 0.8362 (ns) | 1mg/ml HSA | 0.3289 (ns) | 0.3803 (ns) | 0.5165 (ns) | 0.0927 (ns) | 0.1312 (ns) | 1mg/ml HSA | 0.3289 (ns) | 0.3803 (ns) | 0.5165 (ns) | 0.0927 (ns) | 0.1312 (ns) |
| 7.5mg/ml HSA | 0.0775 (ns) | 0.7381 (ns) | 0.1710 (ns) | 0.2317 (ns) | 0.0056 (**) | 7.5mg/ml HSA | 0.0130 (*) | 0.0370 (*) | 0.0233 (*) | 0.0428 (*) | 0.0115 (*) | 7.5mg/ml HSA | 0.0130 (*) | 0.0370 (*) | 0.0233 (*) | 0.0428 (*) | 0.0115 (*) |
| 25mg/ml HSA | 0.0055 (**) | 0.9277 (ns) | 0.0002 (***) | 0.0071 (**) | 0.0036 (**) | 25mg/ml HSA | 0.3131 (ns) | 0.4494 (ns) | 0.2747 (ns) | 0.4638 (ns) | 0.4677 (ns) | 25mg/ml HSA | 0.3131 (ns) | 0.4494 (ns) | 0.2747 (ns) | 0.4638 (ns) | 0.4677 (ns) |

While D increased by 15% at 7.5 mg/mL and by 23% at 25 mg/mL HSA in the CS0 hydrogel, the same HSA concentrations enhanced D by 76% and 107%, respectively, in the CS100 hydrogel. Because GK3 diffusion showed little dependence on hydrogel composition, time-dependent measurements were conducted in only three hydrogel types. These measurements revealed a trend similar to GK6, with D increasing significantly after 21 h by 35-60% at both HSA concentrations, independent of hydrogel composition or HSA concentration.

|  |  |  |  |  |  |
| --- | --- | --- | --- | --- | --- |
| $N_g=2$ | $N_m=2/3$ | GK6 | GK7 | | |
| Difference between gels at 25mg/ml hsa after 21h |  |  |  |  |  |
| Shapiro-Wilk test | CS0 | CS25 | CS50 | CS75 | CS100 |
| P value | 0.4985 | 0.4524 | 0.4053 | 0.5237 | 0.8537 |
| Passed normality test (alpha=0.05)? | Yes | Yes | Yes | Yes | Yes |
| P value summary | ns | ns | ns | ns | ns |
| Brown-Forsythe test |  | Bartlett's test |  |  |  |
| 0.2343 (ns) |  | 0.2454 (ns) |  |  |  |
| Ordinary one-way Anova | CS0 | CS25 | CS50 | CS75 |  |
| CS25 | 0.1207 (ns) | X | X | X |  |
| CS50 | 0.0746 (ns) | 0.9990 (ns) | X | X |  |
| CS75 | <0.0001 (****) | 0.0021 (**) | 0.0037 (**) | X |  |
| CS100 | 0.0050 (**) | 0.5649 (ns) | 0.7152 (ns) | 0.0582 (ns) |  |

|  |  |  |  |  |  |  |  |
| --- | --- | --- | --- | --- | --- | --- | --- |
| Difference between gels after 21h |  |  |  |  |  |  |  |
| 7.5mg/ml HSA |  | 25mg/ml HSA |  |  |  |  |  |
| Shapiro-Wilk test | CS0 | CS50 | CS100 | Shapiro-Wilk test | CS0 | CS50 | CS100 |
| P value | 0.9675 | 0.1835 | 0.5698 | P value | 0.2360 | 0.8867 | 0.1700 |
| Passed normality test (alpha=0.05)? | Yes | Yes | Yes | Passed normality test (alpha=0.05)? | Yes | Yes | Yes |
| P value summary | ns | ns | ns | P value summary | ns | ns | ns |
| Brown-Forsythe test |  | Bartlett's test |  | Brown-Forsythe test |  | Bartlett's test |  |
| 0.3980 (ns) |  | 0.3456 (ns) |  | 0.4650 (ns) |  | 0.1458 (ns) |  |
| Ordinary One-way Anova | CS0 | CS50 |  | Ordinary One-way Anova | CS0 | CS50 |  |
| CS50 | 0.0180 (*) | X |  | CS50 | 0.3994 (ns) | X |  |
| CS100 | 0.0011 (**) | 0.2957 (ns) |  | CS100 | 0.9653 (ns) | 0.2816 (ns) |  |

### In vivo experiments

**Figure S33:** Chemical structure of ABD-RM26' and its LC-MS analysis

**Figure S34:** SPR sensorgrams showing peptide binding to immobilized HSA. A and B) ABD-RM26', C and D) Semaglutide as control

**Table S22:** HSA dissociation constant determined based on SPR model fitting

| Peptide | $K_D$ (M) | $\chi^2$ |
| --- | --- | --- |
| ABD-RM26' | $9.7 \times 10^{-9} (\pm 7.7 \times 10^{-9})$ | $0.03 (\pm 0.009)$ |
| Semaglutide | $1.1 \times 10^{-5} (\pm 1.3 \times 10^{-6})$ | $3.5+ (\pm 0.2)$ |

**Figure S35:** Time-dependent activity changes at the injection site, kidney, and bladder, together with total activity changes in the whole body following subcutaneous injection of GK6, Exe, and RM26'.

**Figure S36:** SPECT/CT images of mice acquired immediately after injection and 60 minutes later after administration of GK6 and exenatide, with the animals being unsedated for 60 minutes between the two time points

### Supporting information for Materials and Methods

#### List of materials

##### Polymers and Biomolecules:

Type I collagen (COL) (100mg, various concentrations, Corning®, from rat tail tendon, Discovery Labware Inc., Bedford, MA), Sodium-Hyaluronate (HA) (200K-5, Lifecore Biochemical, Chaska, MN, USA), Chondroitin sulfate from bovine cartilage (CS) (Standard, Sigma Aldrich SE), Human serum albumin (HSA) (96%, Lyophilized powder, Sigma Aldrich SE), GK6 (96.89%, Biomatik), FITC-GK6 (98.46%, Biomatik), FITC-GK3 (95.8%, Biomatik), KWEA (98.27%, Biomatik), Exenatide (was provided by AstraZeneca).

##### Chemicals and Reagents:

Methacrylic anhydride (94%, Sigma Aldrich, SE), Lithium phenyl-2,4,6-trimethylbenzoylphosphinate (LAP) (Sigma Aldrich, SE), 1-ethyl-3-(3-dimethylaminopropyl)carbodiimide (EDC) (≥99%, Sigma Aldrich, SE), 3,3'-dithiobis(propanoic dihydrazide) (DTP) (Frontier Scientific, Newark, DE, USA), Dithiotreitol (DTT) (99%, Sigma Aldrich, SE), Phosphate-buffered saline (PBS) (Tablets, Sigma Aldrich, SE), Na<sub>2</sub>HPO<sub>4</sub> (≥99%, Bioreagent, Sigma Aldrich, SE), NaH<sub>2</sub>PO<sub>4</sub> (≥99%, ACS reagent, Sigma Aldrich, SE), NaOH (≥99%, pellets, Merck, SE), NaHCO<sub>3</sub> (≥99.7%, ACS reagent, Sigma Aldrich, SE), Na<sub>2</sub>CO<sub>3</sub> (99.95-100.5%, ACS reagent, Sigma Aldrich, SE), NaCl (≥99%, ACS reagent, Sigma Aldrich, SE), EDTA (99-101%, Sigma Aldrich, SE), Ellman's reagent (DTNB) (≥98%, Bioreagent, Sigma Aldrich, SE), N-acetylcysteine (≥99%, TLC, Sigma Aldrich, SE), Dimethylmethylene blue\*(ZnCl<sub>2</sub>)<sub>2</sub> (DMMB) (Technical grade, LGC/TRC, Germany), D<sub>2</sub>O (99.9% D, Sigma Aldrich, SE), DMSO (99.7%, for gas chromatography, Merck, SE), Acetic acid (99%, glacial, Reagent Plus, Sigma Aldrich, SE), HCl (37%, Reagent grade, Scharlab, Esp), FITC-Dextran (10, 40, 70, and 500kDa, 100%, Sigma Aldrich, SE), 70kDa dextran (TCI, China), Fluorescein 5(6)-isothiocyanate (FITC) (≥90%, Bioreagent, Sigma Aldrich, SE), Alexa Fluor™ 633 NHS ester (Fischer Scientific, SE), Tween 20 (Sigma Aldrich, SE), Sodium metabisulfite, Chloramine-T, Citric acid, Methanol were from Sigma Aldrich (St. Louis, MO, USA), Ammonium acetate buffer and Borate buffer were prepared from chemicals supplied by Sigma Aldrich (St. Louis, MO, USA), N-succinimidyl p-(trimethylstannyl)benzoate was synthesised according to published method<sup>[2]</sup>, [<sup>125</sup>I]NaI was from Perkin-Elmer Inc (Waltham, MA, USA), [<sup>111</sup>In]InCl<sub>3</sub> was from Curium Pharma (Stockholm, Sweden).

#### Supplementary Methods

These protocols provide additional experimental details supporting the **Materials and Methods** section of the main manuscript.

##### **Supplementary Method 1: Preparation of hydrogel precursor solution**

The required volume of the MAC solution was transferred into a 5 mL Eppendorf tube. HASH and CSSH solutions were combined in a 1.5 mL Eppendorf tube and gently mixed. The appropriate volumes of LAP solution and PBS were added to the HASH/CSSH mixture to achieve the desired final polymer concentrations as specified in the main Materials and Methods. The HASH/CSSH/LAP mixture was transferred into the MAC solution. The combined precursor solution was homogenized by brief vortexing followed by repeated pipetting using a Gilson Microman E M1000E pipette. Vortexing and pipetting were repeated once to ensure uniform mixing. A final brief vortexing step was performed to facilitate the removal of air bubbles from the precursor solution.

#### **Supplementary method 2: Synthesis of RM26 with a PEG-linker and ABD-RM26'**

The precursor peptides for RM26-PEG ($\text{H}_2\text{N-D-F-Q(Trt)-W(Boc)-A-V-Sar-H(Trt)-Sta-L}$ on Rink Amide resin) and ABD-RM26' ($\text{H}_2\text{N-D-F-Q(Trt)-}\alpha\text{metW(Boc)-A-V-Sar-H(Trt)-Sta-L}$ on Rink Amide resin) were provided by Pepmic Co., Ltd. (Suzhou, Jiangsu, China). Synthesis was conducted using manual Fmoc SPPS on a 10 $\mu\text{mol}$ scale in a 6 mL disposable syringe.

Synthesis of RM26-PEG commenced with pre-swelling of the precursor resin in DCM (0.5 mL) for 30 min, followed by washing with DMF (2x1 min, 0.5 mL). Fmoc-PEG2-OH (4 equiv.), Oxyma Pure (4 equiv.), DIPEA (7 equiv.) and DIC (4 equiv.) were dissolved in DMF (0.5 mL) and added to the resin. After 2 h, the resin was washed with DMF (2x1 min, 0.5 mL), and Fmoc-deprotection was done with 20% piperidine in DMF (3x10 min) followed by washing with DMF (6x1 min, 0.5 mL). Cleavage and purification of RM26-PEG were done as described below.

Synthesis of ABD-RM26' commenced with pre-swelling of the precursor resin in DCM (0.5 mL) for 30 min, followed by washing with DMF (2x1 min, 0.5 mL). Fmoc-4-Amino-1-Carboxymethyl-Piperidine (4 equiv.), Oxyma Pure (4 equiv.), DIPEA (7 equiv.) and DIC (4 equiv.) were dissolved in DMF (1 mL) and added to the resin. After 2 h, a second batch of the coupling cocktail was added, and the mixture was left for 16h. The resin was washed with DMF (10x1 min, 1.0 mL), and Fmoc-deprotection was done with 20% piperidine in DMF (3x10 min) followed by washing with DMF (6x1 min, 0.5 mL). Fmoc-PEG2-OH (4 equiv.), Oxyma Pure (4 equiv.), DIPEA (7 equiv.) and DIC (4 equiv.) were dissolved in DMF (0.5 mL) and added to the resin. After 2 h, the resin was washed with DMF (2x1 min, 0.5 mL), and Fmoc-deprotection was done with 20% piperidine in DMF (3x10 min) followed by washing with DMF (6x1 min, 0.5 mL). Fmoc-Lys(Alloc)-OH (4 equiv.), Oxyma Pure (4 equiv.), DIPEA (7 equiv.) and DIC (4 equiv.) were dissolved in DMF (0.5 mL) and added to the resin. After 2h, the resin was washed with DMF (2x1 min, 0.5 mL) and DCM (4x1 min). $\text{Pd(PPh}_3)_4$ (0.3 equiv.) and $\text{PhSiH}_3$ (20 equiv.) were dissolved in DCM and added to the resin for $2 \times 1$ h. The resin was then washed with DCM (3x1 min) and DMF (3x1 min). 4-Iodophenylbutyric acid (4 equiv.), Oxyma (4 equiv.), DIC (4 equiv.) and DIPEA (7 equiv.) were dissolved in DMF (0.5 mL) and added to the resin. After 2 h, the resin was washed with DMF (2x1 min, 0.5 mL), and Fmoc-deprotection was done with 20% piperidine in DMF (3x10 min) followed by washing with DMF (6x1 min, 0.5 mL). DOTA tris(tBu-ester) (4 equiv.), Oxyma (4 equiv.), DIC (4 equiv.) and DIPEA (7 equiv.) were dissolved in DMF (0.5 mL) and added to the resin. After 4h, the resin was washed with DMF (2x1 min, 0.5 mL) and DCM (4x1 min).

The resin was dried under vacuum for 30 min and treated with 0.5 mL of a cleavage solution consisting of TFA:$\text{H}_2\text{O}$:TIS (95:2.5:2.5). After 4h, the solution was filtered into a Falcon tube, and the resin was again washed with the cleavage solution $3 \times 250 \mu\text{L}$. To the solution, ice-cold $\text{Et}_2\text{O}$ (45 mL) was added to precipitate the crude peptide. The precipitate was washed with ice-cold $\text{Et}_2\text{O}$ ($3 \times 45$ mL), dried under vacuum, and redissolved in a minimal amount of ACN:$\text{H}_2\text{O}$. Purification was done with preparative LC-UV using a 10-40 ACN:$\text{H}_2\text{O}$ + 0.1% TFA gradient.

#### **Supplementary Method 3: Raman spectroscopy**

The laser beam was focused onto the polymer surface using a cylindrical lens. The resulting illuminated area was approximately 100 $\mu\text{m}$ in length and 10 $\mu\text{m}$ in width.

#### **Supplementary Method 4: Determination of thiol content by Ellman's assay**

A reaction buffer (pH 8.0) consisting of 0.1 M $\text{Na}_3\text{PO}_4$ and 1 mM EDTA was prepared. Ellman's reagent solution was prepared by dissolving DTNB at 4 mg $\text{mL}^{-1}$ in reaction buffer. Calibration standards were prepared according to the manufacturer's instructions using N-acetylcysteine instead of cysteine. For sample analysis, 1 mg of HASH or CSSH was dissolved separately in 10 mL nitrogen-flushed reaction buffer. After complete dissolution, 250 μL of the sample solution was mixed with 50 μL of Ellman's reagent and 2.5 mL of reaction buffer. Samples were incubated for 15 min at room temperature before measurement. Thiol concentrations were calculated from the calibration curve (Fig. S7) and used to determine the degree of thiol modification per milligram of polymer.

##### **Supplementary Method 5: FTIR spectroscopy**

Hydrogels were washed in excess Milli-Q water for 2 days with several changes prior to analysis to remove residual PBS. Raw spectra were corrected for CO<sub>2</sub>, H<sub>2</sub>O, ATR effects, and baseline using Jasco Spectra Manager software. Final spectra were processed and visualized using Origin 2018 software.

##### **Supplementary Method 6: SEM and EDS analysis**

Dried samples were mounted on SEM stubs using conductive adhesive tape. Au-Pd coating was applied using a Thermo VG Scientific Polaron SC7640 sputter coater for 50 seconds at a current of 20 mA. SEM imaging was performed on the outer surface of the hydrogels.

##### **Supplementary Method 7: DMMB assay**

The DMMB reagent solution was adjusted to pH 3 using 1 M HCl.

##### **Supplementary Method 8: Alexa Fluor 633 labeling of MAC**

Alexa Fluor 633 solution was prepared in DMSO at 1 mg mL<sup>-1</sup>. The dye solution was added dropwise to the MAC solution under constant stirring. Labeled MAC was precipitated by the addition of an equal volume of 1 M acetic acid / 1.3 M NaCl. The precipitate was centrifuged at 4,000 rcf for 1 h at 4 °C. The pellets were redissolved in 10 mM HCl (half of the original volume) and incubated overnight at 4 °C. The solution was dialyzed using a 3.5 kDa MWCO membrane (Spectra/Por 6, SpectrumLab, CA, USA) against the same NaOH solution used during MAC synthesis, with six buffer exchanges. The final product was frozen and lyophilized prior to use.

##### **Supplementary Method 9: FITC labeling of peptides and HSA**

Peptides and HSA were dissolved in 0.1 M carbonate buffer (pH 9.0) at concentrations of 2-5 mg mL<sup>-1</sup>. FITC was prepared in DMSO at a concentration of 5 mg mL<sup>-1</sup>. For HSA conjugates, dialysis was carried out for 3 days with four buffer exchanges prior to lyophilization.

##### **Supplementary Method 10: Microscale thermophoresis (MST)**

FITC-peptide solutions were prepared at concentrations ranging from 240 to 600 nM in assay buffer (PBS containing 0.05% Tween-20). HSA was prepared as a 1 mM stock solution in the same assay buffer. A 12-point, 1:1 serial dilution of HSA was prepared in a non-binding 384-well plate using assay buffer. Equal volumes of peptide solution were mixed with each HSA dilution and incubated for 10 min at room temperature. Excitation power was set to 10 % for Exenatide and GK3, and 20% for RM26, GK6, and P4. MST laser power was set to 40 %. Concentration ranges of the different peptides used in the measurements were $3.66 \times 10^{-4}$ - $1.88 \times 10^{-1}$ mM for GK6, $1.46 \times 10^{-4}$ - $3 \times 10^{-1}$ mM for GK3, $3.66 \times 10^{-4}$ - $1.88 \times 10^{-1}$ mM for KWEA, $2.44 \times 10^{-4}$ - $1.25 \times 10^{-1}$ mM for RM26, and $3.66 \times 10^{-4}$ - $9.37 \times 10^{-2}$ mM for Exe.

##### **Supplementary Method 11: Image processing and partition coefficient extraction**

Three-dimensional reconstructions were generated in Huygens Professional (v25.04) using either surface rendering or maximum intensity projection (MIP). Raw images were deconvolved using the “standard” strategy in the Deconvolution Wizard prior to segmentation and quantification. The ROI was transferred to the FITC channel by setting the seed value to 0 and copying the ROI from the Alexa 633 channel.

#### Supplementary Method 12: Rheological measurements and swelling analysis

Hydrogel discs (8 mm diameter, $130 \pm 10$ mg) were gently blotted to remove surface PBS prior to measurement. Samples were weighed before and after rheological testing using a Sartorius analytical balance. Gels were incubated in $7.5 \text{ mg mL}^{-1}$ dextran or HSA solutions for 48 h with two solution exchanges, followed by incubation in $25 \text{ mg mL}^{-1}$ solutions and repeat measurements. After the final measurement, gels were washed with Milli-Q water, frozen on dry ice, and lyophilized. The mass swelling ratio was calculated as the ratio of swollen to dry mass. The volumetric swelling ratio was calculated using the density of the swollen gel (assumed equal to PBS) and the theoretical density of the dry polymer.

#### Supplementary method 13: Image analysis and determination of the diffusion coefficient from gradient diffusion measurements

Here we represent examples of the selection of reference region during FRAP analysis. As the figures show, the selection of a reference region similar to the bleaching region results in better model fitting and slightly alters the diffusion coefficient. Therefore, as a selection criterion, the reference region was always selected at least from the bleaching region, at least at the same distance as the reference region size, and similar to the reference region.

**Figure S36:** FRAP analysis workflow of 40kDa dextran in CS50 hydrogel

**Figure S37:** Effect of the selection of the reference region on the diffusion coefficient of 25mg/mL 70kDa dextran in the polymer-rich domain of CS50 hydrogel measured by FRAP

**Figure S38:** Effect of the selection of the reference region on the diffusion coefficient of 25mg/mL HSA in the polymer-rich domain of CS100 hydrogel measured by FRAP

**Figure S39:** Effect of the selection of the reference region on the diffusion coefficient of GK6 in PBS in the polymer-rich domain of CS50 hydrogel measured by FRAP

**Figure S40:** Effect of the selection of the reference region on the diffusion coefficient of GK6 at 25 mg/mL HSA concentration in the polymer-rich domain of CS50 hydrogel measured by FRAP

**Figure S41:** Effect of the selection of the reference region on the diffusion coefficient of GK6 at 25 mg/mL HSA concentration in the polymer-rich domain of CS50 hydrogel after 21 hours of incubation, measured by FRAP

**Figure S42:** Effect of the HSA concentration on the diffusion coefficient of Exe in CS50 hydrogel measured by FRAP

##### Supplementary method 14: Image analysis and determination of the diffusion coefficient from gradient diffusion measurements

In most experiments, irregularities in the fluorescence signal were observed in the gel region close to the interface between the gel and the surrounding solution. Together with the fact that stirring of the donor solution could not be precisely controlled, and that aqueous boundary layer effects were present at the gel-solution interface, this prevented reliable definition of the upper boundary condition using simple physical assumptions.

Therefore, the lower boundary for solving the diffusion equation was defined at a position within the gel sufficiently far from the interface, where such edge effects were visually absent. For this lower boundary, a time-dependent concentration profile was applied in the form of a continuous, piecewise linear function. This function was constructed from experimentally determined fluorescence intensities by fitting linear segments over intervals of five consecutive time points. Continuity of the overall function was enforced across all segments.

The fitting procedure was considered converged when the relative difference in the sum of squared errors between two successive iterations was less than $10^{-6}$.

##### Supplementary Method 15A: $^{125}\text{I}$ labeling of GK6 and Exenatide

*N*-succinimidyl-*p*-(trimethylstannyl)benzoate (5 $\mu\text{L}$, 1 $\text{mg mL}^{-1}$ in 5% acetic acid in methanol) was added to a mixture of 10 $\mu\text{L}$ 0.1% acetic acid and 10 $\mu\text{L}$ $^{125}\text{I}$ stock solution, final pH 5.0. Iodination was initiated by the addition of chloramine-T (20 $\mu\text{L}$, 4 $\text{mg mL}^{-1}$ in water) and allowed to proceed for 5 min at room temperature. The reaction was quenched by the addition of sodium metabisulfite (20 $\mu\text{L}$, 6 $\text{mg mL}^{-1}$). Peptide solution (30 $\mu\text{L}$, 2 $\text{mg mL}^{-1}$ in water) was added, and pH was adjusted to 9.0 using borate buffer (70 $\mu\text{L}$, 0.07 M, pH 9.3). After incubation for 30 min at room temperature, radiolabeled peptides were purified using NAP-5 size-exclusion columns eluted with PBS. Radiochemical purity was analyzed by ITLC using silica gel strips (Tec-Control, Biodex).

#### **Supplementary Method 15B: $^{111}\text{In}$ labeling of ABD-RM26'**

ABD-RM26' (10 nmol in 10 $\mu\text{L}$ Milli-Q water) was buffered with 80 $\mu\text{L}$ ammonium acetate buffer (0.2 M, pH 5.5). $^{111}\text{In}$ (60 MBq in 80-150 $\mu\text{L}$ of 0.05 M HCl) was added to the reaction mixture. Labeling was performed for 10 min at 90 $^{\circ}\text{C}$. Radiochemical purity was analyzed by ITLC using silica gel strips (Tec-Control, Biodex).

#### **Supplementary Method 16: SPECT/CT acquisition and analysis**

Small-animal SPECT/CT imaging was performed to visualize the *in vivo* distribution of the [$^{125}\text{I}$] tracer at multiple time points post-injection (T1-T5 p.i.) using the nanoScan<sup>®</sup> SPECT/CT system (Mediso Medical Imaging Systems Ltd., Hungary). Animals were anesthetized with sevoflurane (3-4%) delivered in a 50:50 mixture of medical oxygen and air at a flow rate of 450 mL/min.

A whole-body CT scan was first acquired prior to radiotracer injection for anatomical reference with the following parameters: semicircular field of view (FOV), 480 projections, X-ray tube settings of 50 kVp and 610 $\mu\text{A}$, binning of 1:4, and an acquisition time of 5 minutes. Subsequently, SPECT imaging was performed over the same scan range as the CT acquisition at each time point (T1-T5 p.i.), using an energy window centered at 28.4 keV for [$^{125}\text{I}$].

During imaging, animals were maintained on a heated bed to prevent hypothermia. CT data were reconstructed using filtered back projection. SPECT data were reconstructed using Nucline software (version 3.05.031.000) with the Tera-Tomo 3D algorithm (3 subsets, 12 iterations), including corrections for scatter and attenuation. The reconstructed SPECT and CT DICOM datasets were co-registered and analyzed using Interview<sup>™</sup> FUSION software (version 3.11.009.000). A CT scan was acquired prior to radiotracer injection for anatomical reference. VOI selection was performed automatically, but the selected volume was reduced by 1 step to make more reasonable selections.

### **Theoretical account of HSA's effect on GK6 partitioning between polymer-rich domains and voids**

The objective of the thermodynamic model is to calculate the partition coefficient of the peptide between these domains as a function of HSA concentration. To this end, we present the thermodynamic framework used to determine the partition coefficient, along with models describing the free energy of complex formation between the peptide and HSA, chondroitin sulfate (CS), and hyaluronic acid (HA), respectively. We modelled the system as a finite-volume hydrogel comprising a polymer-rich domain and a void, in equilibrium with a surrounding liquid phase. The polymer-rich domain was treated as a separate phase, whereas the voids were assumed to be polymer-free and therefore part of the same phase as the external solution, consistent with the experimental observations.

Chemical potentials for all mobile species were derived for both phases, taking into account the following binding equilibria: GK6/HSA, GK6/polyelectrolyte, HSA/polyelectrolyte, and GK6/HSA/polyelectrolyte. The latter describes binding of GK6/HSA complexes to the polyelectrolyte and was assumed to follow the same equilibrium constant as HSA/polyelectrolyte binding. Equilibrium concentrations were obtained by equating the chemical potentials of electroneutral components across all phases. Because the gel volume and the volume fraction of the polymer-rich phase were fixed, water partitioning was not considered.

The experimentally determined dissociation constant $K_D$ was used to describe GK6/HSA binding in both phases. The equilibrium constants for GK6-polyelectrolyte and HSA-polyelectrolyte binding were determined by fitting the model to reproduce the experimentally measured partition coefficients obtained in separate experiments with only one binding species present.

### Thermodynamic model

We model the heterogeneous ECM hydrogel as comprising a rigid polymer-rich domain, occupying a volume fraction $\phi$, and a polymer-lean phase (void). Collagen, chondroitin sulfate (CS), and hyaluronic acid (HA) are assumed to be confined exclusively to the polymer-rich domain. The gel, with volume $V_g$, is in equilibrium with an external solution of volume $V_s$ containing water, monovalent salt ions (+, -), human serum albumin (HSA), peptide (pep), and HSA-peptide complexes (HSA:pep). The equilibrium distribution of all mobile electroneutral solutes is determined by imposing the condition of uniform chemical potential throughout the system.

The chemical potentials in the void ($v$) are:

$$\mu_i^v = \mu_i^{0,w} + kT \ln \frac{C_i^v}{C_0} \quad i \in \{HSA, pep, +, -, HSA:pep\} \quad (1)$$

where $\mu_i^{0,w}$ is the standard chemical potential in water (mole fraction scale), $kT$ is the thermal energy, $C_i^v$ is the molar concentration, and $C_0$ is the concentration of water (55.5 M).

In the polymer-rich domains ($p$), the species are distributed between two states: freely dispersed ($f$) or bound to negatively charged polyion chains ($b$). The chemical potential of a free species in the polymer-rich domains is:

$$\mu_i^{p,f} = \mu_i^{0,w} - kT \ln K_{Ob} + kT \ln \frac{C_i^{p,f}}{C_0} \quad i \in \{HSA, pep, HSA:pep\} \quad (2)$$

The second term on the r.h.s. of eq. (2) is the contribution from steric interaction with the gel network, where $K_{Ob}$ is the obstruction factor according to Ogston's theory<sup>[3]</sup> of fiber networks:

$$K_{Ob} = \exp \left\{ -\phi \left( 1 + \frac{r_h}{r_f} \right)^2 \right\} \quad (3)$$

$\phi$ is the polymer volume fraction in the gel, $r_h$ is the solute's hydrodynamic radius, and $r_f$ is the fiber radius. When applying eq. (2), we set $r_f$ equal to the weighted average of the radii of collagen triple helices and CS/HA chains.

For the polymer-bound species, the chemical potentials are given by:

$$\mu_i^{p,b} = \mu_i^{0,b} + kT \ln \frac{\theta_i}{1 - \sum \theta} \quad i \in \{HSA, pep, HSA:pep\} \quad (4)$$

where $\mu_i^{0,b}$ is the standard chemical potential, $\theta_i$ is the fraction of 'sites' on the polyions occupied by species $i$, and $\sum \theta$ is the total fraction of sites occupied by all bound species. A site is defined by the number $N$ of polyion charges within the polymer segment associated with a bound molecule, such that $\theta_i = C_i^{p,b} / (C_p / N)$, where $C_i^{p,b}$ is the concentration of bound species $i$, and $C_p$ is the concentration of polymer charged groups in the polymer-rich phase. For a single binding species, Eq. (4) reduces to the Langmuir binding isotherm.

The chemical potentials of small ions in the polymer-rich phase are given by:

$$\mu_i^p = \mu_i^{0,w} + kT \ln \frac{C_i^p}{C_0} \quad i \in \{+, -\} \quad (5)$$

### Equilibrium conditions

The binding equilibrium between free peptide and free HSA is described by the dissociation constant $K_D$, available from experiments (Table 1). Assuming 1:1 the stoichiometry of the complex, we get:

$$K_D = \frac{C_{pep}^v C_{HSA}^v}{C_{pep:HSA}^v} = \frac{K_{Ob,pep}^{-1} C_{pep}^p C_{HSA}^p}{C_{pep:HSA}^p} \quad (6)$$

where $K_{Ob,pep}^{-1}$ accounts for the non-ideality of the free peptide solution in the polymer-rich phase. The corresponding factors for HSA and HSA:pep are assumed to be identical and therefore cancel out. The condition for equilibrium between free and polymer-bound species in the polymer-rich phase is given by:

$\mu_i^{p,f} = \mu_i^{p,b}$. Combining that with eqs. (2) and (4), we get:

$$C_i^{p,f} = \frac{K_{Ob,i}C_0}{K_{b,i}} \left( \frac{\theta_i}{1-\sum \theta} \right) \quad i \in \{HSA, pep\} \quad (7a)$$

$$K_{b,i} = \exp\{-(\mu_i^{0,b} - \mu_i^{0,w})/kT\} \quad (7b)$$

The corresponding relationship describing the equilibrium between free peptide and HSA-peptide complexes bound to the polymer network is:

$$C_{pep}^{p,f} = \frac{K_{Ob}C_0}{K_b} \left( \frac{\theta_{HSA:pep}}{1-\sum \theta} \right) \quad (8)$$

In the calculations, the binding constant $K_b$ is evaluated from the experimentally determined molar dissociation constant $K_D = C_0/K_b$.

#### Electroneutrality conditions

The volumes of the polymer-rich domains and voids are sufficiently large to be treated as separate phases. Electroneutrality within the void and polymer-rich phases, respectively, requires:

$$|Z_{HSA}|C_{HSA}^{v,tot} + C_-^v - Z_{pep}C_{HSA}^{v,tot} - C_+^v = 0 \quad (9)$$

$$|Z_{HSA}|C_{HSA}^{p,tot} + C_p + C_-^p - Z_{pep}C_{HSA}^{p,tot} - C_+^p = 0 \quad (10)$$

#### Conservation of mass

Mass conservation yields, taking into account the finite volumes of both the liquid solution and the gel:

$$C_i = C_i^{p,tot} \varphi \frac{V_g}{V_s} + C_i^{v,tot} (1 + (1 - \varphi) \frac{V_g}{V_s}) \quad (11)$$

where $C_i$ is the initial concentration of species $i$ in the liquid solution in contact with the gel. Equation (11) assumes that the equilibrium concentrations of all mobile species are identical in the liquid phase and the void. Because each HSA molecule can bind at most one peptide, the following relation holds in the void:

$$C_{pep}^{v,tot} = C_{pep}^v + C_{HSA:pep}^v \quad (12)$$

$$C_{HSA}^{v,tot} = C_{HSA}^v + C_{HSA:pep}^v \quad (13)$$

and for the polymer-rich phase:

$$C_{pep}^{p,tot} = C_{pep}^{p,f} + C_{pep}^{p,b} + C_{HSA:pep}^{p,f} + C_{HSA:pep}^{p,b} \quad (14)$$

$$C_{HSA}^{p,tot} = C_{HSA}^{p,f} + C_{HSA}^{p,b} + C_{HSA:pep}^{p,f} + C_{HSA:pep}^{p,b} \quad (15)$$

#### Phase equilibrium conditions

From the requirement of uniform chemical potential of electroneutral species throughout the system, the partitioning of freely dispersed HSA, peptide, and HSA-peptide complexes between the polymer-rich phase and the void is given by:

$$\frac{C_{pep}^{p,f}}{C_{pep}^v} = K_{Ob,pep} \left( \frac{C_+^p}{C_-^v} \right)^{Z_{pep}} \quad (16)$$

$$\frac{C_{HSA}^{p,f}}{C_{HSA}^v} = K_{Ob,HSA} \left( \frac{C_+^p}{C_-^v} \right)^{|Z_{HSA}|} \quad (17)$$

$$\frac{C_{HSA:pep}^{p,f}}{C_{HSA:pep}^v} = K_{Ob,HSA} \left( \frac{C_+^p}{C_-^v} \right)^{|Z_{HSA}-Z_{pep}|} \quad (18)$$

where the obstruction factor is assumed to be the same for HSA and HSA:pep.

Electrostatic effects arising from counterion binding are accounted for in the calculation of the free energy of complex formation in the polymer-rich phase. However, for simplicity, counterion binding is neglected when describing partitioning between the phases. Accordingly, the distribution of small ions is assumed to be governed by:

$$C_+^p C_-^p = C_+^v C_-^v \quad (19)$$

Furthermore, because salt is present in large excess, it is a good approximation to set:

$C_+^v = C_-^v = C_{salt}$. Eq. (19) in combination with eq. (9) gives:

$$C_+^p = -\frac{1}{2} (Z_{pep} C_{pep}^{p,tot} - |Z_{HSA}| C_{HSA}^{p,tot} - C_p) + \sqrt{\frac{1}{4} (Z_{pep} C_{pep}^{p,tot} - |Z_{HSA}| C_{HSA}^{p,tot} - C_p)^2 + C_{salt}^2} \quad (20)$$

$$C_-^p = C_+^p + Z_{pep} C_{pep}^{p,tot} - |Z_{HSA}| C_{HSA}^{p,tot} - C_p \quad (21)$$

#### Partition coefficients

To calculate the partition coefficients of the peptide and HSA in the mixed systems, the thermodynamic model was used to determine $C_{HSA}^{p,tot}$, $C_{HSA}^{v,tot}$, $C_{pep}^{p,tot}$, and $C_{pep}^{v,tot}$ by solving the system of equations (Eqs. 1-21) for predefined values of the parameter set $\{K_{b,HSA}, K_{b,pep}, K_D, C_{pep}, C_{salt}, C_p, \emptyset, \phi\}$. The partition coefficients were then calculated as:

$$K_{HSA} = \frac{C_{HSA}^{p,tot}}{C_{HSA}^{v,tot}} \quad (22)$$

$$K_{pep} = \frac{C_{pep}^{p,tot}}{C_{pep}^{v,tot}} \quad (23)$$

For $K_D$, we used the experimentally determined value for GK6 (Table 1). The binding constants $K_{b,HSA}$ and $K_{b,pep}$ were obtained by fitting the thermodynamic model to reproduce the experimentally determined partition coefficients of HSA and the peptide, measured in separate experiments. The resulting values are summarized in Table 2.

#### Molecular thermodynamic modelling

To theoretically account for the values of the binding constants $K_{b,HSA}$ and $K_{b,pep}$, a molecular thermodynamic approach was used to calculate the free energy associated with the binding of HSA and GK6 to chondroitin sulfate (CS) and hyaluronic acid (HA) chains.

*Polyelectrolyte-HSA complexation.* As a prototype system, we considered the complexation between HSA and short polyacrylate (PA) chains, for which detailed information is available in the literature. Shun and coworkers identified the dominant PA binding site on HSA using molecular dynamics (MD) simulations and showed that the calculated free energy of complexation was in good agreement with binding constants determined by isothermal titration calorimetry at different salt concentrations<sup>[4]</sup>. The binding site appears as a groove on one side of the protein, allowing the polyion chain to adopt a stretched configuration while interacting with multiple basic amino acid residues.

In their MD simulations, Shun and coworkers modeled the PA chain as 25 charged beads with a radius of 2 Å, connected by partially angular-restricted harmonic bonds. This resulted in a flexible chain with an average bond length of 4 Å, corresponding to an average charge spacing of $b=3.5$ Å along the chain. Based on the average configuration of the PA chain within the binding site, we estimated that the PA-HSA complex involves approximately 6 net positive protein charges and 15 negative polyion charges, corresponding to a patch length $L=52.5$ Å.

The MD simulations further showed that the PA chain enters the binding site via a mechanism resembling reptation in polymer solutions. In contrast, CS and HA chains are covalently attached to the network at both ends and therefore cannot access the entire binding site in the same way. To account for this limitation, the effective accessible length $L$ was adjusted such that the partition coefficient $K_{HSA}$ for CS100, calculated using the thermodynamic model, matched the experimentally determined value. The same value of $L$ was subsequently used in the modelling of CS0-HSA complexes. The model parameters used in the calculations are summarized in Table S2.

We considered two contributions to the electrostatic free energy of the complex: one arising from the net charge of the complex $G_{compl}^{PB}$ and one from the spatial separation of polyion and protein charges within the complex $G_{compl}^{cap}$. The term $G_{compl}^{PB}$ was calculated using Poisson-Boltzmann (PB) theory for a cylinder of radius $r_2$ with a uniform surface charge density corresponding to the net charge of the complex per unit length. The term $G_{compl}^{cap}$ was evaluated as the energy of a cylindrical capacitor, with an inner charged surface of charge density $\sigma$ and radius $r_1$, and an outer charged surface of radius $r_2$. The expression for the energy per unit area of a cylindrical capacitor is:

$$\frac{E}{area} = \frac{\sigma^2 r_1 \ln(r_2/r_1)}{2\epsilon_0 \epsilon_r} \quad (24)$$

where $\epsilon_0$ is the permittivity of vacuum, $\epsilon_r$ is the dielectric constant of the medium. From eq. (24) we obtain the following expression

$$\frac{G_{compl}^{cap}}{kT} = \frac{l_B Z_{cap}^2 \ln(r_2/r_1)}{L} \quad (25)$$

where $l_B = e^2/(4\pi\epsilon_0\epsilon_r kT)$ is the Bjerrum length. In the calculations, we set $T=307$ (34 °C) and $\epsilon_r=20$, the latter being a typical value used in the literature to describe electrostatic interactions at water-poor sites near protein surfaces<sup>[5]</sup>.

Poisson-Boltzmann (PB) theory was also used to calculate the electrostatic free energy of the free polyion chain $G_{pol}^{PB}$ and the electrostatic free energy of the empty surface patch $G_{patch}^{PB}$. The polyion was modeled as a charged cylinder with radius $r_{pol}$ and area per surface charge $a_{pol} = 2\pi r_{pol}b$, while the empty surface patch was represented as a charged cylinder with radius $r_2$ and area per surface charge $a_{patch} = 2\pi r_2 L/Z_{patch}$. For these calculations, we used $\epsilon_r=78.5$, corresponding to water at $T=307$.

The electrostatic free energy change associated with the binding of the polyion to the patch is then given by:

$$\Delta G^0 = G_{compl}^{PB} + G_{compl}^{cap} - G_{pol}^{PB} - G_{patch}^{PB} \quad (26)$$

All Poisson-Boltzmann (PB) calculations were performed using the PBcell software developed by Bengt Jönsson (Computer program written by Prof. Bengt Jönsson, Lund University, 1998.). The free energy of a cylinder with finite length $L$ was approximated as the free energy per unit length of an infinitely long cylinder (with identical radius and surface charge density) multiplied by $L$, thereby neglecting end effects.

To calculate the binding equilibrium constant, we used:

$$K_{b,HSA} = \exp\{-(\mu_{iHSA}^{0,b} - \mu_{HSA}^{0,w})/kT\} \equiv \exp\{-\Delta G^0/kT\} \quad (27)$$

#### Polyelectrolyte-peptide complexation

For the GK6 peptide, the complex is assumed to consist of one peptide molecule with contour length $l$ and charge $Z_{pep} = l/b_{pep}$, and a segment of the polyion with length $L$ and charge $Z_{pol} = L/b_{pol}$. Here, $b_{pep}$ denotes the spacing between charges along the peptide in its stretched conformation, and $b_{pol}$ is the corresponding spacing along the polyion chain. The net charge of the complex is then given by:

$$Z_{compl} = Z_{pep} - Z_{pol} = Z_{pep} \left(1 - \frac{L/b_{pol}}{l/b_{pep}}\right) \quad (28)$$

The peptide is assumed to make $n$ helical turns around the polyion chain, therefore:

$$n = \frac{L}{2\pi p} \quad (29)$$

$$\frac{L}{l} = \frac{p}{\sqrt{p^2 + r_2^2}} \quad (30)$$

where $2\pi p$ is the pitch and $r_2$ is the radius of the peptide helix. The structural basis for this helical conformation is supported by molecular dynamics simulations of HA-peptide complexes by Kulik et al<sup>[6]</sup>, which show that the peptide folds around the polyion in a twisted configuration.

As in the case of polyion-HSA complexation (see above), the electrostatic free energy of the complex is expressed as $G_{compl}^{PB} + G_{compl}^{cap}$. The first term represents the electrostatic free energy of a uniformly charged cylinder with charge $Z_{compl}$ and radius $r_2$, calculated using the cylindrical Poisson-Boltzmann (PB) model. The second term accounts for the electrostatic contribution arising from the separation between the negative charges on the polyion and the peptide within the helical structure, and is calculated using Eq. (25), where $r_1$ is the radius of the polyion chain.

The electrostatic free energies of the free polyion and free peptide were calculated using the cylindrical PB model as described above. In these calculations, we set $\epsilon_r = 78.5$. The standard free energy change $\Delta G^0$ was obtained from Eq. (26), and the binding equilibrium constant was subsequently calculated using Eq. (31).

$$K_{b,pep} = \exp\{-\Delta G^0/kT\} \quad (31)$$

All parameters used in the calculations are presented in Table S22.

**Table S22.** Molecular thermodynamic modelling. Parameters defining the systems, calculated binding free energies ($\Delta G^0$), binding constants ($K_b$), partition coefficients ($K(calc.)$), and experimental partition coefficients ($K(exp.)$) at 307 K.

|  | <u>PA-HSA</u> | <u>CS100-HSA</u> | <u>CS0-HSA</u> | <u>CS100-GK6</u> | <u>CS0-GK6</u> |
| --- | --- | --- | --- | --- | --- |
| $Z_{patch}$ | +6 | +5.2 | +5.2 | - | - |
| $Z_{pol}$ | -15 | -9.0 | -4.5 | -10 | -6.5 |
| $Z_{pep}$ | - | - | - | +5 | +5 |
| $L$ (Å) | 53 | 45 | 45 | 51 | 66 |
| $l$ (Å) | - | - | - | 66 | 66 |
| $b_{pol}$ (Å) | 3.5 | 5 | 10 | 5 | 10 |
| $b_{pep}$ (Å) | - | - | - | 13 | 13 |
| $r_{pol}$ (Å) | 3-3.5 | 5 | 5 | 5 | 5 |
| $r_{compl}$ (Å) | 6 | 6 | 6 | 7 | 7 |
| $r_1$ (Å) | 2.5-3 | 5 | 5 | 5 | 5 |
| $r_2$ (Å) | 6 | 6 | 6 | 7 | 7 |
| $-\Delta G^0/kT$ | 5.5-11 | 11.0 | 8.05 | 10.9 | 5.24 |
| $K_b/10^3$ | 0.24-60 | 58 | 3.1 | 54 | 0.19 |
| $K(calc.)$ | - | 1.1<sup>a</sup>, 0.83<sup>b</sup> | 0.49<sup>a</sup>, 0.44<sup>b</sup> | 2.7 | 1.1 |
| $K(exp.)$ | - | 1.1<sup>a</sup>, 0.94<sup>b</sup> | 0.95<sup>a</sup>, 0.86<sup>b</sup> | 2.7 | 1.7 |

a) 1 mg/mL HSA

b) 25 mg/mL HSA

### Statistical Validation of Hydrogel Sampling Methodology: Intra-Gel Heterogeneity vs. Batch-to-Batch Variability

#### Rationale and Theoretical Framework

To evaluate whether sampling multiple positions within a single hydrogel specimen underestimates experimental variance (pseudoreplication), a systematic variance components analysis was conducted. The central hypothesis assumes that local micro-heterogeneity, arising from structural domain variations within a single gel piece, is the dominant contributor to variance in measured partition coefficients ($K$) and diffusion coefficients ($D$), rather than variance introduced by independent gel preparation batches prepared at different time points. Under this hypothesis, sampling from separately prepared hydrogel batches of identical composition is not expected to yield greater variance than sampling from multiple spatially distinct regions within a single hydrogel piece. To rigorously test this hypothesis, variance ratios (F-values) were evaluated:

$$F = \frac{s_{between}^2}{s_{within}^2}$$

where $s_{between}^2$ represents the variance measured between independently prepared hydrogel batches, and $s_{within}^2$ represents the variance measured across multiple sampling sites within a single hydrogel piece. An F-value < 1.0 indicates that spatial intra-gel variance exceeds inter-batch variance, demonstrating that sampling within a single gel piece provides a conservative estimate of overall experimental error and conversely.

Under the null hypothesis, i.e., $s_{within}^2 = s_{between}^2$, the obtained F-values from the variance tests are, due to randomness, supposed to follow an F-distribution. In the case of just one test, the reported p-value, used for the null hypothesis assessment, is the cumulative probability, under the null hypothesis, of finding an F-value larger than the observed.

In this case, we do several variance tests, and with that, the cumulative probability of the obtained F-values is, under the null hypothesis, expected to be close to the corresponding cumulative F-distribution given by the null hypothesis. A p-value for the comparison of the obtained and the hypothesized cumulative distributions was determined using a Kolmogorov-Smirnov (K-S) goodness-of-fit test<sup>[7]</sup>. The test statistic used was the maximum distance between distributions ($Diff_{max}$) and the corresponding p-value was calculated as follows:

$$p = 2 \sum_{j=1}^n -1^{j-1} e^{-2(j \cdot \lambda)^2} \quad \text{where } \lambda = \left( \sqrt{N} + 0.11 + \frac{0.12}{\sqrt{N}} \right) \cdot Diff_{max}$$

where $N$ is the total number of observed F-values and $n = 10$ ensures numerical convergence of the sum.

##### Human Serum Albumin (HSA) Diffusion and Partitioning Coefficient Variance Analysis

PBS and hydrogel batches across five compositions (CS0, CS25, CS50, CS75, CS100) were prepared, and the variances of the diffusion (D) and partition (K) coefficients for human serum albumin (HSA) at three concentrations (1, 7.5, and 25 mg/ml) were determined across three independent preparations.

The individual F-tests for HSA diffusion coefficients yielded F-values ranging from 0.03 to 8.05 (Table SX). Only 6 out of 18 conditions (33%) exhibited $F > 1.0$, indicating that in two-thirds of the observed intra-gel spatial variances were greater than corresponding inter-batch variances. The cumulative distribution function showed strong agreement with the cumulative $F_{2,4}$-distribution function (K-S test $p = 0.574$; Figure SX), confirming no statistically significant difference between intra-gel and batch-to-batch variance.

**Table S23.** Variance ratios (F-values) and associated p-values for HSA diffusion coefficients (D). Individual F-tests comparing between-batch variance to within-gel variance across PBS and five hydrogel formulations (CS0, CS25, CS50, CS75, CS100) and three HSA concentrations (1, 7.5, and 25 mg/ml). Values are reported as F-value : p-value.

|  | PBS | CS0 | CS25 | CS50 | CS75 | CS100 |
| --- | --- | --- | --- | --- | --- | --- |
| **1.0mg/ml** | 1.50 ; 0.33 | 5.73 ; 0.07 | 8.05 ; 0.04 | 0.63 ; 0.58 | 0.75 ; 0.53 | 0.40 ; 0.70 |
| **7.5mg/ml** | 1.09 ; 0.42 | 0.22 ; 0.81 | 2.36 ; 0.21 | 0.21 ; 0.82 | 0.03 ; 0.97 | 0.08 ; 0.92 |
| **25mg/ml** | 1.67 ; 0.30 | 0.04 ; 0.96 | 0.54 ; 0.62 | 0.24 ; 0.80 | 0.11 ; 0.90 | 0.14 ; 0.87 |

**Figure S43.** Cumulative distribution function (CDF) of F-values (blue dots) from variance tests comparing within-gel variance to between-batch variance (N = 18) plotted against the theoretical $F_{2,4}$-distribution (red solid line).

For HSA partition coefficients (K), 14 out of 15 conditions exhibited $F < 1.0$ (Table SX), suggesting that spatial micro-heterogeneity within individual gel pieces consistently exceeds batch-to-batch variation for equilibrium partitioning.

**Table S24.** Variance ratios (F-values) and associated p-values for HSA partition coefficients (K). Individual F-tests comparing between-batch variance to within-gel variance across five hydrogel formulations and three HSA concentrations. Values are reported as F-value : p-value.

|  | CS0 | CS25 | CS50 | CS75 | CS100 |
| --- | --- | --- | --- | --- | --- |
| **1.0mg/ml** | 1.634 ; 0.303 | 0.550 ; 0.615 | 0.315 ; 0.746 | 0.135 ; 0.877 | 0.208 ; 0.820 |
| **7.5mg/ml** | 0.140 ; 0.874 | 0.438 ; 0.673 | 0.234 ; 0.801 | 0.107 ; 0.901 | 0.564 ; 0.608 |
| **25mg/ml** | 0.189 ; 0.835 | 0.992 ; 0.447 | 0.332 ; 0.735 | 0.576 ; 0.603 | 0.007 ; 0.993 |

##### Dextran Diffusion and Partitioning Coefficient Variance Analysis

To assess the generality of this behavior across neutral macromolecules, Dextran standards of varying molecular weights (10, 40, 70, and 500kDa) were evaluated in CS0, CS50, and CS100 hydrogels.

A variance test, including an older (previous) hydrogel batch, showed elevated F-values for Dextran diffusion coefficients (9 out of 12 tests with $F > 1.0$; Table S3), and a K-S test of the cumulative distribution relative to the $F_{2,5}$-distribution showed a $p = 0.004$ (Figure SX). However, when evaluating variance across different gel compositions prepared within the same timeframe (Table SX), the K-S test $p = 0.895$ strongly indicates that the null hypothesis holds, i.e., the between-batch variance is not higher than the within-batch variance (Figure SX).

**Table S25.** Variance ratios (F-values) and associated p-values for dextran diffusion coefficients (D) across batches. Individual F-tests comparing between-batch variance to within-gel variance across three hydrogel formulations (CS0, CS50, CS100), including historical batch comparison. Values are reported as F-value : p-value.

|  | CS0 | CS50 | CS100 |
| --- | --- | --- | --- |
| **10 kDa** | 2.85 ; 0.15 | 12.96 ; 0.01 | 7.37 ; 0.03 |
| **40 kDa** | 5.85 ; 0.05 | 0.50 ; 0.63 | 1.14 ; 0.39 |
| **70 kDa** | 20.55 ; 0.00 | 0.52 ; 0.62 | 0.02 ; 0.98 |
| **500 kDa** | 5.43 ; 0.06 | 18.10 ; 0.01 | 11.84 ; 0.01 |

**Figure S44.** Cumulative distribution function (CDF) of F-values (blue dots) from variance tests comparing within-gel variance to between-batch variance (N = 12) plotted against the theoretical $F_{2,5}$-distribution (red solid line).

The p-value ($p = 0.004$) indicated that potential collagen batch variability causes a discrepancy of diffusion coefficients for at least one of the gels relative to the other gels. To directly address whether short-term batch preparation introduces significant variation, a follow-up prospective study was performed comparing two newly, independently prepared hydrogel batches. When variance testing these contemporary batches, 50% of F-values were $\leq 1.0$ (Table S6), and the obtained cumulative distribution function matched the theoretical $F_{2,3}$-distribution function with exceptional alignment (K-S test $p = 0.9997$; Figure SX). This suggests that hydrogels prepared under consistent operational conditions exhibit no additional inter-batch variance beyond native intra-gel micro-heterogeneity. The discrepancy observed with historical batches (Table SX) is attributed to variability between collagen batches due to its biological source rather than batch-to-batch preparation irreproducibility. It is important to mention that hydrogel batches used in one experimental setting were prepared from the same collagen batches.

**Table S26.** Variance ratios (F-values) for dextran diffusion coefficients (D) obtained from variance tests of hydrogel batches prepared at the same time. Comparison of variances between two independently synthesized hydrogel preparations. Values are reported as F-value : p-value.

|  | CS0 | CS50 | CS100 |
| --- | --- | --- | --- |
| 10 kDa | 1.18 ; 0.36 | 0.02 ; 0.89 | 1.36 ; 0.33 |
| 40 kDa | 0.38 ; 0.58 | 0.44 ; 0.56 | 0.88 ; 0.42 |
| 70 kDa | 27.09 ; 0.01 | 1.26 ; 0.34 | 0.03 ; 0.88 |
| 500 kDa | 0.35 ; 0.60 | 2.88 ; 0.19 | 13.30 ; 0.04 |

**Figure S45.** Cumulative distribution function (CDF) of F-values (blue dots) from variance tests comparing within-gel variance to between-batch variance (N = 12) plotted against the theoretical $F_{2,3}$-distribution (red solid line).

For Dextran partition coefficients (K), 11 out of 12 tests exhibited $F < 1.0$ (Table S5), suggesting that spatial heterogeneity within individual gel pieces dominates over batch-to-batch variance.

**Table S27.** Variance ratios (F-values) for dextran partition coefficients (K). Individual F-tests comparing between-batch variance to within-gel variance across three CS hydrogel compositions and four Dextran molecular weights. Values are reported as F-value : p-value.

|  | CS0 | CS50 | CS100 |
| --- | --- | --- | --- |
| 10 kDa | 0.01 ; 0.99 | 0.77 ; 0.52 | 0.45 ; 0.66 |
| 40 kDa | 1.15 ; 0.40 | 0.03 ; 0.97 | 0.46 ; 0.66 |
| 70 kDa | 0.33 ; 0.74 | 0.20 ; 0.83 | 0.25 ; 0.79 |
| 500 kDa | 0.52 ; 0.63 | 0.25 ; 0.79 | 0.14 ; 0.88 |

##### Methodological Conclusion

In summary, across both therapeutic proteins (HSA) and neutral polysaccharide probes (Dextran), variance in partition and diffusion coefficients is overwhelmingly dictated by spatial micro-heterogeneity within individual hydrogel pieces rather than preparation variability across independent batches. Because intra-gel spatial variance $s_{between}^2$ meets or even exceeds batch-to-batch variance $s_{between}^2$, sampling multiple positions within a single hydrogel specimen will give a reasonable estimate of the variance of the studied quantity (D or K) of the system without introducing pseudoreplication or underestimating experimental error. However, the analysis also revealed that batch-to-batch variability in collagen, inherent to its biological origin, can introduce measurable batch-to-batch differences in the hydrogel matrix. Notably, this effect was prominent only for polydisperse dextran probes possessing broad molecular weight distributions, whereas monodisperse HSA, characterized by a well-defined molecular weight, remained unaffected.
